# PERIODIC AND APERIODIC SPECTRAL SIGNATURES OF BEING MOVED BY ART

**DOI:** 10.64898/2026.06.18.732701

**Authors:** Deysha Poyser, Eugenio Rodríguez

## Abstract

Intense aesthetic experiences are among the most complex responses arising from the interaction of mind, brain, and context. Observations from fMRI suggest that when viewers feel highly moved by artworks, the underlying neural states differ from those accompanying less intense responses, particularly through recruitment of the DMN. Using electroencephalography and Bayesian category-specific cumulative link mixed models, we investigated whether such putative peak aesthetic responses exhibit threshold-specific neurodynamics rather than linear scaling with intensity. Twenty-two Chilean participants viewed 113 diverse local artworks whilst rating how moved they felt on a four-point scale. We analysed both canonical oscillatory power (θ–γ) and aperiodic components (offset and exponent) during the contemplation window and the post-elicitor window. Threshold-specific effects were found: spectral features differentiated the highest rating category from moderate responses, rather than scaling uniformly across all intensity levels. During artwork visualisation, power in the β_1_ and β_2_ bands, as well as the interaction of β_1_ with the 1/f exponent, predicted the transition to the most intense response; during the post-elicitor window, the aperiodic 1/f exponent predicted the transition from very low to higher-intensity responses. Modelling individual differences in spectral signatures (in the α and γ bands) credibly improved predictive performance (approximate leave-one-out cross-validation; elpd_loo), suggesting that neural variability reflects meaningful mechanistic heterogeneity in aesthetic processing rather than mere noise. These findings speak to a broader question, how the brain marks the intensity of conscious experience, and, more specifically, support the hypothesis that being intensely moved constitutes a qualitatively distinct neural state, characterised by specific configurations of oscillatory dynamics and cortical excitability that modulate the transition from low and moderate to peak engagement.

## 1. INTRODUCTION

Empirical aesthetics has largely emphasised the valence dimension of the aesthetic responses using the operational labels of preference, beauty, liking, pleasantness (Silvia & Brown, 2007; Chatterjee & Vartanian, 2014; Leder & Nadal, 2014; Redies, 2015; Pelowski et al., 2017; Wassiliwizky & Menninghaus, 2021) while the intensity dimension of aesthetic experience has remained comparatively underexplored. This is a significant gap considering the pervasiveness and relevance of intensity to experience in general (Kuppens et al., 2013), as it may be a key dimension to understanding how a mere perceptual encounter becomes an aesthetic, meaningful, self-relevant event.

Being moved is among the repertoire of intense or potent responses humans report (Menninghaus et al., 2015; Cova & Deonna, 2014; Zickfeld et al., 2019; Cullhed, 2020), characterised by mixed valence, personal significance, and absorption. Being moved applies to artworks, which, although created by others and not containing explicitly personal content, are capable of evoking archetypal responses as deeply personal as this. Interestingly, being moved by visual art has been elicited and measured in laboratory settings using self-report ratings of intensity as a proxy of the aesthetic response richness beyond like/not like (Vessel et al., 2012, 2019; Williams et al., 2018; Strijbosch et al., 2021), making it an empirically tractable model for studying how intensity shapes aesthetic experience.

Vessel and colleagues (2012, 2013) investigated hemodynamic correlates of aesthetic intensity whilst participants viewed diverse visual artworks and rated how moved they felt on a 1–4 scale. They identified two distinct patterns of brain activation: occipitotemporal and subcortical regions showed linear scaling with rating intensity, whilst regions of the default mode network (DMN), medial prefrontal cortex, posterior cingulate cortex, lateral temporal cortex, and temporoparietal junction, exhibited a threshold-like response, with strong deactivation for ratings 1–3 but marked attenuation (less deactivated) specifically at the highest rating. This selective response suggests that being highly moved involves a qualitatively distinct neural state rather than a progressive increase in engagement. Subsequent fMRI studies have confirmed the DMN involvement, and further showed that individual differences in aesthetic engagement are reflected in resting-state network architecture (Belfi et al., 2019; Vessel et al., 2019; Williams et al., 2018; Durkin et al., 2025).

Aesthetic responses vary substantially across individuals: the same artwork can move one person profoundly while leaving another indifferent (Jacobsen & Beudt, 2017; Vessel et al., 2018; Williams et al., 2018; Brielmann et al., 2023; Bignardi et al., 2024; Knoll et al., 2024). Rather than mere arbitrariness or noise to be minimised, this variability may encode the multiple ways in which the intensity of an aesthetic encounter can be realisable: from perceptual and affective capacities to cultural and biographical context, among other factors. Characterising individual differences in spectral dynamics, rather than averaging them away, may therefore reveal structural neural mechanisms underlying intense aesthetic experience and also shed light on the level at which these diverse factors exert their influence.

Whilst fMRI has established that intense aesthetic responses recruit a threshold-specific network configuration, it cannot resolve the temporal and spectral dynamics through which this state emerges and is sustained. EEG studies of aesthetic experience have reported modulations across α, beta, and γ bands (Strijbosch et al., 2021; for a review Welter & Lotte, 2024), but none have tested whether these dynamics exhibit threshold-specific rather than graded patterns, none have incorporated aperiodic spectral analysis, and none have modelled inter-individual differences in spectral sensitivity. These are not minor omissions: the aperiodic exponent indexes the cortical excitation/inhibition balance (Gao et al., 2017) and has been linked to state transitions and individual differences in cognitive flexibility (Voytek et al., 2015; Waschke et al., 2021), making it a theoretically motivated candidate for distinguishing qualitatively different neural states. The present study addresses this gap directly: using EEG recorded during graded aesthetic ratings of visual artworks, we tested whether periodic and aperiodic spectral features exhibit threshold-specific modulation at the transition to peak aesthetic responses, and whether these dynamics vary systematically across individuals. If spectral signatures show selective effects specifically at the highest intensity boundary rather than scaling uniformly, that constitutes evidence that being highly moved is a qualitatively distinct cortical state, not merely an extreme on a continuum. To this end, we recorded EEG while 22 participants rated Chilean visual artworks on a four-point scale of how moved they felt, and applied Bayesian category-specific cumulative link mixed models (CSCLMM) that allows us to model putative distances between categorical responses and separate signal variance into group and individual sources all while managing the uncertainty inherent in multilevel neural data.

## 2. MATERIALS AND METHODS

This study had two main objectives: (1) to characterise and model the aesthetic response of being moved in terms of both periodic and aperiodic components of the EEG signal; and (2) to test whether neural power dynamics exhibit a threshold-like pattern associated with the intensity of the aesthetic response.

To address these objectives, two spectral representations of the EEG signal were employed. First, power spectral density (PSD) estimates were used to characterise and model the structure and inter-individual variability of the periodic and aperiodic components of the signal, and to formally test the presence of threshold-like dynamics. As a complementary approach, a time-frequency decomposition was used to visualise and contrast power dynamics across rating categories over time and space via cluster-based permutation analyses.

Statistical modelling followed a progressive refinement logic motivated by three theoretical questions. First, do spectral features show threshold-specific effects at the population level? Second, do those effects persist after accounting for individual differences in spectral sensitivity? Third, do periodic and aperiodic components interact in their influence on aesthetic intensity, beyond their independent contributions? Each class of models addresses one question. Class 2 models inherit the credible predictors from class 1, while class 3 models build on interactions and integrate individual slopes as final refinement. Further details in §2.6.

Data, materials, and analysis code are indexed via Zenodo and GitHub, with access conditions noted below. Artwork image files are shared (Poyser, 2026b) in accordance with our agreement with MAVI-UC (Visual Art Museum, Santiago, Chile). The EEG and behavioural dataset (Poyser, 2026a), which includes demographics and ratings, is available upon reasonable request. Analysis scripts (Poyser, 2026c) are publicly available.

In the interest of transparency regarding AI tools in scientific production, we declare the use of Claude Sonnet 4.6 (Anthropic) solely to improve language after the manuscript was drafted, and to assist with the development, debugging, and annotation of analysis code. AI-assisted literature discovery tools (Consensus and Research Rabbit) were used to supplement the identification of relevant sources; all cited works were read and selected by the authors. All content and analyses remain the sole responsibility of the authors.

### 2.1. Participants

A total of 22 subjects (14 women, 8 men; mean age = 31.4 years, SD = 6.4) took part in the study. Inclusion criteria were: age 18 years or older, normal or corrected-to-normal vision, no current psychopharmacological treatment, and self-reported interest in visual art. Subjects were recruited through open calls targeting individuals with an interest in art, distributed via social media and physical posters placed near galleries and museums in Santiago, Chile. All subjects provided digital informed consent prior to participation, and received a compensation of an app equivalent of 5 EUR. One subject’s data was discarded because of an abnormal signal. All procedures were in accordance with the ethical standards of the Ethics Committee for Social Sciences, Arts, and Humanities of the Pontifical Catholic University of Chile and the Declaration of Helsinki.

No formal a priori power analysis was performed. Instead, a Bayesian estimation framework was adopted, in which inference rests on the full posterior distributions of model parameters, providing both effect estimates and their associated uncertainty, rather than on binary significance testing. The precision of the resulting estimates is reported via 95% credible intervals throughout. To assess whether the individual-difference structure was identifiable at the current sample size, we conducted a parameter recovery simulation in which synthetic datasets were generated from the posterior estimates of the fitted model and the same model was refitted to each; results are reported alongside the individual-differences discussion in §4.4 (see Figure 11).

### 2.2. Procedure

After reading and providing informed consent, subjects were prepared for the EEG recording session. Subjects were seated approximately 50 cm from a computer monitor and instructed to maintain visual fixation and minimise movement throughout the task. The experimental task was implemented in PsychoPy (Peirce et al., 2019). Each trial began with a fixation cross presented for 1 s, followed by the presentation of a single artwork image (one of 113) for 6 s. This was followed by a 3 s black screen, corresponding to a post-stimulus interval during which no motor response was required. An evaluative window with no time restriction then allowed subjects to rate their experience on the provided scale (Figure 1), followed by a variable inter-trial interval (2–3 s). After viewing thirty artworks each, the subjects were offered three opportunities to rest for as long as they needed. All subjects completed the full session using one or two of the available breaks.

**Figure 1.**
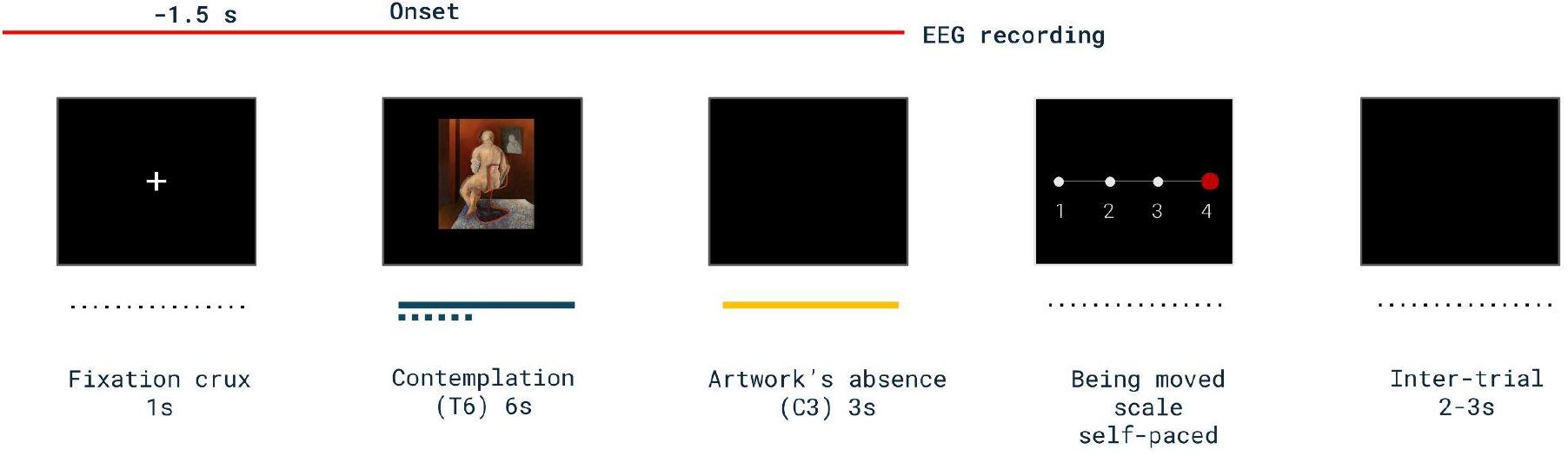
Experimental design. Each trial began with a fixation cross presented for 1 s, followed by the presentation of a single artwork for 6 s (contemplation window, T6). This was followed by a 3 s post-stimulus interval during which the artwork was absent (savouring window, C3). Participants then provided a self-paced rating of how much they felt moved by the artwork on a four-point scale (1–4), followed by a variable inter-trial interval (2–3 s).

Subjects were instructed to rate their immediate, intuitive response to each artwork based on how much they felt moved by it. Ratings were provided on a four-point scale ranging from 1 (not at all moved) to 4 (highly moved). The instruction was adapted from Vessel et al. (2012) and emphasises that artworks could range from beautiful to bizarre or ugly. Subjects were encouraged to base their ratings on whether the image felt powerful, profound, or resonant, rather than on aesthetic preference alone.

### 2.3. Artwork set

The visual artwork set (Poyser, 2026b), provided by MAVI-UC (Visual Art Museum, Santiago, Chile), comprises 113 digitised images selected and edited by the first author according to the following criteria. Only two-dimensional artworks were included, excluding photographs, and artists’ signatures were removed to minimise familiarity effects. All images were resized to fit within 800 × 800 pixels using bicubic interpolation, preserving original proportions and letterboxing with black borders against a black background screen. The set is diverse and consists of modern and contemporary Chilean artworks produced between 1920 and the present.

Artworks were presented in a fixed sequence across participants. This choice reflects the recognition that artworks are singular objects, not interchangeable tokens of a stimulus class, and that presentation order constitutes an interpretive context that shapes aesthetic experience (O’Doherty, 1986; Obrist, 2014). In predictive processing terms (Van De Cruys et al., 2024), what was just seen shapes the prediction landscape for the next encounter. We reasoned that randomisation would not neutralise order effects in this context, because each image introduces a distinct perceptual and semantic context; it would instead introduce multiple uncontrolled curatorial arrangements across participants. Fixing the sequence ensures that all participants traverse the same contextual trajectory, facilitating comparability of aggregated responses and reducing a source of between-subject variance that would otherwise be confounded with individual differences in aesthetic sensitivity. We acknowledge the trade-off: any systematic order effects are confounded with stimulus identity. However, the self-paced format allowed participants to regulate their own engagement and rest as needed. Also, the large number of trials limits the influence of any single position, and the mixed-model framework with random intercepts for artworks partially absorbs stimulus-specific variance. In addition to this, to assess whether order patterns threaten the spectral findings, trial position was added as a covariate to test its relevance as confound. No credible effect was found nor credible impact on results (see details in §4.6).

### 2.4. Signal acquisition and preprocessing

EEG data were acquired using a BioSemi ActiveTwo system with 64 scalp electrodes arranged according to the international 10–20 system. Electrode offsets were kept below 80 kΩ. Signals were amplified and recorded at a sampling rate of 2048 Hz. EEG data were preprocessed in MATLAB. Continuous recordings were downsampled to 512 Hz, band-pass filtered between 0.3 and 45 Hz, and initially referenced to the mastoids before being re-referenced to the common average. Data were epoched from −1.5 s to 10.5 s relative to artwork onset. Baseline correction was applied using the −1.5 to 0 s pre-stimulus interval. Segments containing excessive noise and bad channels were identified by visual inspection, removed, and subsequently interpolated. Independent component analysis (ICA) was then performed to identify stereotypical non-neural components (e.g. ocular and muscular artefacts), which were removed prior to further analysis. From each trial epoch, two temporal windows were extracted. The first corresponded to artwork contemplation (0–6 s; hereafter referred to as T6), during which the artwork was visible. The second corresponded to a post-elicitor, savouring window (6–9 s; hereafter referred to as C3), during which the artwork was no longer present and no explicit task was required, as the evaluative phase followed immediately afterwards. For computational economy, electrodes were assigned to 26 predefined ROIs based on anatomical location and hemispheric laterality (spanning frontopolar/prefrontal, frontal, fronto-central, central, centro-parietal, parietal, parieto-occipital, occipital, and temporal groups, with left, midline, and right subdivisions) using a fixed channel-to-ROI mapping (Table 1A).

**Table 1.** 12 Bayesian CSCLMMs were planned following complexity progression, allowing for the identification of population and individual level influences. “#” id number of each model, “class models” describes one of the three layers of analysis, “window” specifies the window data analysed. For the R syntax, refer to Table 4A.

| ID model | Model class | Window/Data |
| --- | --- | --- |
| 1 | Class 1/ Periodic components, population level | T6 |
| 2 | Class 1/ Periodic components, population level | C3 |
| 3 | Class 1/ Aperiodic components, population level | T6 |
| 4 | Class 1/ Aperiodic components, population level | C3 |
| 5 | Class 2/ Periodic components, adjusted by individual differences | T6 |
| 6 | Class 2/ Periodic components, adjusted by individual differences | C3 |
| 7 | Class 2/ Aperiodic components, adjusted by individual differences | T6 |
| 8 | Class 2/ Aperiodic components, adjusted by individual differences | C3 |
| 9 | Class 3/ Aperiodic & periodic interaction, population level | T6 |
| 10 | Class 3/ Aperiodic & periodic interaction, population level | C3 |
| 11 | Class 3/ Aperiodic & periodic interaction,adjusted by individual differences | T6 |
| 12 | Class 3/ Aperiodic & periodic interaction,adjusted by individual differences | C3 |

### 2.5. Spectral Analysis

#### 2.5.1 PSD and time-frequency representations

Power spectral densities were computed because they provide noise-reduced stationary spectral estimates that are well suited for spectral parameterisation, in contrast to time-resolved wavelet-based power estimates. PSDs were estimated using Welch’s method for each subject, trial, and region of interest (ROI) over the 2–45 Hz frequency range, using a 2000 ms sliding window with 1000 ms overlap. The FFT length was set to the next power of two of the window length. Aperiodic and periodic spectral features were extracted in Python (v3.11.13) using the FOOOF algorithm (Donoghue et al., 2020) in fixed aperiodic mode, fitted over the 2–45 Hz range. This procedure yielded estimates of the aperiodic offset and exponent, goodness-of-fit (R^2^), and parameters of spectral peaks rising above the aperiodic component, including centre frequency, power, and bandwidth, which were used to characterise the oscillatory content. To facilitate further interpretation, the continuous output of the FOOOF was sorted into fixed bins, i.e. the canonical frequency bands: θ, 4-8 Hz; α, 8-12 Hz; β_1_, 12-16 Hz; β_2_, 16-20 Hz; beta-3, 20-30 Hz; γ, 30-45 Hz.

Time-frequency decomposition was performed using Morlet wavelets implemented in the FieldTrip toolbox (Oostenveld et al., 2011). Single-trial power estimates were computed over frequencies from 1 to 45 Hz in 1-Hz steps. The number of wavelet cycles increased linearly with frequency, ranging from 3 cycles at 1 Hz to 10 cycles at 45 Hz, balancing temporal resolution at lower frequencies with spectral resolution at higher frequencies. Power estimates were computed at all available time points and zero-padded to the next power of two to optimise spectral estimation.

#### 2.5.2 Descriptive analysis

To describe the periodic and aperiodic components, we rely on FOOOF outputs (as described above) associated with ratings categories and the T6 and C3 windows. We described the dominant features, tendencies and variability across subjects. To describe categorical differences in the dynamics of power over time, we rely on the TF charts to build contrasts between category responses over time and further complement it with a spatially informed inference via cluster-based permutation analyses. For this purpose, power was averaged within consecutive 500 ms time bins and within canonical frequency bands.

Permutation statistics were performed using FieldTrip’s non-parametric Monte Carlo framework with a dependent-samples t-statistic, applied to subject-level time-frequency representations. In order to align the sensor-level analyses with the ordinal structure identified by the cumulative link mixed models, contrasts were defined to mirror the model cutpoints rather than single rating categories. Specifically, two contrasts were tested: (i) rating 1 versus ratings 2–4 (corresponding to the 1|2 transition), and (ii) ratings 1–3 versus rating 4 (corresponding to the 3|4 transition). Within each subject, power spectra belonging to the same contrast were pooled at the trial level, yielding trial-weighted condition averages.

Cluster formation was defined across neighbouring electrodes using a distance-based criterion (40 mm). Cluster-level inference relied on the maximum cluster-level sum of t-values as the test statistic. A two-sided test was applied with a cluster-forming threshold of α = 0.05 and a cluster-level significance threshold of α = 0.05, based on 2,000 random permutations. Complete statistical output is provided in the Annex (Table 3A).

**Table 2.** Bayesian R^2^ (conditional and marginal) for all fitted models. Conditional R^2^ includes both fixed and random effects; marginal R^2^ reflects fixed effects only. Posterior medians and 95% credible intervals are reported. Pop. = population; ind. = individual; exp. = exponent.

| ID | Model | R<sup>2</sup><br>conditional | R<sup>2</sup>c 95% CI | R<sup>2</sup><br>marginal | R<sup>2</sup>m 95% CI |
| --- | --- | --- | --- | --- | --- |
| Contemplation window (T6) |  |  |  |  |  |
| 1 | Periodic, pop. level | 0.25 | [0.23, 0.28] | 0.04 | [.02, .07] |
| 3 | Aperiodic, pop. level | 0.25 | [0.22, 0.27] | 0.004 | [< .001, .01] |
| 5 | Periodic + ind. slopes | 0.28 | [0.25, 0.30] | 0.05 | [.01, .11] |
| 7 | Aperiodic exp. + ind. slopes | 0.25 | [0.22, 0.27] | 0.005 | [< .001, .02] |
| 9 | Interaction, pop. level | 0.25 | [0.23, 0.28] | 0.03 | [.008, .05] |
| 11 | Interaction, pop + slopes | 0.28 | [0.26, 0.31] | 0.075 | [.025, 0.13] |
| Post-elicitor window (C3) |  |  |  |  |  |
| 2 | Periodic, pop. level | 0.23 | [0.21, 0.26] | 0.01 | [.003, .02] |
| 4 | Periodic + ind.<br>slopes | 0.25 | [0.22, 0.27] | 0.009 | [.001, .03] |
| 6 | Aperiodic, pop.<br>level | 0.25 | [0.22, 0.27] | 0.02 | [.003, .03] |
| 8.a | Aperiodic + ind.<br>slopes | 0.26 | [0.23, 0.28] | 0.027 | [.002, .07] |
| 8.b | Exponent only +<br>ind. slopes | 0.26 | [0.23, 0.28] | 0.026 | [.001, .07] |
| 10 | Interaction, pop<br>level | 0.25 | [0.23, 0.27] | 0.019 | [0.006, 0.04] |

**Table 3.** Expected predictive accuracy of models (for T6) and their comparison. Via Leave-One-Out Cross-Validation “loo_compare()”, the expected log pointwise predictive density (elpd) was estimated and compared to the model with the largest ELPD (the model in the first row, elpd= −3056.3) by making pairwise comparisons between each model against the best first row.

|  |  |  |
| --- | --- | --- |
| T6 |  |  |
| Model | elpd_diff | se_diff |
| 11 / Interactionist with ind. slopes | 0.0 | 0.0 |
| 5 / Periodic + slopes | -2.5 | 3.9 |
| 9 / Interactionist pop lvl | -30.0 | 8.3 |
| 1 / Periodic pop lvl | -31.1 | 8.7 |
| 7 / Aperiodic with ind. slopes | -34.8 | 9.6 |
| 3 / Aperiodic pop lvl | -38.0 | 9.9 |

### 2.6. Statistical Modelling

A Bayesian approach is advantageous for restricted neural data such as ours, and more generally, it supports the integration of prior knowledge and future replication through the explicit specification of priors (Körding, 2014). To ensure the quality, transparency, and reproducibility of the Bayesian analyses, we followed the Bayesian analysis reporting guidelines (BARG) as presented by Kruschke (2021) which, in addition to documenting the parameters and outputs, suggests formalising the models (§2.6.3). All analyses were conducted in R (version 4.4.3) using the brms (2.22.0), cmdstanr (0.8.0), Stan (version 2.32.7) and tidyverse (2.0.0) packages.

All models were compared within each temporal window using leave-one-out cross-validation LOO-CV (Vehtari et al., 2017) and Bayesian R^2^ (Gelman et al., 2019). LOO quantifies out-of-sample predictive accuracy and penalises complexity that does not improve generalisation. Following Sivula et al. (2025), models with an absolute elpd (expected log pointwise predictive density) difference below 4 are considered practically indistinguishable in predictive accuracy; for larger differences, the elpd_diff and its standard error are reported to allow readers to assess magnitude and uncertainty. Bayesian R^2^ decomposes explained variance into a conditional component (fixed + random effects) and a marginal component (fixed effects only).

#### 2.6.1. Hypothesis testing via Category-Specific Cumulative Link Mixed Models (CSCLMM)

To test whether oscillatory dynamics predicted the intensity of the aesthetic response in a thresholded or step-like manner, we employed Category-Specific Cumulative Link Mixed Models (CSCLMMs). These models extend standard ordinal regression by allowing the effects of the predictors to vary across response thresholds (cutpoints = τ). This makes them particularly well-suited to testing hypotheses about non-linear or threshold-specific effects in ordinal outcomes, as is the case here.

#### 2.6.2. Data structure and predictors

Spectral features were derived from trial-level EEG data using FOOOF parametrisation methods. For periodic activity, oscillatory peak power was extracted and grouped into canonical frequency bands (delta to γ). Band-specific power estimates were aggregated at the trial level and used as continuous predictors reflecting the relative strength of rhythmic activity in each band. For aperiodic activity, the exponent and offset of the aperiodic (1/f-like) component of the power spectrum were estimated per trial, providing broadband descriptors of spectral shape. In both cases, predictors were modelled at the trial level with ROI collapsed by averaging across all 26 ROIs within each subject, and the ordinal aesthetic rating (1–4) as the dependent variable.

#### 2.6.3 Models specification

Let *y*_*i*_ ∈ {1, 2, 3, 4} denote the rating for trial *i*. Ratings were analysed using CSCLMM with a logit link, implemented in brms. These models estimate cumulative probabilities

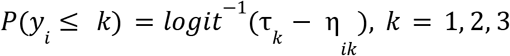

where *k* indexes the ordered response thresholds (cutpoints 1|2, 2|3, and 3|4), τ_*k*_ denotes the threshold-specific intercepts, and η_*ik*_ the category-specific linear predictor. The observed ordinal response is assumed to arise from an unobserved continuous variable 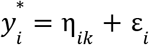 where ε_*i*_ follows a logistic standard distribution. Because η_*ik*_ is indexed by threshold *k*, then predictors are allowed to differentially modulate the probability of crossing each response boundary.

Three model classes were estimated for the contemplation (T6) and the post-elicitor (C3) windows. On top of the theoretical assumptions (§1), model construction followed a progressive refinement logic based on parsimony and computational resources. A total of 12 models were specified (Table 1), with the Class 3 interactionist specification representing the most complete theoretical implementation of the threshold hypothesis. Figures 2 and 3 schematise this logic for the T6 and C3 windows, respectively.

**Figure 2.**
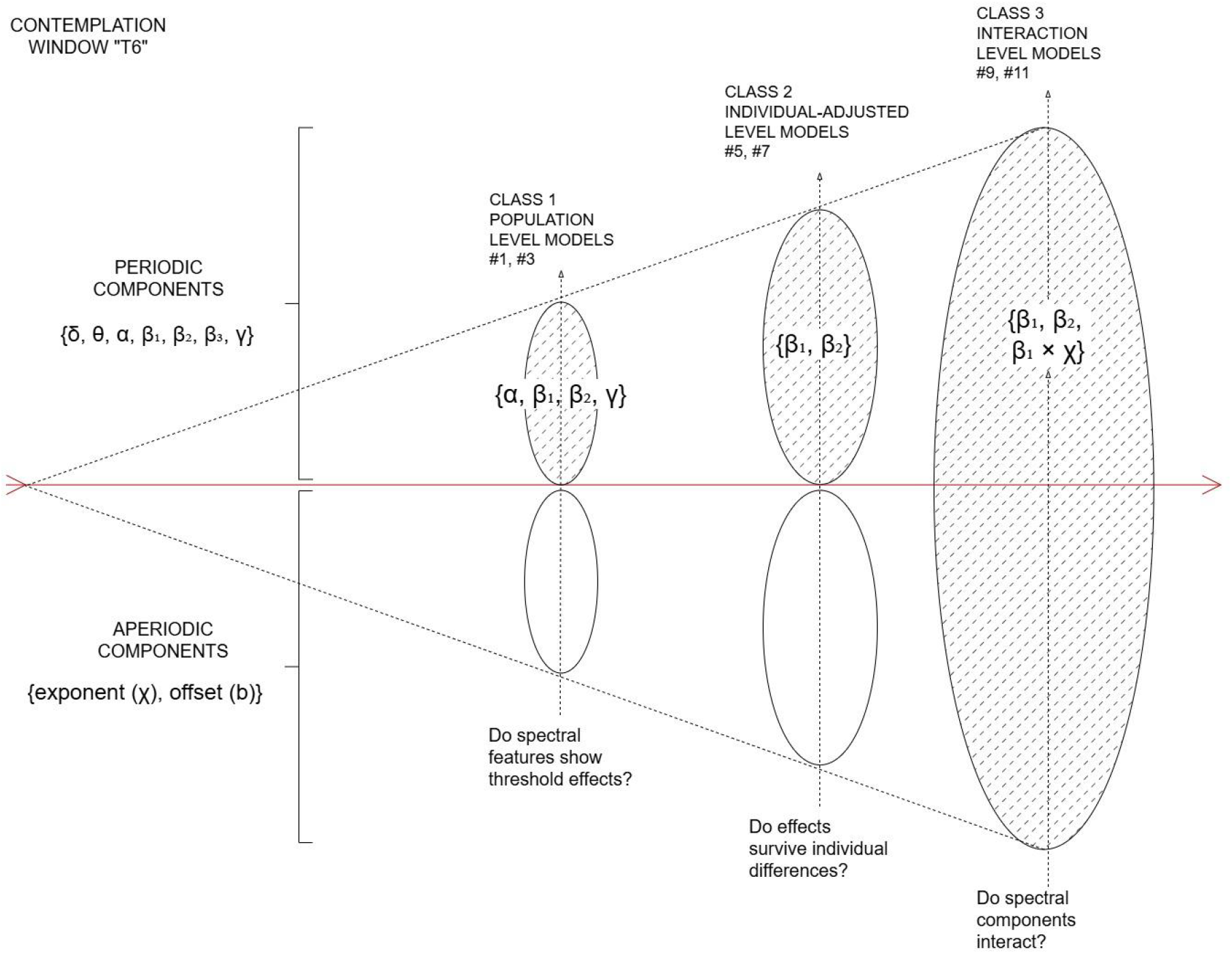
Progressive modelling logic and credible predictors for the contemplation (T6) window. Starting from the full periodic and aperiodic predictor set (right), three model classes address successive theoretical questions (bottom labels). Class 2 inherits the credible predictors from the previous one (red arrow), with ellipse size reflecting increasing model complexity. Predictor sets within each ellipse show the spectral features that survived at each level. Note that α and γ, credible at Class 1, disappear at Class 2, having been absorbed by individual-specific random slopes. At Class 3, periodic and aperiodic tracks merge into a single interactionist specification; yielding the best fitting model by LOO-CV and the basis for interpretation throughout.

**Figure 3.**
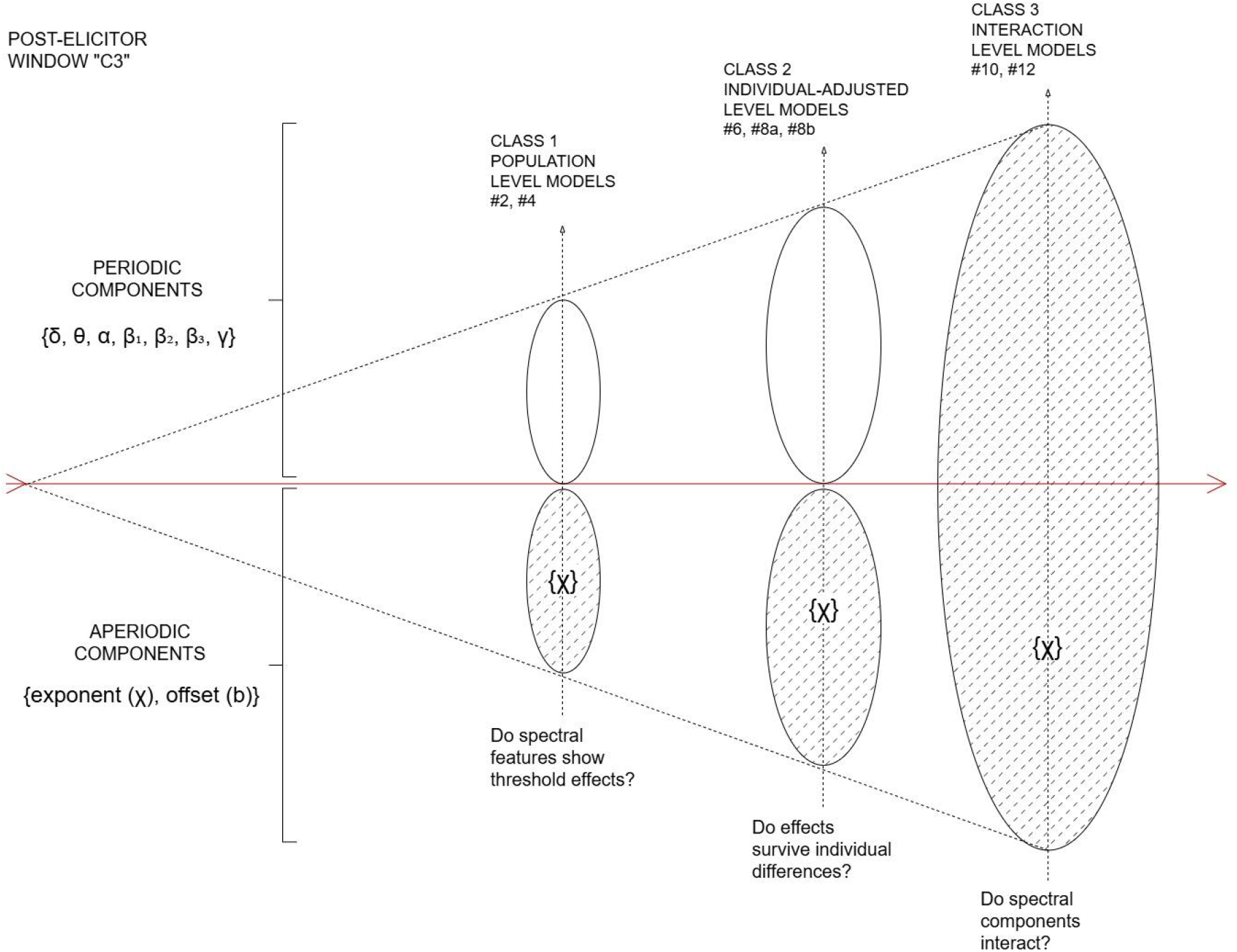
Progressive modelling logic and credible predictors for the post-elicitor (C3) windows. Starting from the full periodic and aperiodic predictor set (right), three model classes address successive theoretical questions (bottom labels). Each class inherits only the credible predictors from the previous one (red arrow), with ellipse size reflecting increasing model complexity. Predictor sets within each ellipse show the spectral features that survived at each level. At Class 3, periodic and aperiodic tracks merge into a single interactionist specification; yielding the best fitting model by LOO-CV and the basis for interpretation throughout. Note that only the aperiodic exponent exhibited credible effects.

## I. Class 1 model. Population–level effects

The first class model tested population-level effects of either periodic oscillatory power across canonical frequency bands, with *f* ∈ {δ, θ, α, β_1_, β_2_, β_3_, γ) or aperiodic components exponent χ, and offset *b*, independently. Both sub-models included threshold-specific fixed effects and random intercepts for subject and artwork only:

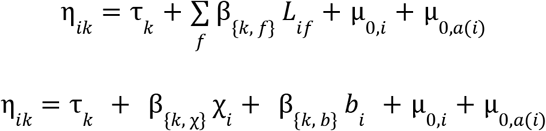

where β_{*k, f*}_ is the threshold-specific population coefficient for band *f* and *L*_*if*_ is the z-scores peak power for *f* at trial _*i*_, χ_*i*_ and *b*_*i*_ are the z-scored aperiodic exponent and offset for trial *i*, and µ_0,*i*_*~ N*(0, σ^2^) and µ_0,*a*(*i*)_ *~ N*(0, σ^2^) are subject and artwork random intercepts, respectively. Models 1-4 (Table 1 and 4A) are the periodic and aperiodic implementations.

**Table 4.** Expected predictive accuracy of models (for C3) and their comparison. Via Leave-One-Out Cross-Validation “loo_compare()”, the expected log pointwise predictive density (elpd) was estimated and compared to the model with the largest ELPD (the model in the first row) by making pairwise comparisons between each model against the best first row.

|  |  |  |
| --- | --- | --- |
| C3 |  |  |
| Model | elpd_diff | se_diff |
| 8.b/ Aperiodic exponent only + ap.<br>exponent slopes | 0.0 | 0.0 |
| 8.a/ Aperiodic components +<br>exponent slopes | -2.5 | 1.2 |
| 10/ Interactionist | -6.7 | 7.2 |
| 4/ Aperiodic pop level | -11.4 | 4.6 |
| 6/ Periodic + slopes | -31.0 | 12.8 |
| 2/ Periodic pop level | -49.3 | 12.2 |

## II. Class 2 model. Individual-difference-adjusted effects

Class 2 includes random slopes only for predictors that show credible Class 1 effects: α, β_1_, β_2_, γ for periodic models; χ for the aperiodic model. In addition to random intercepts, random slopes for these spectral predictors were included at the subject level as follows:

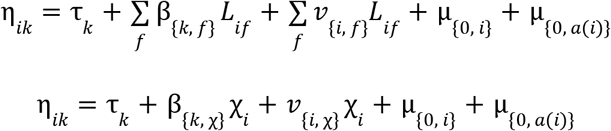

where *f* ∈ {α, β_1_, β_2_, γ} for the periodic sub-model, 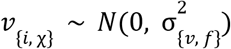 is the subject-specific random slope for component *f*, and 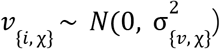 is the subject-specific random slope for the aperiodic exponent. Random slopes are shared across thresholds (e.g., not category-specific) and specified without cross-random-effect correlations to improve sampling stability, as category-specific effects in cumulative models are currently considered experimental in `brms`. Models 5-8 (Table 1 and 4A) are the periodic and aperiodic implementations in R.

## III. Class 3 models. Cross-domain interactions

The third class model examines interactions between periodic and aperiodic effects. Two sub-models were specified differing in their random-effects structure: population-level interactions and those interactions with individual slopes for α and γ power bands.

Models 9 and 10 (Table 1 and 4A) test threshold-specific main effects and interactions between β_1_, β_2_ and χ, with random intercepts only:

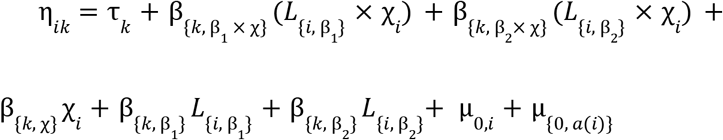

Model 11 is an extension that adds (a) non-category-specific fixed effects for α and γ power, which serve as adjustment covariates with a single coefficient shared across thresholds; and (b) subject-level random slopes for all five spectral predictors (α, β_1_, β_2_, γ And χ), each specified via separate grouping terms to avoid estimating cross-correlations. Note that β_α_ and β_γ_ are constant across thresholds, reflecting their role as adjustment covariates rather than primary predictors. Again layer credibility criteria was applied; this last model was fitted for the T6 window only, as periodic effects did not reach credibility in window C3 under any Class 1 or Class 2 specification, removing the theoretical motivation for a cross-domain interaction model in that window.

## IV. Priors and estimation

Weakly informative priors were assigned to regularise estimation while remaining compatible with plausible effect sizes on the logit scale. Population-level coefficients (both category-specific and non-category-specific) received *N*(0, 1) priors. On the logit scale, a coefficient of ±1 corresponds approximately to a 2.7 fold change in cumulative odds per standard deviation of the predictor; effects larger than ±2 would imply implausibly large shifts in response probabilities given the within-subject design. The *N*(0, 1) prior therefore assigns most mass to realistic effect sizes while remaining only weakly informative. Standard deviations of subject- and artwork-level random effects received half-normal priors (SD = 1).

Threshold intercepts were given ordered normal priors centred at −1.5, 0, and 1.5, reflecting the approximately balanced ordinal scale.

All models were estimated using Hamiltonian Monte Carlo via brms with the cmdstanr backend (four chains, overdispersed initial values). Convergence was assessed via 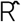 (≤ 1.01), bulk and tail effective sample sizes, trace plots, and pairwise posterior scatter plots. Prior predictive checks (Figure 6A) confirmed that the chosen priors produced simulated response distributions within the plausible range. Posterior predictive checks (Figure 7A) were performed globally and per response category. Model comparison used approximate leave-one-out cross-validation (LOO-CV, Table 3 and 4) with Pareto-k diagnostics (all k < 0.7). Model fit is summarised via Bayesian R^2^ (conditional and marginal, Table 2). Convergence diagnostics and visualisations for cases with divergences, and full model summaries are also reported in the Annex (see §***Note on Table 5A and Figures 8A–12A*)**.

## 3. RESULTS

### 3.1. Individual variability among spectral components

Subjects exhibited stable individual differences in aperiodic EEG parameters across both temporal windows (Figure 1A). The aperiodic 1/f exponent and offset showed distinct subject-level profiles, with high within-subject reliability (exponent: ICC(3,1) = 0.96, [0.91, 0.98]; offset: ICC(3,1) = 0.94, [0.89, 0.98]), indicating that inter-individual variability substantially exceeded window-related variance. This reliability persisted when retaining spatial and trial-level variability: exponent ICC(3,1) = 0.97, [0.93, 0.99]; offset ICC(3,1) = 0.95, [0.88, 0.98].

Subjects also showed distinct oscillatory profiles across rating levels (Figure 2A), with particularly heterogeneous patterns in the α band. Reliability of periodic parameters was computed on band-assigned FOOOF peak parameters across windows (T6 and C3, Table 2A) and varied by frequency band. Peak amplitude showed high cross-window stability across all bands (ICC range ≈ 0.83–0.95), indicating that individual differences in oscillatory strength were preserved despite task-related modulation. Peak centre frequency showed lower reliability, particularly for α (ICC = 0.66) and β_2_ (ICC = 0.59), where individual differences accounted for ~60% of the variance. Reliability increased at higher frequencies (β_3_ ICC = 0.91; γ ICC = 0.97). Peak bandwidth showed intermediate stability (ICC ≈ 0.78–0.89). Oscillatory power thus reflects a more stable signal, while dominant frequency features show greater rating and trial-level variability.

### 3.2. Association of periodic components with rating response during the contemplation window (T6)

By only modelling shared effects (Class 1 models), oscillatory power in canonical frequency bands showed selective associations with aesthetic rating thresholds during contemplation (T6), with no credible effects during the post-elicitor window (C3).

Most frequency bands showed small and uncertain associations with threshold probabilities during T6. However, α, β_1_, β_2_, and γ power each modulated specific cutpoints (Figure 3A). Increased α power was associated with a reduced probability of crossing the lowest cutpoint (1|2: β = −0.24, [−0.44, −0.03]). β_1_ reduced the likelihood of higher ratings at both the lowest (1|2: β = −0.29, [−0.51, −0.07]) and highest (3|4: β = −0.29, [−0.53, −0.05]) thresholds, while β_2_ affected only the highest threshold (3|4: β = −0.28, [−0.48, −0.08]). Uniquely among the bands tested, γ was the band with a positive association (3|4: β = 0.24, [0.02, 0.45]), indicating that increased γ power raises the probability of the highest rating.

When individual spectral sensitivities were included (Class 2, Figure 2), α and γ effects were fully absorbed by subject-specific random slopes in the contemplation window (T6). Credible and selective effects emerged exclusively within the β band (Figure 4). β_1_ power exhibited influence at both the lowest (1|2: β = −0.31, [−0.58, −0.05]) and highest (3|4: β = −0.32, [−0.60, −0.05]) cutpoints and β_2_ maintained its selective negative association at the highest cutpoint only (3|4: β = −0.28 [−0.53, −0.01]). This model (5, Table 1) explained 28% of rating variance (conditional R^2^ = 0.28, [0.25, 0.30]; marginal R^2^ = 0.05, [0.01, 0.11]), with the majority attributable to random rather than fixed effects. The latent space was well defined and asymmetric: the highest rating threshold (3|4) was positioned farther from the scale centre (2.07) than the lower thresholds (2|3 = 0.18; 1|2 = −1.41).

**Figure 4.**
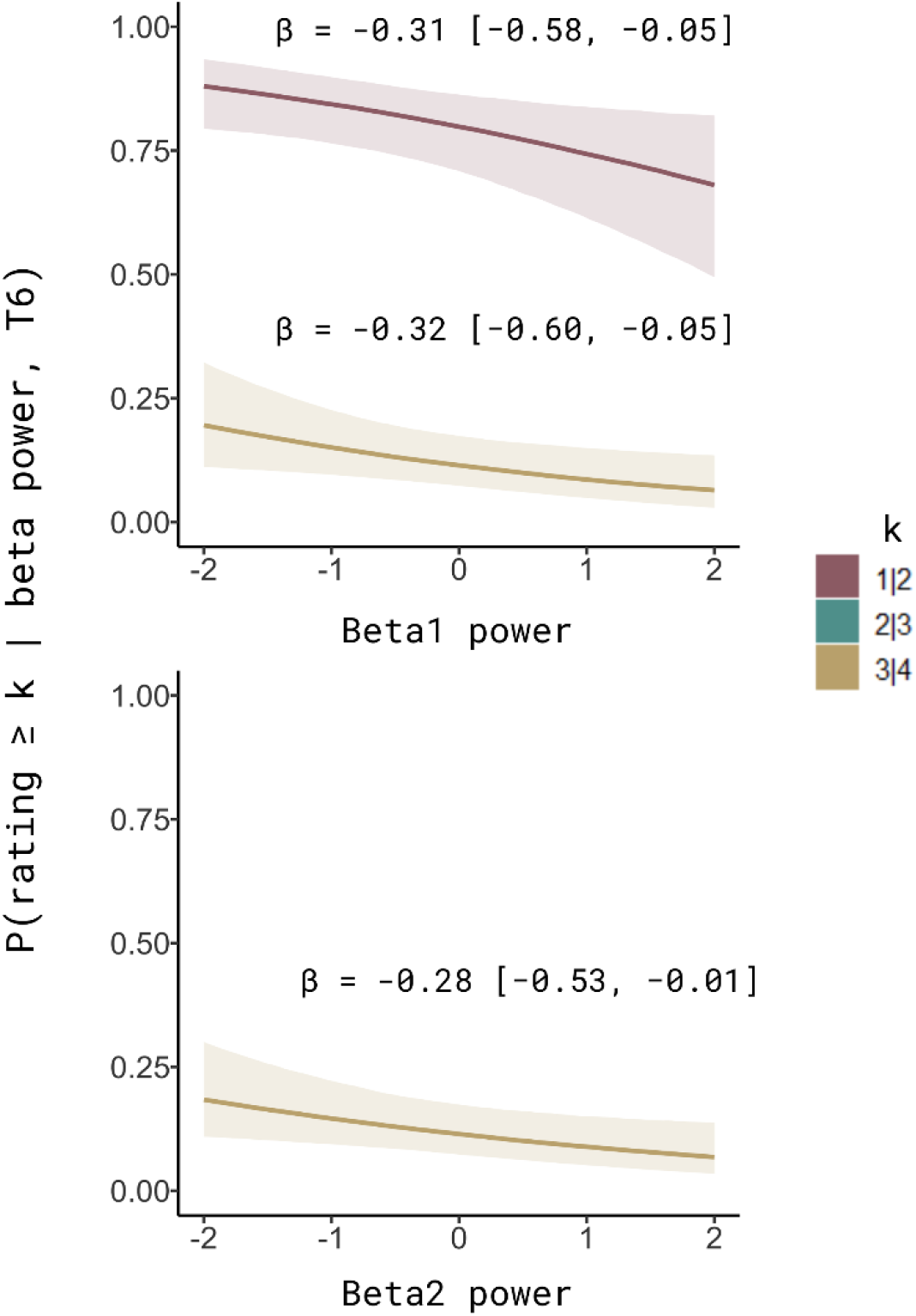
Individual-difference-adjusted conditional effects of oscillatory band power during contemplation (T6). Posterior estimates show how z-scored β-band power (β_1_, β2) credibly modulates the probability of crossing selected cutpoints on the latent aesthetic response scale. Shaded areas indicate 95% credible intervals. Estimates are derived from a CSCLMM fitted to ratings (r_1_–r_4_), including band-specific power predictors at T6 and random effects accounting for inter-individual differences in both baseline rating propensity and baseline band power.

Inter-individual variability was predominantly captured by α (SD = 0.61), indicating substantial heterogeneity in how α power relates to aesthetic judgments across subjects. In contrast, β_1_, β_2_, and γ showed more moderate variance (SDs ≈ 0.26, 0.29, 0.29, respectively). This pattern indicates that what appeared at the population level as α and γ effects were not shared across participants but instead reflected systematic between-subject heterogeneity; in other words, different individuals showing α and γ power displaced the latent aesthetic response in opposing directions.

### 3.3. Timing and scalp distribution of β-band effects on rating transitions

To characterise the timing and scalp distribution of the β effects identified so far, we conducted cluster-based permutation tests on time-resolved β_1_ and β_2_ power over consecutive 500 ms bins during the T6 window (Figure 5).

**Figure 5.**
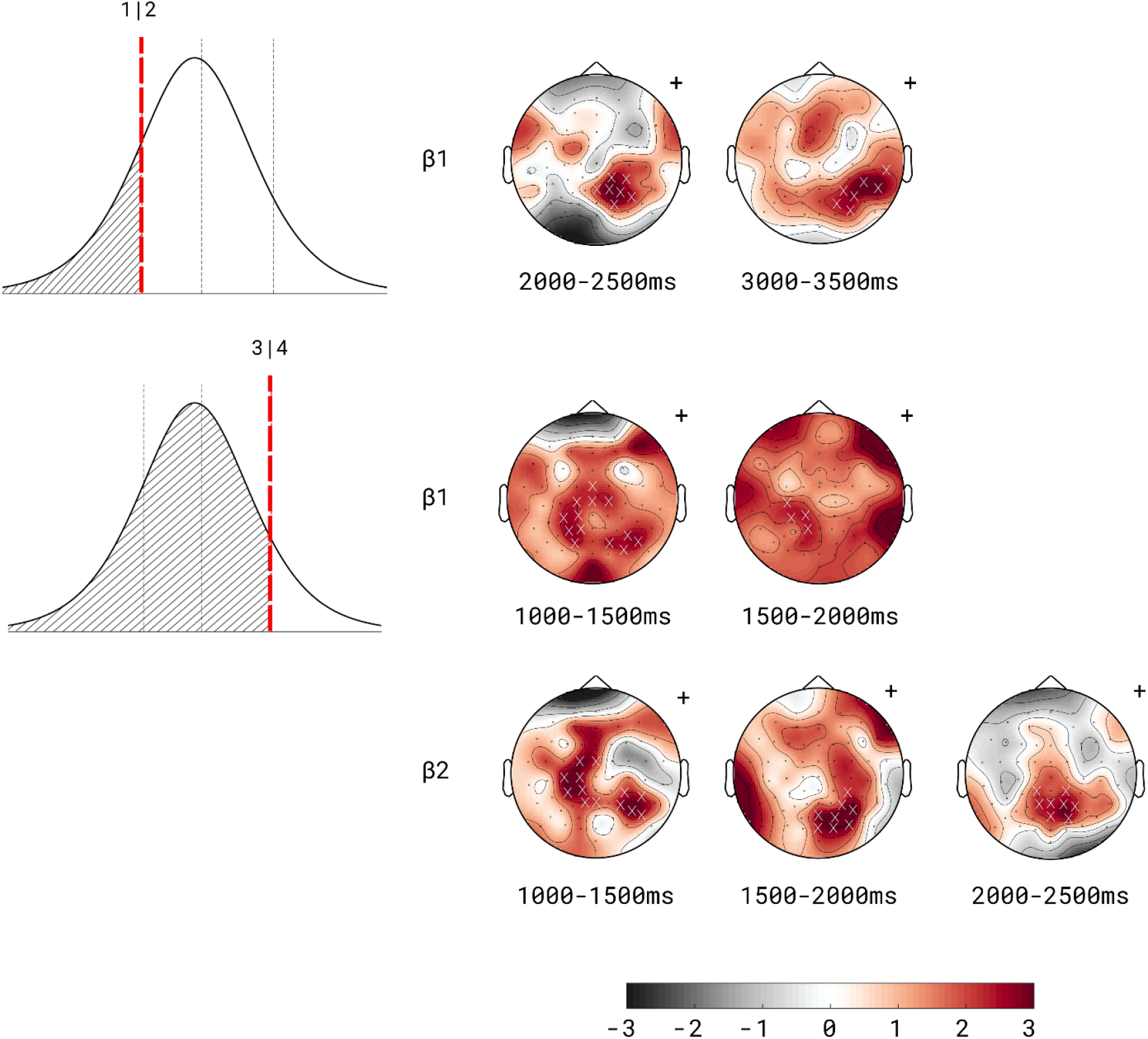
Spatio-temporal contrasts of β_1_ and β2 power during the T6 window, evaluated with cluster-based permutation tests aligned with ordinal model thresholds. Contrasts correspond to the 1|2 cutpoint (log-odds of transitioning from rating 1 to higher ratings) and the 3|4 cutpoint (log-odds of entering the highest rating category). Scalp maps show differences (red: positive, black: negative, all differences were positive) in band power within 500 ms time bins; white crosses indicate electrodes belonging to significant clusters. Cluster statistics reflect consistent threshold-related differences across subjects, not maximal absolute power.

Only the β_1_ clusters distinguished the 1|2 boundary (1 vs. all contrast). β_1_ cluster effects emerged later, during mid-to-late contemplation (2000–2500 ms: p = 0.012, cluster mass = 18.5, 7 channels; 3000–3500 ms: p = 0.015, cluster mass = 18.6, 7 channels). The first cluster peaked at electrode P2 over the centro-parietal regions, and the second peaked at CP6, biased to the right posterior area and forming a patch over the temporoparietal regions.

At the 3|4 threshold (all vs. 4 contrast), significant clusters emerged early in the contemplation window for both β_1_ and β_2_. β_1_ showed significant clusters from 1000–2000 ms over centro-parietal regions (1000–1500 ms: p = 0.018, cluster mass = 21.2, 9 channels; 1500–2000 ms: p = 0.044, cluster mass = 9.3, 4 channels), and β_2_ from 1000–2500 ms over parieto-occipital regions (1000–1500 ms: p = 0.004, cluster mass = 21.2, 8 channels; 1500–2000 ms: p = 0.005, cluster mass = 23.2, 8 channels; 2000–2500 ms: p = 0.013, cluster mass = 19.3, 8 channels). In the earliest bin (1000–1500 ms), the two bands showed a very similar pattern over centro-parieto-occipital regions, with β_1_ peaking at P3 and P8 and β_2_ at electrode C3 and P6. From 1500–2000 ms onward, a sub-band topographic differentiation emerged: a left-lateralised β_1_ cluster peaking at CP1 distinguished lower ratings from the highest, while β_2_ was right-lateralised, peaking at channel P6, with this parieto-occipital cluster extending to 2500 ms and peaking at P2. Across all contrasts, β power was higher for lower-rated trials.

### 3.4. Association of aperiodic components with rating response during post-elicitor window

The aperiodic 1/f exponent showed threshold-specific associations with aesthetic ratings exclusively during the post-elicitor window (C3); no credible effects were observed during artwork contemplation (T6). At the population level (Class 1), higher exponent values in C3 reduced the probability of crossing the 1|2 (β = −0.43, [−0.71, −0.15]) and 2|3 (β = −0.29, [−0.55, −0.05]) boundaries; the effect at 3|4 was in the same direction but uncertain. The aperiodic offset showed no credible effects in either window. As with the periodic modelling, threshold intercepts were well-separated and asymmetric towards the 3|4 boundary, confirming adequate ordinal resolution of the response scale.

When individual spectral sensitivities were included (Class 2), the aperiodic exponent showed substantial inter-individual variability in C3 (SD = 0.49), consistent with the stable individual profiles described in §3.1; again no credible effects emerged in T6. In C3, retaining the aperiodic offset as a covariate (Model 8.a) showed that the exponent effect remained credible only at the 1|2 threshold (β = −0.42, [−0.77, −0.08]), while the offset itself still showed no credible effect at any threshold. Dropping the offset (Model 8.b) sharpened the estimates: the exponent now showed credible effects across all three thresholds (1|2: β = −0.38, [−0.66, −0.12]; 2|3: β = −0.27, [−0.53, −0.01]; 3|4: β = −0.29, [−0.57, −0.02]), with the 1|2 effect consistently the strongest across both specifications.

The 1/f aperiodic exponent thus showed robust, threshold-specific effects in C3 that persisted after accounting for individual spectral differences (Figure 6). Substantial variability in baseline rating propensity remained at both the subject level (SD = 0.92, [0.67, 1.26]) and artwork level (SD = 0.63, [0.52, 0.76]). At the population level, aperiodic 1/f exponent accounted for roughly 3% of the variance in aesthetic intensity responses, and ~26% was explained when individual spectral differences were included (marginal R^2^ = 0.027, [0.001, 0.07]; conditional R^2^ = 0.26, [0.23, 0.28]).

**Figure 6.**
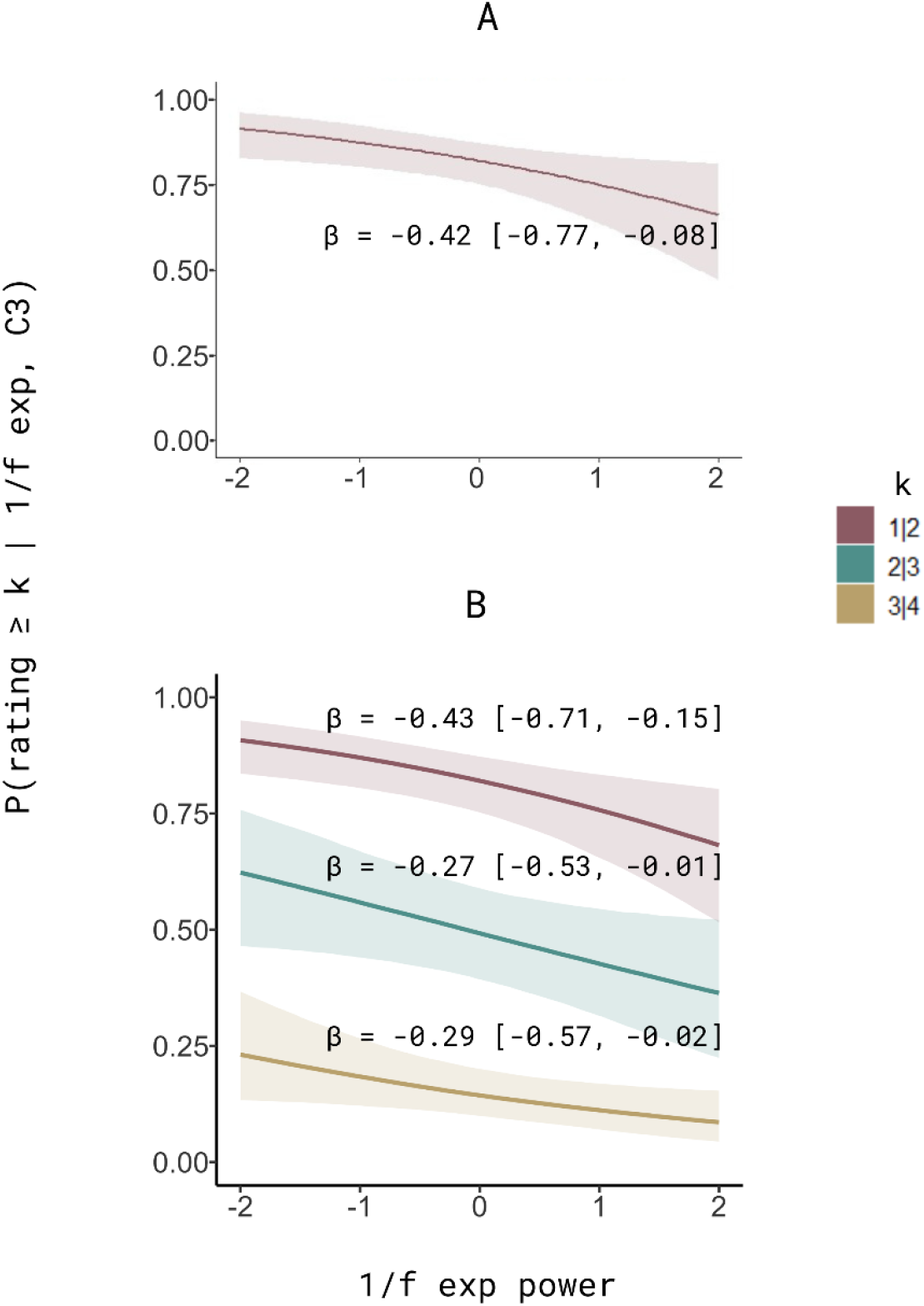
Individual-difference-adjusted conditional effects of 1/f exponent during post-elicitor (C3) based on Model 8.a (panel A) and 8.b (panel B) (model’s details in Table 1 and 4A). Model 8.a maintained the aperiodic offset as a fixed variable to corroborate the stability of the aperiodic exponent effect. Note that without the aperiodic offset (Model 8.b), the effect of the 1/f exponent influenced all thresholds. Shaded areas indicate 95% credible intervals.

### 3.5. Interaction between aperiodic and periodic components and its influence on the rating response per window

A credible interaction between β_1_ power and the aperiodic exponent emerged at the 3|4 threshold during contemplation (T6; β = −0.18, [−0.37, −0.01]; Class 3, Model 9). No interaction was observed in C3. Independently of this interaction, β_1_ and β_2_ continued to reduce the probability of crossing the highest threshold (3|4) at T6; in this specification β_1_ no longer affected the 1|2 threshold, whilst β_2_ showed a new effect at the intermediate threshold (2|3).

Adding individual random slopes for α, β_1_, β_2_, γ, and χ (Model 11) improved predictive accuracy (see §3.7.1) and concentrated all credible fixed effects selectively at the 3|4 boundary: β_1_ (β = −0.33, [−0.64, −0.05]), β_2_ (β = −0.30, [−0.54, −0.03]), and the β_1_ × χ interaction (β = −0.22, [−0.45, −0.01]), with slightly wider credible intervals relative to Model 9 (Figure 7).

**Figure 7.**
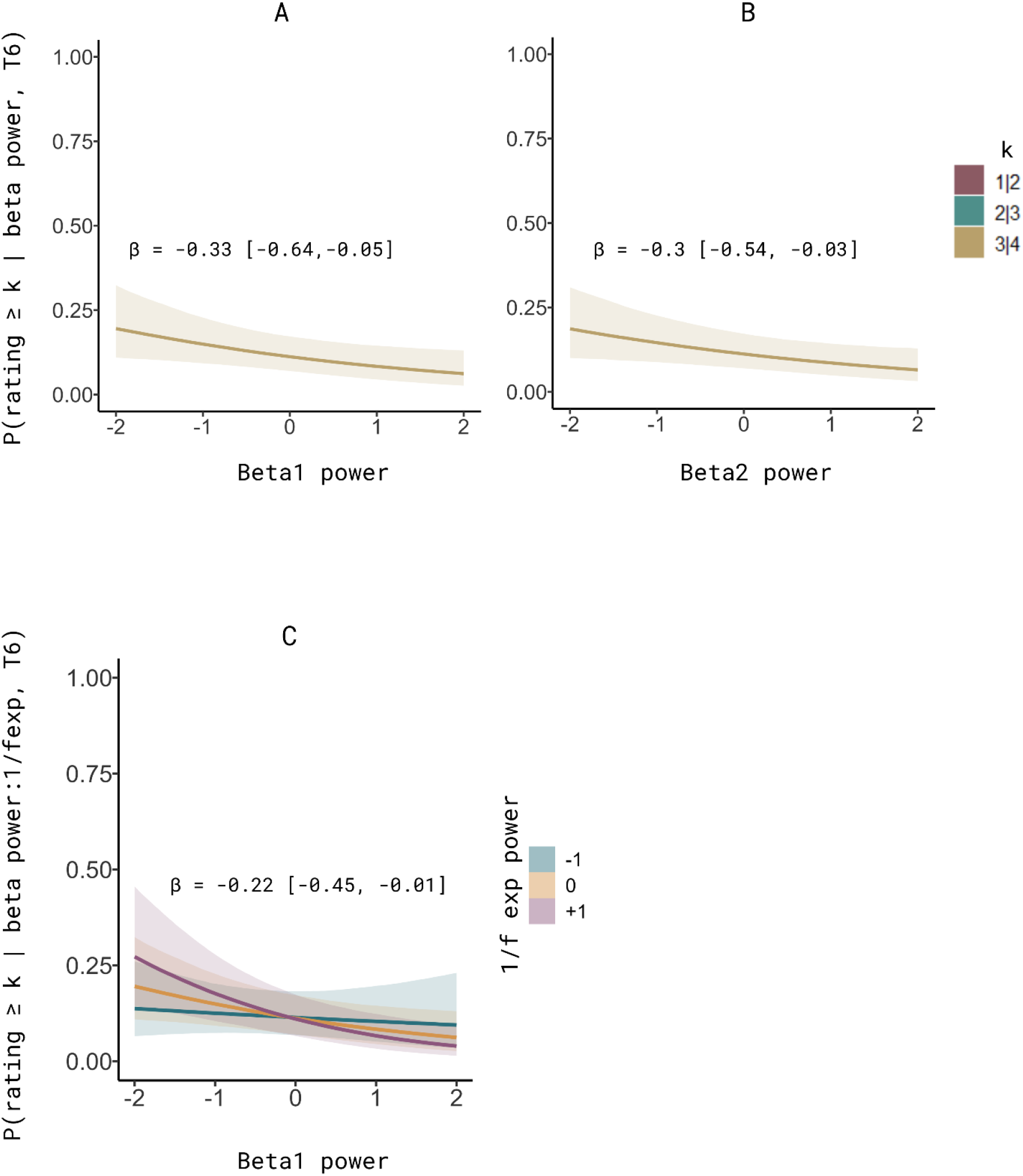
Interactionist model for the contemplation window, T6 (model 11, plus α, β1, β2, γ *and aper*_*i*_*od*_*i*_*c* χ as random effects). Graphs A and B show the credible posterior estimates of β_1_-band and β2-band power, respectively. These estimates selectively and negatively modulated the probability of crossing the 3|4 cut points. Graph C visualises the interaction between β_1_ and the aperiodic exponent with a negative effect on the probability of crossing 3|4. This interaction effect appears to be potentiated by the magnitude of the β_1_ power and the exponent; however, less aperiodic power seems to mitigate the negative effect. The shaded areas indicate the 95% credible intervals.

In the post-elicitor window (C3), where no periodic interaction was observed, the aperiodic exponent retained credible effects at the 1|2 and 2|3 thresholds with the same sign as in Model 8.b (Figure 8).

**Figure 8.**
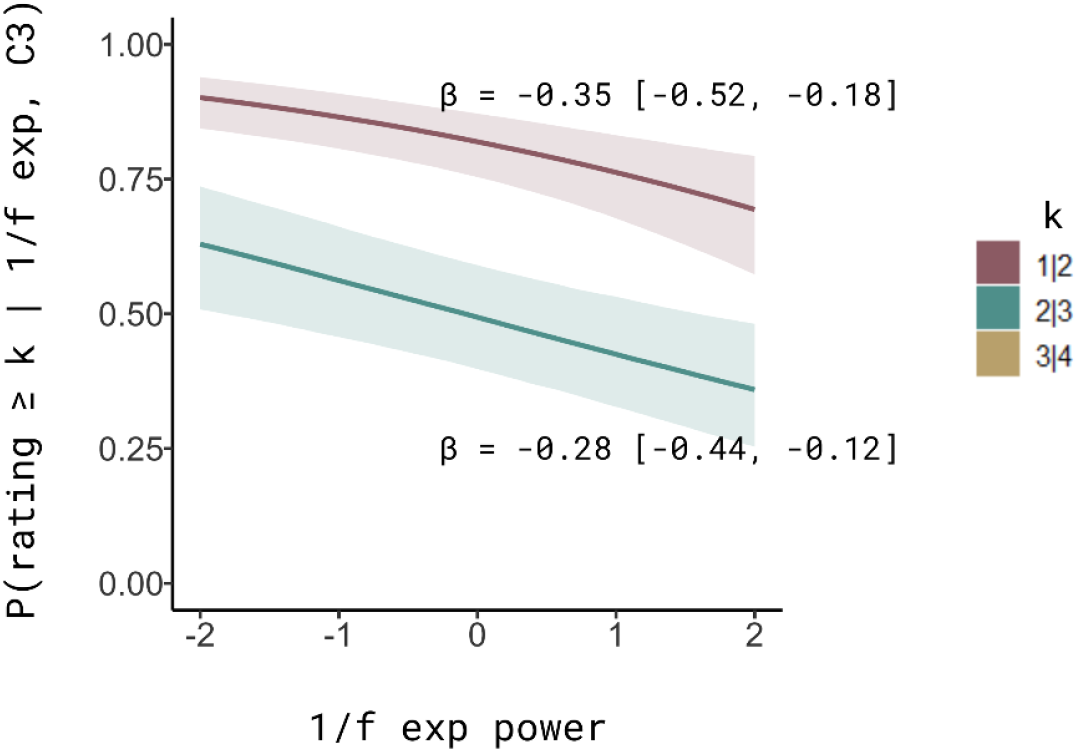
Interactionist model for the post-elicitor window, C3 (model 10). Credible conditional effects were observed for the aperiodic exponent at the first two cut-points. Posterior estimates show that, when accounting for interactions between β power and the aperiodic exponent, it is only the latter that credibly and independently modulates the probability of crossing the lower cut-points on the latent aesthetic response scale. Note that at the population level, effects were observed across the three cutpoints and certainty for the highest cutpoint (3|4) decreased when considering individual sensitivities and/or interactions. Shaded areas indicate 95% credible intervals.

### 3.6. Variance explained

Population-level spectral effects were modest but reliable, and their contribution grew with model complexity, more markedly for the T6 window than for the C3 window. Marginal R^2^ increased from less than 1% for aperiodic-only models (Model 3: R^2^m = 0.004) to approximately 7.5% for the full interactionist specification (Model 11: R^2^m = 0.075, [0.025, 0.13]). Conditional R^2^ across T6 models was in the range of 0.25-0.28, confirming that random effects (participant- and artwork-level intercepts as spectral slopes) dominated explained variance in every specification. The difference between windows is structural: during contemplation, periodic features, particularly β-band power, drove the population-level signal, whereas during the post-elicitor period, only the aperiodic exponent contributed (Figures 2 and 3). Each successive modelling layer, periodic structure, individual sensitivities, interactions, captured additional systematic variance above individual differences, even when this gain was too small relative to total variance to shift LOO rankings appreciably (§3.7).

#### 3.7.1 Predictive accuracy: contemplation window (T6) models

Model 11 (interaction + individual slopes) and Model 5 (periodic + individual slopes) were the two best-performing specifications by LOO, with a difference of only −2.5 (SE = 3.9) – well below the threshold at which models begin to differ meaningfully (Table 3). All remaining models showed elpd differences exceeding 30 units from the best specification, indicating substantially worse predictive accuracy. Adding interaction terms did not improve out-of-sample prediction beyond individual spectral sensitivities alone, though marginal R^2^ continued to increase (§3.6).

#### 3.7.2. Predictive accuracy: post-elicitor window (C3) models

Model 8.b (exponent-only + individual slopes) achieved the best predictive accuracy and was practically indistinguishable from Model 8.a, which retained the aperiodic offset (elpd_diff = −2.5, SE = 1.2; Table 4). Periodic models performed substantially worse: Model 2 (periodic, population-level) fell nearly 50 elpd units below the best specification (elpd_diff = −49.3, SE = 12.2), confirming that oscillatory power did not contribute to rating prediction during C3.

Compared to T6, the R^2^ pattern in C3 was more compressed. Conditional R^2^ ranged from 0.23 to 0.26, and marginal R^2^ remained low across all specifications (0.01–0.027). The aperiodic models with individual slopes achieved the highest marginal R^2^ (Models 8.a and 8.b: ≈ 0.026–0.027), consistent with limited population-level signal in this window. Adding interaction terms (Model 10: R^2^m = 0.019) did not improve upon the simpler aperiodic specification.

### 3.8. Synthesis: a threshold-by-window architecture of spectral effects

The results reveal a structured asymmetry across temporal windows and rating thresholds. The latent response space showed asymmetric threshold spacing, suggesting that the transition to peak aesthetic experience requires crossing a wider latent interval than transitions between lower categories.

Both aperiodic and periodic spectral parameters showed high inter-individual variability, confirming that spectral heterogeneity is a structural feature of the data worth attending to (§3.1).

The best model for the artwork contemplation window (11, Table 3) showed that the 3|4 boundary was modulated by oscillatory signatures: β_1_ and β_2_ power, and the interaction between β_1_ and the aperiodic exponent (Figure 7). β_1_ and β_2_ effects were spatially concentrated over centro-posterior and parietal regions, emerging around 1000-2500 ms post-onset (§3.3). Only clusters in β_1_ and between 2000-3500 ms distinguished the lowest from higher ratings. α and γ, by contrast, carried no robust population-level effect during T6 once individual sensitivities were modelled, marking them as individually structured rather than shared. The best model for the post-elicitor window (8.b, Table 4) showed that the three boundaries were dominated by the aperiodic exponent (Figure 6). Model 8.b is retained on grounds of parsimony, as the effects at 2|3 and 3|4 lost credibility once individual sensitivities and/or interactions were considered (Figure 8).

Together, shared spectral signatures (β_1_, β_2_, χ) and individual ones (α, γ) constitute complementary levels of a threshold-by-window architecture, suggesting that both common neural mechanisms and individual differences in spectral organisation contribute to how intensity differences in aesthetic experience might be encoded. The evidence suggests a non-uniform effect of periodic and aperiodic components on the probability of the highest rating during artwork contemplation, and that cortical excitability might distinguish the lowest from any higher, all against a background of structural inter-individual differences.

## 4. DISCUSSION

### 4.1. Threshold-like neurodynamic evidence for being highly moved by artworks

The results indicate that periodic and aperiodic spectral features exert credible, selective effects on the probability of crossing rating category cutpoints along the latent dimension of the aesthetic response intensity. Notably, these effects are not uniform: they differ across temporal windows, rating cutpoints, and variance structure, suggesting distinct patterns of association that are consistent with functionally differentiated neural mechanisms that might operate at different levels and stages of the unfolding aesthetic experience. More particularly, the results are evidence in favour of the hypothesis here tested about peak responses being associated to qualitatively different neurodynamics from less intense responses.

The geometry of the modelled latent response space (Figure 9), and more importantly the selective threshold sensitivity of spectral features, provides evidence for the threshold hypothesis. The interactionist model (Model 11/T6 window), consolidated the main credible fixed and random effects, it achieved the best predictive accuracy and explained variance. In this model, the highest rating cutpoint (3|4) was positioned farther from the centre of the latent scale than the lower thresholds. The resulting inter-threshold ratio (Liddell & Kruschke, 2018; Bürkner & Vuorre, 2019), (*k*_3|4_ − *k*_2|3_)/(*k*_2|3_ − *k*_1|2_) ≈ 1.19 was consistent across model specifications. On its own, this asymmetry is descriptive without a formal comparison against an equidistant specification. However, the stronger evidence comes from the category-specific structure. Different neural predictors associate with different thresholds, low-frequency (β_1_ and β_2_) β power at 3|4, the aperiodic exponent at 1|2 and 2|3, at both population and individual levels. If the latent geometry were an artefact of response bias or category usage, there would be no principled reason for neurophysiologically distinct mechanisms to carve the space at different joints. This selective mapping provides external validation that the latent space tracks functionally distinct transitions in the response process, consistent with a non-linear transition into peak aesthetic engagement.

**Figure 9.**
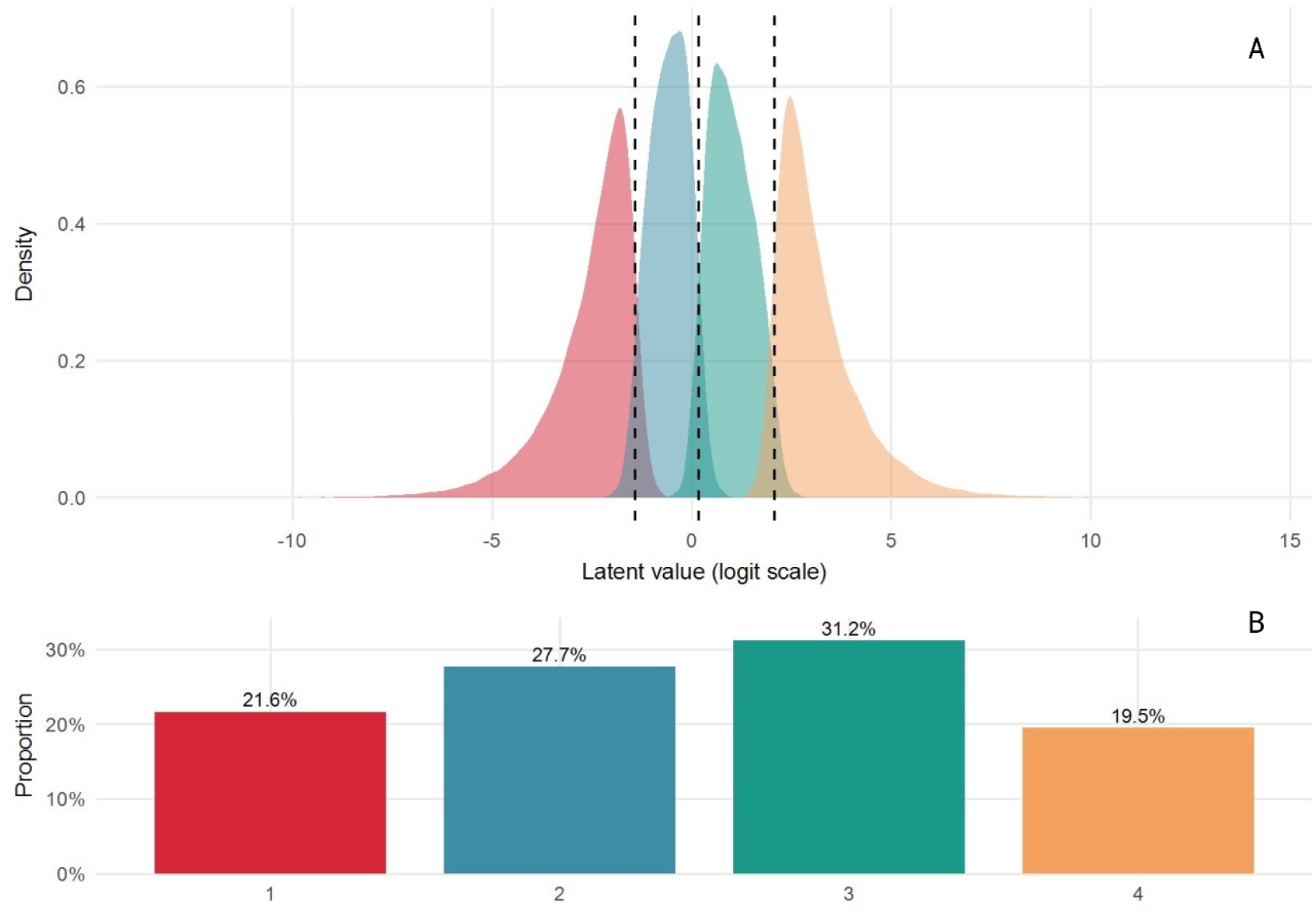
Latent response distributions and observed ratings. Panel A shows the posterior densities of the latent variable for each rating category and the κ cutpoints (dashed lines) from model 11 (the interactionist CSCLMM with better predictive accuracy and variance explained). Panel B displays the observed frequency of ratings.

However, precautions must be taken from the philosophy of measurement (Borgstede & Eggert, 2023). As Torretti (1990/2012) argues, the theoretical terms in a formal model acquire meaning not from their mathematical definition alone but from the network of empirical constraints and theoretical commitments that anchor them to the phenomena they purport to describe. The CLMM latent space is not inherently psychologically meaningful; meaning arises only when linked to substantive psychological theory and external validation (Borsboom et al., 2004). While an ordinal and Bayesian framework improves interpretatibility, estimation and model checking, these contribute to epistemic suitability (Bürkner & Vuorre, 2019), not ontological guarantees about the nature of psychological states. Inappropriate latent structures can still be severely biased yet well-fitting (Michell, 1997), and reification of well-fitting latent models remains a risk in psychological measurement (Borsboom et al.,2004).

Then, the present findings are consistent with, but do not constitute, theory-based measurement in the sense of Borgstede and Eggert (2023). The latent scale is theoretically motivated and externally validated by behavioural and neural predictors, but it does not yet constitute a theoretical term derived from a formalised theory of aesthetic experience. Its psychological interpretability rests on the convergence between predicted and observed geometry, not on formal derivation. In this study we rely on aesthetic and emotion theory (Cova & Deonna, 2014; Cullhed, 2020; Figal, 2015; Kuppens et al., 2013; Menninghaus et al., 2019; Sloterdijk, 2017) and empirical aesthetics evidence (Belfi et al., 2019; Cela-Conde et al., 2013; Durkin et al., 2025; Fiske et al., 2017; Menninghaus et al., 2015; Vessel et al., 2012, 2013, 2019; Williams et al., 2018) to motivate and guide our hypotheses and models, so our observations constitute evidence to the extent that the pertinence of these theoretical frameworks is accepted. We recognise the experiential richness and complexity of aesthetic phenomena, as well as the theoretical difficulty involved. Therefore, we regard further phenomenological description as important and scarcely integrated – consider the arguments of Bitbol and Petitmengin (2017) and Wassiliwizky & Menninghaus (2021), and these examples: Petitmengin et al. (2019), Starr & Smith (2021) and Høffding et al. (2022). A phenomenal level might provide a basis for formal approaches, as it can refine and constrain evidence and concepts, as well as generate questions and highlight gaps.

A rating-sensitive temporal dissociation, also organises the findings. Population-level oscillatory effects, specifically β_1_ and β_2_ power band activity emerged during active artwork contemplation (T6), where they selectively reduced the probability of crossing the highest rating cutpoint (3|4). In contrast, the aperiodic 1/f exponent operated in the post-elicitor window (C3), modulating negatively threshold-crossing probability across the three cutpoints of the response scale. These two spectral components, the periodic and aperiodic ones, converged during contemplation (T6) through a non-additive interaction between the power of β_1_ and the 1/f exponent, and selectively at the 3|4 threshold. Individual spectral differences add a further layer of specificity, detailed in §4.4

We will now consider each of these elements in turn. We will examine the temporal dissociation between the influences of periodic and aperiodic components on contemplation and post-elicitor windows. This can be interpreted as the broad context in which more specific accounts of the functional role of aperiodic exponents, β_1_ and β_2_ power appear to shape the neurodynamic landscape of being moved by art.

### 4.2. Temporal dissociation between periodic and aperiodic signals

The most organising feature of the results is a differential pattern in which spectral features carry credible associations with aesthetic intensity across the two temporal windows. During artwork contemplation (T6), oscillatory β power showed the most precise and consistent effects on threshold-crossing probability. During the post-elicitor interval (C3), the aperiodic 1/f exponent emerged as the dominant predictor. Importantly, this pattern held across model specifications, including those with different random-effects structures and predictor combinations.

These observations can be interpreted by its contrast with the temporal dynamics reported by the fMRI study of Belfi et al. (2019). This study used a comparable paradigm (static artworks viewed for 1, 5, and 15 seconds, followed by a post-elicitor period) to examine how large-scale brain networks might respond across the time-course of aesthetic experience. Their central finding was that for highly rated artworks, the DMN response was time-locked to both stimulus onset and offset: regardless of viewing duration, the signal returned to baseline in coordination with the disappearance of the image, while for non-moving artworks this temporal alignment was absent (inconsistent dynamics).This specifies differences in the neurodynamics of DMN engagement during more or less intense aesthetic encounters: one coupled to the stimulus, and one regarding its absence.

Of particular relevance to our design is the 5-second exposure condition, closest in duration to our 6-second contemplation window. In this condition, the differentiation between high- and low-rated trials was not confined to the stimulus period. At one second after image offset, DMN activity (particularly the posterior cingulate cortex) the lateral visual network, the caudate, and the nucleus accumbens all continued to distinguish between levels of aesthetic intensity, with differentiation persisting for up to three seconds past offset. Yet the behavioural data from the same study tell a different story about the experiential timecourse: the slow component of their pleasure decay model (τ_long ≈ 425 s) indicates that felt pleasure persists at a low level for minutes after stimulus offset. Something sustains the behavioural response at a timescale that regularly is not studied in laboratory settings. However, and more specifically, temporal resolution of the BOLD signal is insufficient to distinguish whether the networks sustain activity or not at these timescales.

Our finding that the aperiodic exponent during the post-elicitor period (C3) predicts aesthetic intensity might speak directly to this temporal gap. The C3 window corresponds precisely to the period where Belfi et al.’s 5-second data show multi-network differentiation still active. A shallower (flatter) aperiodic exponent, reflecting a more excitation-dominated cortical state, predicted higher aesthetic intensity during this window, while a steeper exponent, indicating a shift toward inhibition-dominated dynamics, predicted lower ratings. Interestingly, when we consider population and individual levels effects of the aperiodic exponent, we observe that it influence all cutpoints, however, its influence on the first cutpoint 1|2, β =− 0. 42, across models is invariantly the strongest, reinforcing the suggestive idea that a no aesthetic engagement might imply an active inhibitory regime. We also saw cutpoint selective exponent effects on T6 only for the highest cutpoint 3|4, which might suggest that in addition to β top-down inhibitory influences, a global inhibitory regime might be particularly necessary for not crossing to a peak state.

The specificity of this effect to C3, rather than T6, raises the question of whether the aperiodic exponent indexes a state that is initiated by the aesthetic encounter or one related to the intrinsic architecture and so, antecedent to stimulation. The high reliability of individual aperiodic profiles across windows (ICC > 0.94 for both exponent and offset, §3.1) suggests that the exponent reflects, at least in part, a stable individual characteristic rather than a transient, stimulus-induced modulation. Under this interpretation, the E/I balance captured by the exponent may represent a tonic neural property that predisposes certain neural states within an individual toward more or less intense aesthetic engagement. The fact that this property becomes a credible predictor specifically during C3 may reflect the removal of stimulus-driven variance that, during T6, masks its contribution.

This interpretation suggests the role of the ontogenetic factors modulating the response, how the history of the organism shapes its encounters: the perceptual, interpretive, and associative capacities that fuel aesthetic engagement. The aperiodic exponent has shown to track developmental trajectories across the lifespan, with flatter slopes (lower exponents) observed in older adults and associated with reduced cognitive performance (Voytek et al., 2015; Donoghue et al., 2020). While the present study cannot address this directly, the substantial between-person variance in aperiodic parameters and utheir predictive relationship to aesthetic intensity suggest that individual differences in E/I balance may constitute a meaningful source of variation in aesthetic responsiveness. It is worth noting that the absence of credible aperiodic offset effects in either window. The aperiodic offset (previously known as “broadband shifts”)reflects broadband spectral power and has been associated with overall neural spiking rate (Manning et al., 2009; Miller et al., 2009; Donoghue et al., 2020). The null result here suggests that aesthetic intensity is not simply a function of how active the brain is in a global sense, but rather of how that activity is organised spectrally. More interestingly, it suggests that neurodynamics are crucial to understanding the functional differences of ongoing, complex experiences such as aesthetics, which surely rely on non-stationary dynamics rather than sustained, stereotypical activity.

#### 4.2.1 Periodic and aperiodic interaction during contemplation, T6 window

A credible interaction between β_1_ power and the aperiodic 1/f exponent emerged at the 3|4 threshold during the contemplation window (T6) (Figure 7, model 11), indicating that the influence of β_1_ on the probability of peak aesthetic response depends on the concurrent aperiodic state. This interaction was non-additive: neither the simple sum of β_1_ and exponent effects nor the individual main effects alone could account for the pattern observed at the highest threshold. No comparable interaction was detected in the post-elicitor window (C3).

The selectivity of this interaction, emerging only at the 3|4 threshold and only during active contemplation, is consistent with the broader pattern of results, in which the transition to peak aesthetic intensity appears to be governed by a qualitatively different configuration of neural dynamics than transitions between lower rating categories. If β power constrains the transition to peak engagement (as observed and argued in next section, §4.3), and if the aperiodic exponent reflects the tonic E/I context in which this “gating” operates, then evidence for an interaction suggests that beta-mediated regulation is not equally effective under all background states.

Specifically while still speculative, the combination of high β power (strong maintenance of the current processing mode) and a steep exponent (inhibition-dominated background) may represent a doubly constrained neural configuration, one in which both the oscillatory gating mechanism and the tonic E/I balance work against the transition to peak aesthetic engagement. Conversely, low β power against a flatter (more excitation-shifted) spectral background may represent the most permissive configuration for this transition.

This interpretation should be treated as such; the knowledge around the functional relationship between aperiodic and periodic components is still early. The interaction term in an ordinal regression model captures statistical non-additivity, which can arise from multiple underlying mechanisms and despite the non redundant character of periodic and aperiodic components it is not possible to distinguish the real nature of its covariance. It could reflect a biophysical interaction — for instance, if the aperiodic background modulates the efficacy or prevalence of β oscillatory bursts — or it could reflect a statistical dependency arising from the fact that both parameters are estimated from the same power spectrum, they share underlying neural mechanisms. The FOOOF parameterisation procedure separates periodic peaks from the aperiodic background precisely to minimise this confound (Donoghue et al., 2020), but cannot guarantee that their effects are independent. Future work using independent estimation methods, or direct manipulation of E/I balance (e.g., pharmacological or neurostimulation approaches), would be useful to establish whether the interaction reflects a causal coupling between oscillatory gating and background excitability.

Despite these caveats, the present finding adds to a growing body of evidence that periodic and aperiodic components of the EEG signal are functionally relevant in specific ways, and that their interaction may carry information beyond what either component conveys alone (Donoghue et al., 2020; Ouyang et al., 2020; Waschke et al., 2021). To our knowledge, this is the first report of such an interaction in the context of aesthetic experience, and it suggests that models of aesthetic processing should consider how oscillatory dynamics operate within, and are modulated by, the broader spectral context.

### 4.3. β power population level role in the aesthetic engagement

Both β_1_ (12–16 Hz) and β_2_ (16–20 Hz) power showed credible negative associations with the probability of crossing the highest rating threshold (3|4) during artwork contemplation (T6). This effect was robust: it survived the inclusion of random slopes for individual differences in band-specific power, and it was the only periodic effect that remained credible at the group level after accounting for inter-individual heterogeneity. Higher power, in the range of 12-20 Hertz range, during contemplation reduced the probability that a given trial would reach the highest aesthetic intensity, suggesting that elevated β activity constrains the transition from moderate to peak engagement.

This pattern is consistent with, and extends, the functional characterisation of β oscillations that has developed over the past two decades. Engel and Fries (2010) proposed that β-band activity supports the maintenance of the current sensorimotor or cognitive set, a “status quo” hypothesis in which elevated β power signals that the brain is actively preserving its current processing configuration rather than transitioning to a new one. Subsequent work has refined this framework in two important directions mostly from the study of working memory. First, Spitzer and Haegens (2017) argued that beta-mediated ensemble formation does not merely preserve a state but actively (re)activates endogenous content representations, holding a particular model or representation active in the service of current task demands. Haegens and colleagues (2017) demonstrated that this function is supramodal: β power tracks the content of a perceptual decision regardless of which sensory modality delivers the evidence, suggesting that β indexes the maintenance of an abstract internal representation rather than modality-specific processing. Second, Lundqvist and colleagues (2024) have shown that what appears as sustained β power in trial-averaged analyses is better understood as the aggregate of transient high power bursts,brief episodes during which a cortical pattern is reinstated. Higher and smooth average β power would reflect a greater rate or duration of such bursts, each of which momentarily reactivates the currently held representation, in any case single-trial burst detection would be needed. A trial-level approach can be more appropriate for functionally characterising the neurodynamics that might distinguish different levels of intensity in the experience. If we attend the arguments for cognitive processes occurring in discrete time windows, instead of sustained, stationary states, from Lundqvist and Wutz, (2022) such approach would be sensitive to real continuous transformation of constant signal into spatiotemporally coherent objects, scenes, and events that might distinguish not only invariants to the neurodynamics according to the intensity felt but differences in its implementation, which seems core to our phenomena governed by diversity.

If β oscillations carry and reinstate an internal model, then elevated β power during artwork contemplation would reflect the viewer actively maintaining a particular interpretive or evaluative representation of the artwork, a prior, in the broad sense, against which incoming sensory information, its encoding and reactivation might be reflected in burst of γ dynamics (Lundqvist et al., 2016, 2018). Lower β power for more intensely rated trials would then indicate that this prior is being held less tightly: the system spends less time reinstating its current model, creating conditions under which the sensory input can drive processing more freely and the viewer can transition into a less constrained mode of engagement. Time frequency contrast (Figure 10) between broadband activity for rating 4 trials and each of the other category responses aligns with this interpretation, events of high power in β_1_, particularly for rating 1, 2 (shown as intermittent blue activity between *~* 1 − 2. 5 *s* and *~* 5 − 6 *s* in Figure 10) suggests an initial dominance of such priors, along with differences in what appears to be bursts of γ, the way and extent to which new information is encoded and retrieved.

**Figure 10.**
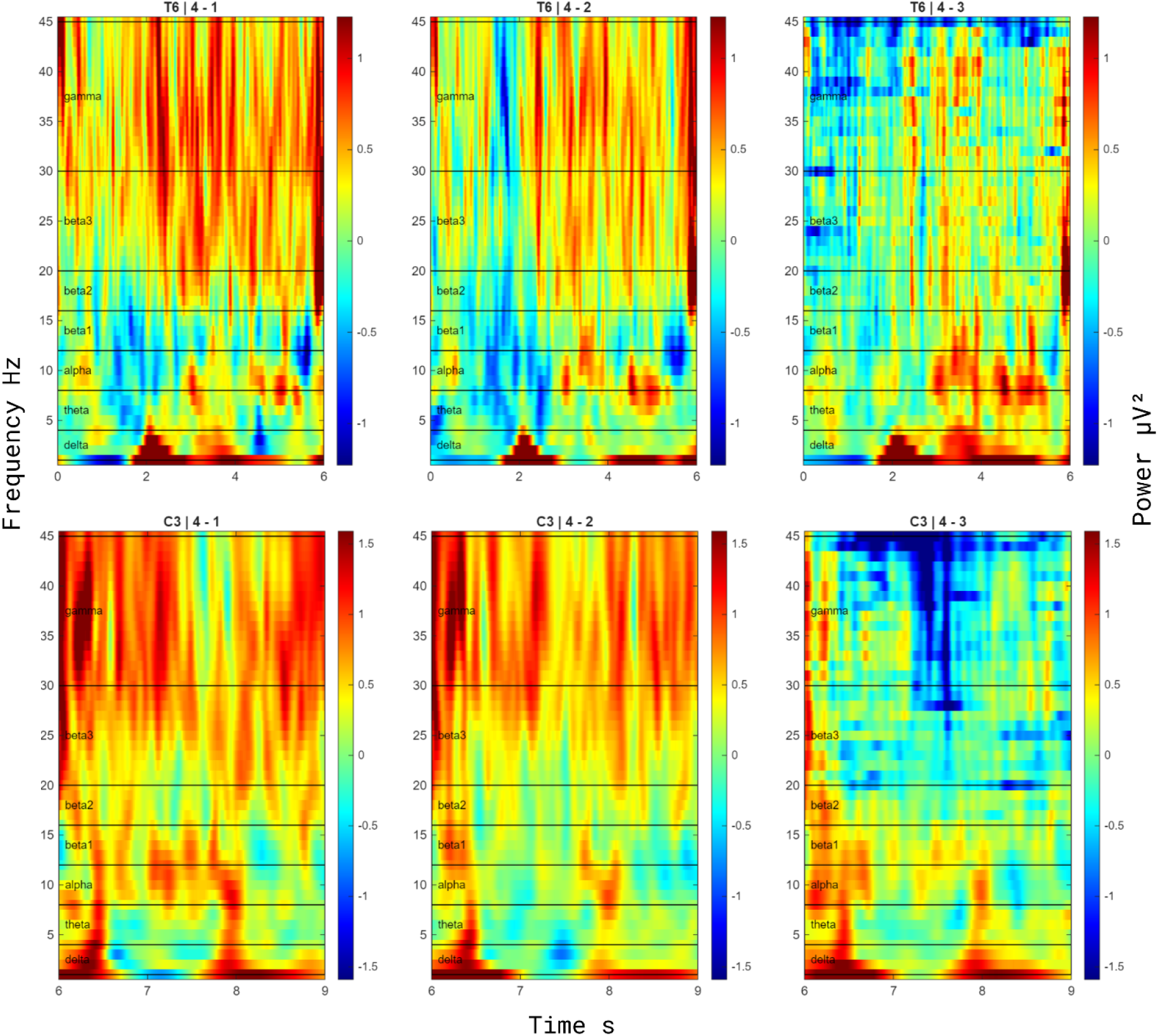
Time-frequency contrasts of broadband power during contemplation (T6, top row) and post-elicitor (C3, bottom row) windows. Each panel displays the difference in ROI-averaged power (μV^2^) between rating 4 trials and trials of a single lower rating category (left to right: 4 − 1, 4 − 2, 4 − 3). Cool colours indicate higher power for the lower rating category; warm colours indicate higher power for rating 4. During T6, intermittent negative deflections in the β_1_ range (12–16 Hz) are visible in the 4 − 1 and 4 − 2 contrasts, concentrated between approximately 1–2.5 s and 5–6 s post-onset, consistent with transient β bursts that are more prevalent for lower-rated trials. Positive deflections in the γ range (30–45 Hz) emerge in the 4 − 3 contrast during mid-to-late contemplation (T6, right panel), suggesting frequency-specific differentiation at the boundary between moderate and peak aesthetic responses. During C3, broadband differences are less structured, consistent with the absence of credible periodic effects in this window.

The sub-band structure of the effects is informative in this regard. β_1_ (12–16 Hz) showed credible effects at both the lowest (1|2) and highest (3|4) thresholds, suggesting that it might operate as a general gating mechanism across the evaluative range. β_2_ (16–20 Hz), by contrast, was selective to the highest threshold. This dissociation is in line with evidence from the motor and decision-making literature that lower and upper portions of the β range serve partially distinct functions: frequencies below approximately 20 Hz have been more consistently associated with the maintenance of internal models and prior expectations, while higher β frequencies have been linked to anticipatory and attentional processes (Kilavik et al., 2013; Betti et al., 2021). Betti and colleagues (2021) specifically identified the lower β range as the spectral signature of the brain’s intrinsic dynamic architecture, the frequency at which resting-state connectivity patterns are preserved during natural vision and through which hub regions of the default mode, dorsal attention, and somatomotor networks coordinate their integration. The fact that β_1_, situated squarely within this range, showed the broadest effects across the rating scale is consistent with the idea that it indexes a general prior-maintenance function that modulates aesthetic engagement at multiple levels of intensity. β_2_’s selectivity to the 3|4 threshold may reflect a more specific role in mediating the final transition into peak aesthetic experience, though the functional basis of this selectivity remains to be established. The tripartite partition adopted here was defined prior to modelling; while the precise boundaries between β sub-bands remain unsettled, the differential pattern across the 12–16 and 16–20 Hz ranges is an empirical observation of this study, not an assumption. Betti et al. (2021) recognise that β can be divided into at least two sub-bands, low β (13–20 Hz) and high β (20–30 Hz). Our results converge with this broader distinction, effects concentrated entirely within the low-β range, while suggesting that finer partitioning within that range may capture functionally relevant differences, as β_1_ and β_2_ showed distinct threshold sensitivities despite both falling within Betti et al.’s low-β band.

In the framework of predictive coding, these findings admit a further interpretation, although developed primarily in the sensory cortex for relatively simple perceptual events, and extending them to a 6-second aesthetic contemplation involves a significant leap. β oscillations have been proposed to carry top-down predictions down the cortical hierarchy, while γ oscillations carry prediction errors upward (Arnal & Giraud, 2012; Bastos et al., 2012; Betti et al., 2021). Under this scheme, elevated β during contemplation would correspond to the imposition of strong top-down predictions on sensory processing, and the graded β suppression observed for more intensely rated trials would correspond to a weakening of those predictions, allowing prediction errors generated by the artwork to propagate more effectively and drive model updating.

#### 4.3.1. Temporal and spatial dynamics of β effects

Cluster-based permutation analyses on time-resolved β power during T6 provided converging evidence for the threshold-specific pattern identified in the ordinal models, while adding temporal and spatial information (§3.3). Across all contrasts, 1 vs. 2-3-4 (idem est dicere 1|2) and 1-2-3 vs. 4 (idem est dicere 3|4), β_1_ and β_2_ power bands were consistently higher for lower-rated trials, confirming the direction of the effect observed through modelling.

It is important to note that cluster-based permutation tests provide inference about the existence of an effect, they reject the null hypothesis that there is no difference between conditions anywhere in the spatio-temporal space, but do not allow for strong conclusions about the precise spatial or temporal boundaries of that effect (Maris & Oostenveld, 2007). To partially remedy this, we created subwindows of 500 ms for the comparison, which doesn’t mean clusters last that long, just that they appeared there. The topographic and temporal patterns reported here should therefore be interpreted as indicative rather than definitive. With this caveat, two observations merit attention.

First, the temporal ordering of effects across thresholds is suggestive. Clusters at the 3|4 boundary emerged earlier (from 1000 ms) than effects at the 1|2 threshold (from 2000 ms). If this pattern is reliable, it could indicate that the neural processes distinguishing peak from non-peak aesthetic responses are established earlier during contemplation than those distinguishing minimal from moderate engagement, consistent with the possibility that the transition to peak aesthetic intensity involves an early configuration of cortical state rather than a gradual accumulation over time. However, given the inferential limitations of cluster-based methods, this temporal ordering should be treated as hypothesis-generating rather than confirmatory. We would like to point out a stimulating observation from the phenomenology of being moved by art that seems to converge with the idea of an early realisation predictive of the intensity of the experience. According to Høffding et al’s study (2022) which relies on interviews and theoretical understanding, the aesthetic experience is an unfolding process characterised by an initial aesthetic “seizure” where a “direct and unreasoned impression comes first” –quotation from Dewey (1934/1980) – that precedes all stable configuration of what the art in question is about.

Second, the clusters’ topography suggests that rating-related β power differences are concentrated over posterior rather than anterior regions. This distribution is consistent with the known source topography of low-β oscillations: TMS perturbation and MEG atlasing studies have shown that natural frequencies in the low-β range originate predominantly from lateral occipito-parietal regions, whereas high-β oscillations are generated in motor and prefrontal areas (Rosanova et al., 2009; Ferrarelli et al., 2012; Capilla et al., 2022, as argued in Di Dona and Ronconi (2023) Figure 1). Complementary evidence from Samaha et al. (2017) indicates that pre-stimulation β power over posterior parietal cortex, but not occipital α power, predicts TMS-induced phosphene perception, further establishing β as the functionally dominant rhythm of parietal networks.

This posterior β activity does not operate in isolation. Hillebrand et al. (2016) demonstrated, using MEG source-reconstructed phase transfer entropy, that information in the β band flows predominantly from parieto-occipital regions toward frontal areas, while the reverse anterior-to-posterior flow occurs in the θ band; forming a frequency-specific reentry loop through which information reverberates between posterior and frontal subsystems. The posterior β effects observed in our data should therefore be understood not as reflecting purely local processing but as likely participating in this broader recurrent architecture.

Di Dona and Ronconi (2023) have proposed that posterior β oscillations specifically support the magnocellular-dorsal visual pathway, providing coarse spatial representations that, through feedback from parietal cortex, guide attention-demanding process of object identification in ventral areas. Within this framework, β modulation is linked to tasks requiring spatial integration and segregation, perceptual reorganisation, and figure-ground parsing, visual processes that are central to the playful exploration of artworks. That our posterior β clusters differentiated aesthetic intensity ratings is compatible with this account, insofar as contemplating visual art involves precisely the kind of active spatial engagement, exploring compositional configurations, parsing figure from ground, integrating local elements into global structure, that recruits dorsal-stream β mechanisms. The observed β desynchronisation for higher-rated trials may thus reflect a release from spatially constrained perceptual analysis, consistent with the transition toward more integrative processing that characterises intense aesthetic experience.

### 4.4. Individual differences on the neurodynamics of the aesthetic response

A consistent finding across models was that inter-individual variability was a structured and informative component of the data. Two independent observations ground this inter-individual variability as signal rather than noise. First, the spectral features are highly reliable across temporal windows (§3.1), so the between-subject variance reflects stable individual characteristics rather than measurement error; second, incorporating that variance as random slopes improves out-of-sample predictive accuracy (Tables 3–4), and the corresponding variance components are recoverable at this sample size (Figure 11), so the gain is not attributable to overfitting.

**Figure 11.**
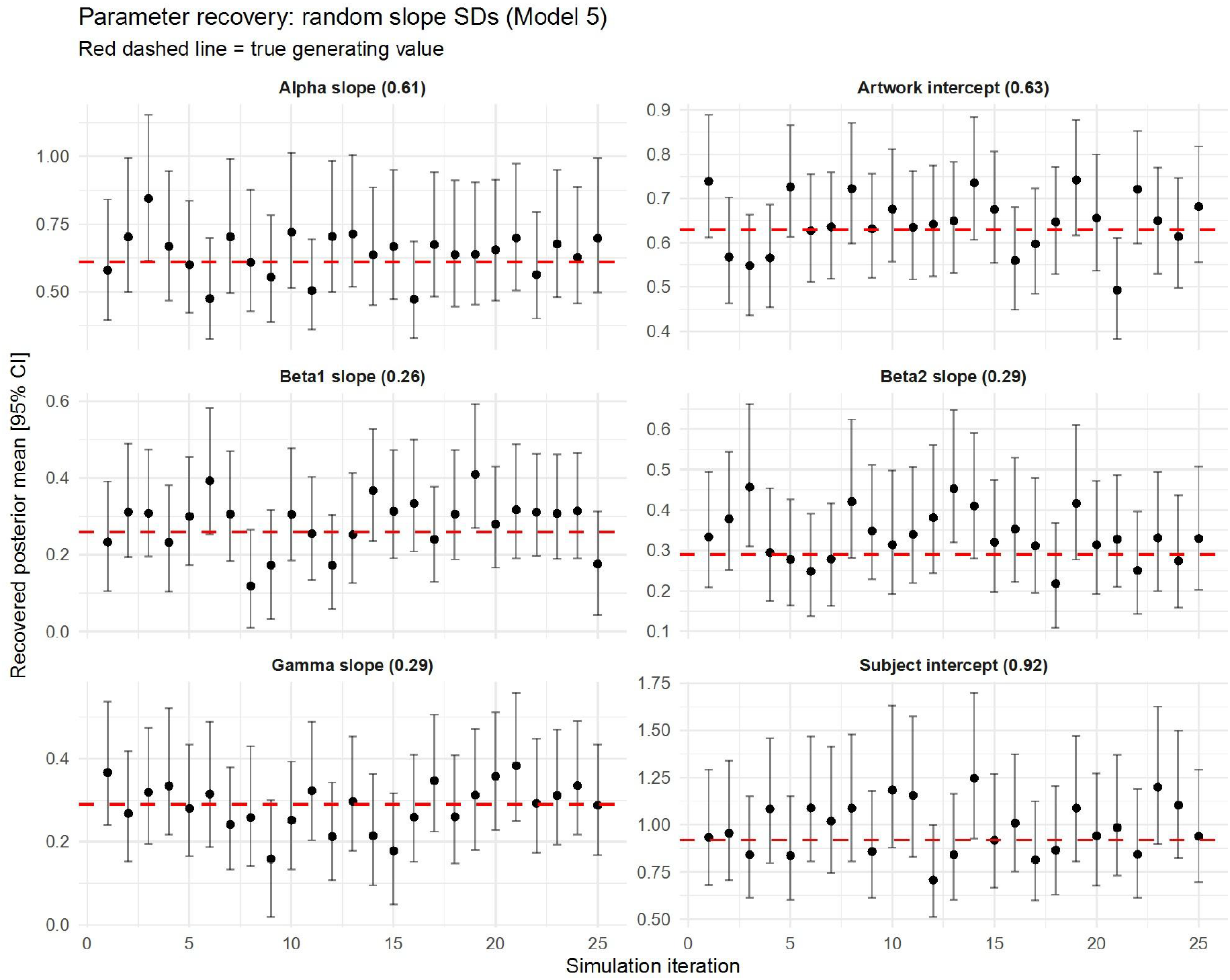
Parameter-recovery simulation for Model 5 (periodic predictors with individual slopes). Synthetic datasets (n = 25) were generated from the fitted posterior and the model refitted to each; points show the recovered posterior mean and bars the 95% credible interval per iteration, with the red dashed line indicating the true generating value. Random-slope SDs were recovered with acceptable accuracy, the generating value falling within the 95% interval in the large majority of iterations for every parameter. Small SDs (β_1_, β_2_, γ ≤ 0.29) showed modest upward bias, consistent with posterior right-skew at the non-negative (zero) boundary when the per-parameter likelihood is weak at n = 22, whereas larger SDs (α = 0.61; artwork intercept = 0.63) were estimated precisely.

Random-effects estimates revealed substantial variability at both the subject level (Model 11, the interactionist: SD = 1.08, [0.75, 1.52]) and the artwork level (SD = 0.67, [0.56, 0.80]), indicating that both who is viewing and what is viewed contribute meaningfully to aesthetic response intensity. These behavioural random effects persisted after the inclusion of spectral predictors, indicating that oscillatory and aperiodic features account for only part of the variance in aesthetic ratings. Including random slopes for frequency-band and aperiodic terms consistently improved predictive accuracy (Tables 3–4), and the conditional variance explained exceeded the marginal (population-level) variance by one to two orders of magnitude (Table 2). In other words, spectral inter-individual variance is a structural component of the intensity of the aesthetic response.

Including random slopes for frequency-band power also revealed a specific pattern of individual variability (Model 5). α power absorbed the largest share of between-person variance (SD = 0.61), substantially exceeding that of β_1_ (SD = 0.26), β_2_ (SD = 0.29), and γ (SD = 0.29). That is, individuals differed most in how α power related to their aesthetic ratings: for some, higher α was associated with more intense responses; for others, the opposite held. This heterogeneity was sufficiently large that, once individual differences were accounted for, α no longer showed a credible group-level effect. The same was true of γ, whose positive association with the 3|4 threshold in the population-level model disappeared after individual variability was modelled. The contrast between α/γ and β is therefore not that the former are unrelated to aesthetic intensity, but that their relationship is idiosyncratic, varying substantially in direction and magnitude across individuals, whereas beta’s relationship is more consistent at the group level while still revealing individual structure. A similar pattern was observed for the aperiodic exponent, with subject-level SDs of 0.30 in T6 and 0.48–0.49 in C3.

To be sure whether the individual-difference structure was identifiable at this sample size (N=22), we conducted a parameter recovery simulation for Model 5 since parsimony; models with an absolute elpd difference below 4 (−2.5, Table 3) are considered practically indistinguishable in predictive accuracy (Sivula et al., 2025). Synthetic datasets were generated from the posterior estimates of the fitted model (n = 25 iterations), and the same model was refitted to each. Random slope SDs were recovered with acceptable accuracy across all parameters (Figure 11): larger SDs (α: 0.61; artwork intercept: 0.63) were estimated precisely, while smaller SDs (β_1_: 0.26; β_2_, γ: 0.29) showed modest upward bias, attributable to posterior right-skew near the zero boundary when the likelihood is weak at n = 22 The subject intercept SD (0.92) showed slight overestimation, as expected when between-person variance is absorbed by a single grouping term. These results support the interpretability of the random-effects structure under the present design constraints.

This heterogeneity carries methodological and theoretical implications. Methodologically, averaging spectral effects across individuals obscures meaningful signal rather than recovering a common pattern; these differences are better understood by modelling them directly. Theoretically, the structured nature of the variability raises a question about the underlying architecture: what kind of neural organisation could produce both the consistency of β-band effects and the diversity observed in α, γ, and aperiodic components? A large-scale network perspective, considered next, offers one way of approaching this question.

### 4.5. A network-level interpretation

Before turning to that question, two null results require comment. Neither θ nor delta power showed credible associations with aesthetic intensity at any threshold in either temporal window. The θ null is noteworthy given the theoretical expectations linking θ oscillations to the DMN: Strijbosch et al. (2021) predicted that midfrontal θ power would decrease for highly moving artworks, based on EEG-fMRI evidence that midfrontal θ indexes DMN activation (Scheeringa et al., 2008, 2009). Their data did not support this prediction either; no θ differences emerged across their three response categories. Strijbosch et al. suggested that the θ–DMN link may be specific to the contexts in which it was originally established (resting state, working memory) rather than generalising to aesthetic processing. Our results converge with this interpretation using a different analytic framework (Bayesian ordinal models with category-specific effects) and a different operationalisation of intensity (four-level graded ratings rather than tertile splits).

This convergent null does not imply that the DMN is uninvolved in aesthetic experience, a conclusion that would contradict substantial fMRI evidence (Vessel et al., 2012, 2013; Belfi et al., 2019; Williams et al., 2018; Durkin et al., 2025). It implies that scalp-level θ power is not the appropriate index for the present analytic design. The relationship between oscillatory bands and large-scale networks is not one-to-one. Samogin et al. (2020) demonstrated that different resting-state networks show preferential intra-network connectivity in different frequency bands: the DMN and visual network in α, the dorsal attention network in beta, the ventral attention and language networks in γ. Inter-network connectivity also varies by frequency, meaning that the same pair of networks may be coupled in one band and decoupled in another. A scalp power analysis, as conducted here and by Strijbosch et al., captures the amplitude of local oscillatory activity but cannot distinguish whether that activity supports within-network coherence, between-network communication, or neither. The absence of θ effects therefore constrains interpretation at the level of spectral power, not at the level of network engagement.

This distinction reframes the individual differences observed in our data. All spectral features that showed credible effects, α, β_1_, β_2_, γ, and the aperiodic exponent, also showed structured inter-individual heterogeneity. The diversity of functional configurations through which different individuals achieve intense aesthetic engagement is consistent with the principle articulated by Sporns and Kötter (2004): that evolved neural architectures maximise the diversity of functional motifs on a constrained structural backbone. We do not observe network motifs directly, but the diversity of functional states they may enable. That different oscillatory bands preferentially support connectivity within different large-scale networks (Samogin et al., 2020; Hillebrand et al., 2016; Bressler & Menon, 2010) suggests a plausible mechanism: individual differences in band-specific power may reflect differences in which network configurations are recruited during aesthetic experience, rather than differences in the magnitude of a single shared process.

It is important to clarify that “structural” in Sporns and Kötter refers to anatomical connectivity, the physical wiring of the brain, not to functional regularity across individuals. The inter-individual variability we observe is functional: different spectral profiles associated with the “same” experiential outcome. The inference from functional diversity to structural motif diversity is indirect and would require connectivity-based evidence that the present study cannot provide. Testing this interpretation will require approaches capable of mapping the spectral heterogeneity identified here onto distinct patterns of large-scale network dynamics: EEG functional connectivity, source localisation, or simultaneous EEG-fMRI. The temporal dimension offers a distinctive advantage in this regard, as it can resolve the dynamics of networks with spatially overlapping regions, an expected feature of the high-integration states characteristic of complex experiences such as aesthetic engagement.

### 4.6. Limitations

Several limitations should be acknowledged. The sample comprised 22 participants, yielding approximately 2,470 trial-level observations per temporal window. While the Bayesian framework with LOO cross-validation mitigates overfitting concerns, and all Pareto-k diagnostics remained below 0.7, the individual-difference estimates (22 random-slope values per predictor) should be considered preliminary. Parameter recovery indicated that the random-effects structure was identifiable under the present design (see §4.4 and Figure 11), supporting our treatment of the variance components as estimable signal rather than artefact; the magnitudes of individual sensitivities, however, remain provisional and warrant replication in larger samples.

The Chilean sample and the use of Chilean artworks were deliberate design choices motivated by the goal of broadening the cultural contexts represented in empirical aesthetics as well as recognizing the situated character of art production and reception; however, this specificity obviously limits generalisability to other populations and stimulus sets.

As argued in §2.3, the fixed order of artwork presentation was a deliberate theoretical choice, though not free of limitations. Aside from the advantage the mixed model offers by absorbing random variation, we examined rating distributions across session thirds, which revealed a modest shift from rating 2 to rating 4 over the session (Figure 12). This could reflect participants refining their evaluative criteria as the session progressed, a process that is itself theoretically relevant to aesthetic engagement rather than a simple limitation or confound. Still, to assess whether this pattern threatened the spectral findings, trial position was added as a covariate (Table 18A): β-band effects remained credible and unchanged in magnitude, and trial position itself showed no credible independent association with ratings (β = 0.09, [−0.05, 0.24]), indicating that the spectral findings are not impacted by session progression.

**Figure 12.**
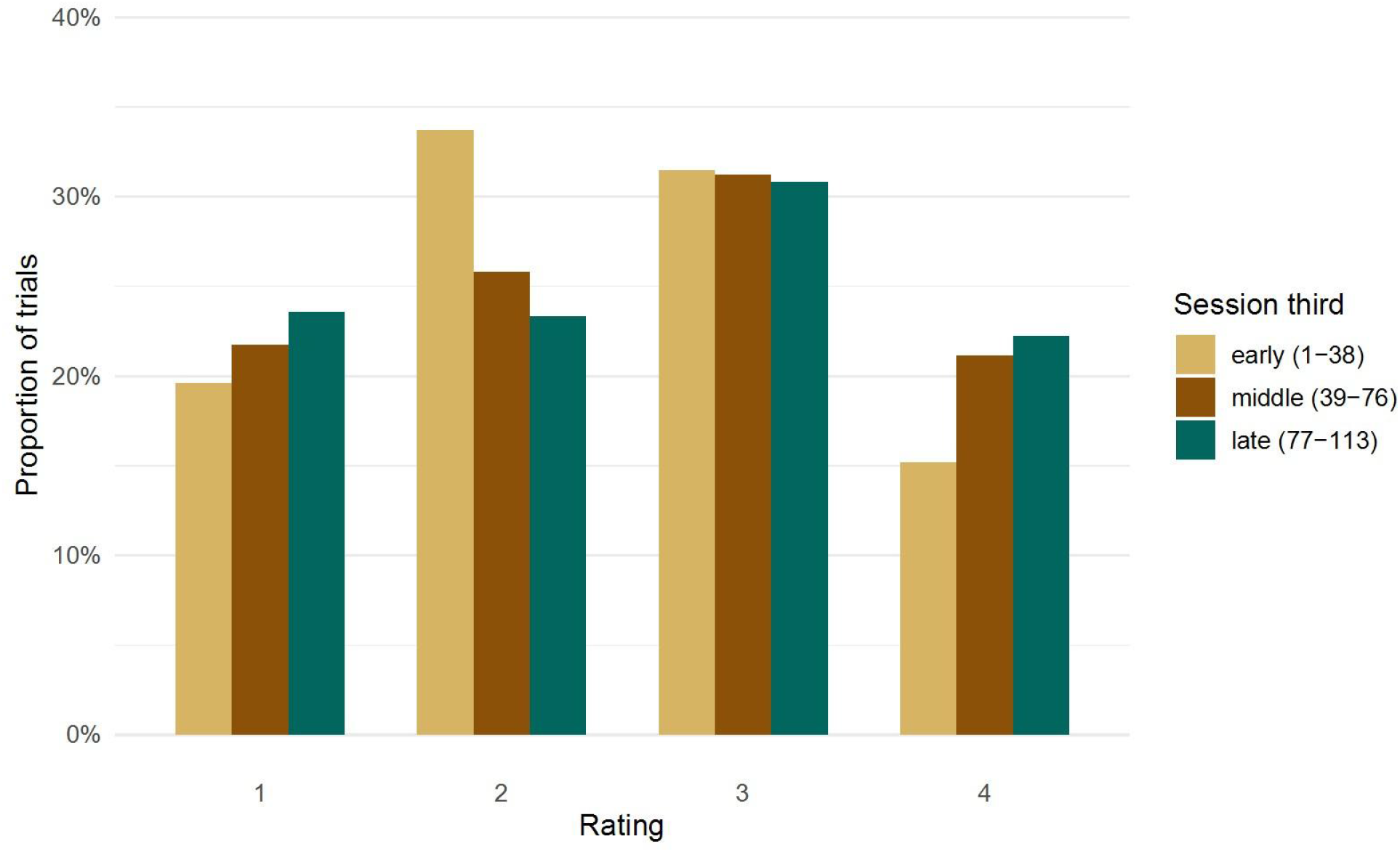
Rating distribution across session thirds. Bars show the proportion of trials assigned to each rating category (1–4) within early (trials 1–38), middle (39–76), and late (77–113) thirds of the session.

The study employed single-channel ROI-averaged scalp EEG without source localisation, precluding spatial inference beyond the broad topographic patterns identified in the cluster-based permutation analyses. The main ordinal models operate on epoch-averaged power spectral densities, one spectral estimate per trial per band per window, and therefore cannot resolve within-epoch temporal dynamics. The time-frequency contrasts and cluster permutation tests supplement this with temporal and spatial information, but these analyses are descriptive and separate from the cumulative models.

The complexity-progression strategy (§2.6.3) would have excluded the aperiodic offset from further modelling, as it showed no credible effects in either window. In the post-elicitor window (C3), however, suspected collinearity between exponent and offset motivated an additional check: Model 8.a retained both parameters to test whether the offset contributed beyond the exponent. It did not (elpd_diff = −2.5, SE = 1.2 relative to the exponent-only Model 8.b), confirming the stability of the exponent effects and clarifying that the staged exclusion of offset was not masking a suppressed contribution.

In the same window context, model 10 (C3 interaction model) produced 183 divergent transitions (2.3% of post-warmup draws), with funnel-like geometry visible in pairwise posterior plots (Figure 11A) involving the interaction terms and intercepts, indicating that the posterior for this specification was difficult to sample reliably. However, the key C3 finding, credible aperiodic exponent effects across rating thresholds, is established in Model 8.b, which showed no sampling difficulties.

One evident limitation is that this study used a cross-sectional design that precludes causal claims about any of the spectral-rating associations reported. Whether β power constrains aesthetic intensity, or intense aesthetic engagement suppresses beta, or both are consequences of a common upstream process, cannot be resolved with the present data. Experimental manipulation of oscillatory states (e.g., via transcranial stimulation) would be needed to establish directionality.

Finally, all models fixed the discrimination parameter to 1.0. Fixing it to a constant is the standard identification convention for cumulative logit models and merely sets the latent scale; the value itself is innocuous. What it assumes is homoscedasticity — constant latent dispersion across participants and conditions. Were discrimination instead to vary and be left unmodelled, that heteroscedasticity could be absorbed into the cutpoints and appear as unequal threshold spacing. Our latent geometry, notably the 3|4 boundary lying farthest from the scale centre, is therefore conditional on this assumption — the same caveat raised in §4.1, where we declined to treat the asymmetry as evidence in its own right. Crucially, the threshold hypothesis does not rest on the geometry: it is carried by the category-specific mapping of distinct neural predictors onto distinct cutpoints (§4.1), which is unaffected by the discrimination constraint. Modelling discrimination directly, via an unequal-variance ordinal specification, is a natural extension for future work.

## Supporting information

ANNEX

