## Supplementary material for "PERIODIC AND APERIODIC SPECTRAL SIGNATURES OF BEING MOVED BY ART": ANNEX

ANNEX II

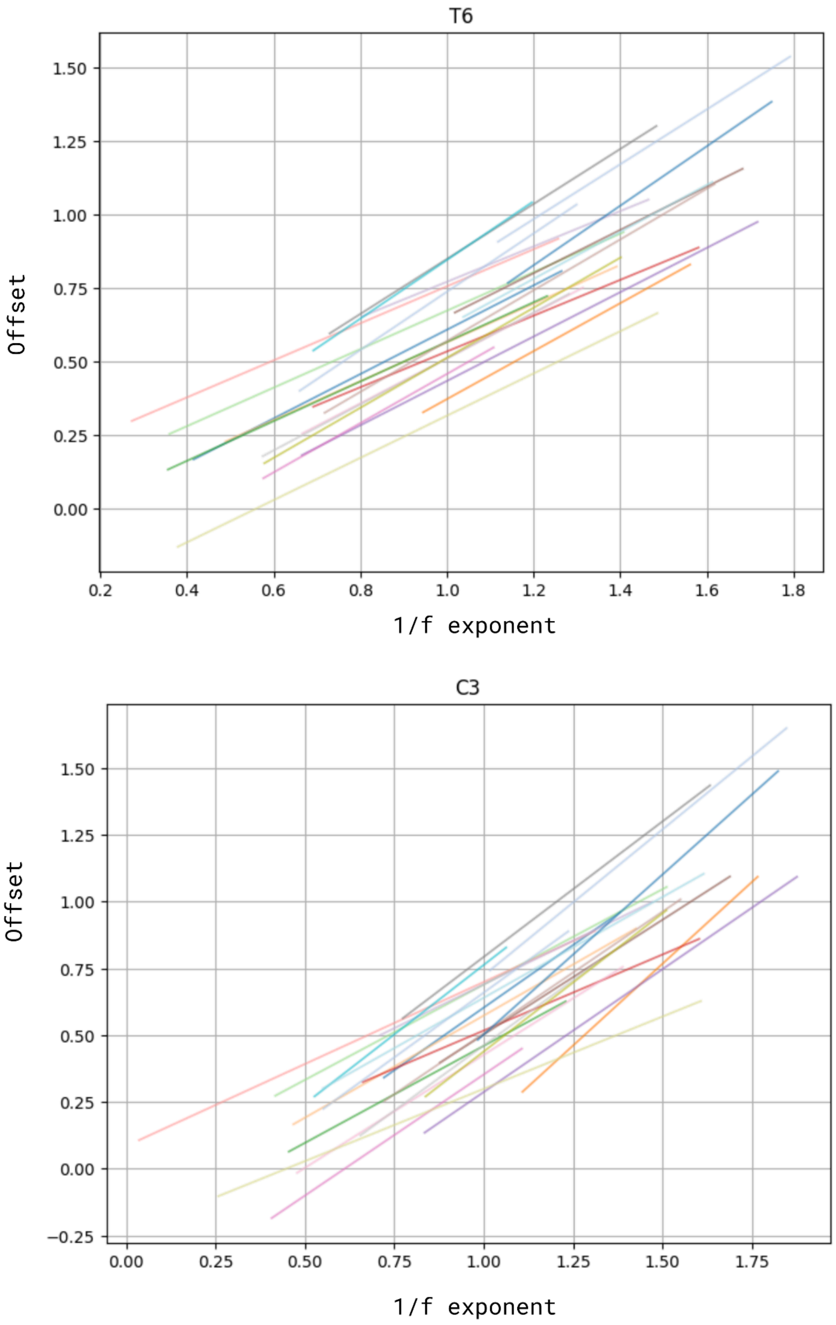


Figure 1A. Subject-level aperiodic offset–exponent coupling across ratings in T6 and C3. For each window (T6 upper, C3 lower), a line was fitted per subject relating the ROI-averaged aperiodic offset to the ROI-averaged 1/f exponent across rating-specific subject means. The plot shows inter-individual variability in the offset–exponent relationship within each window.

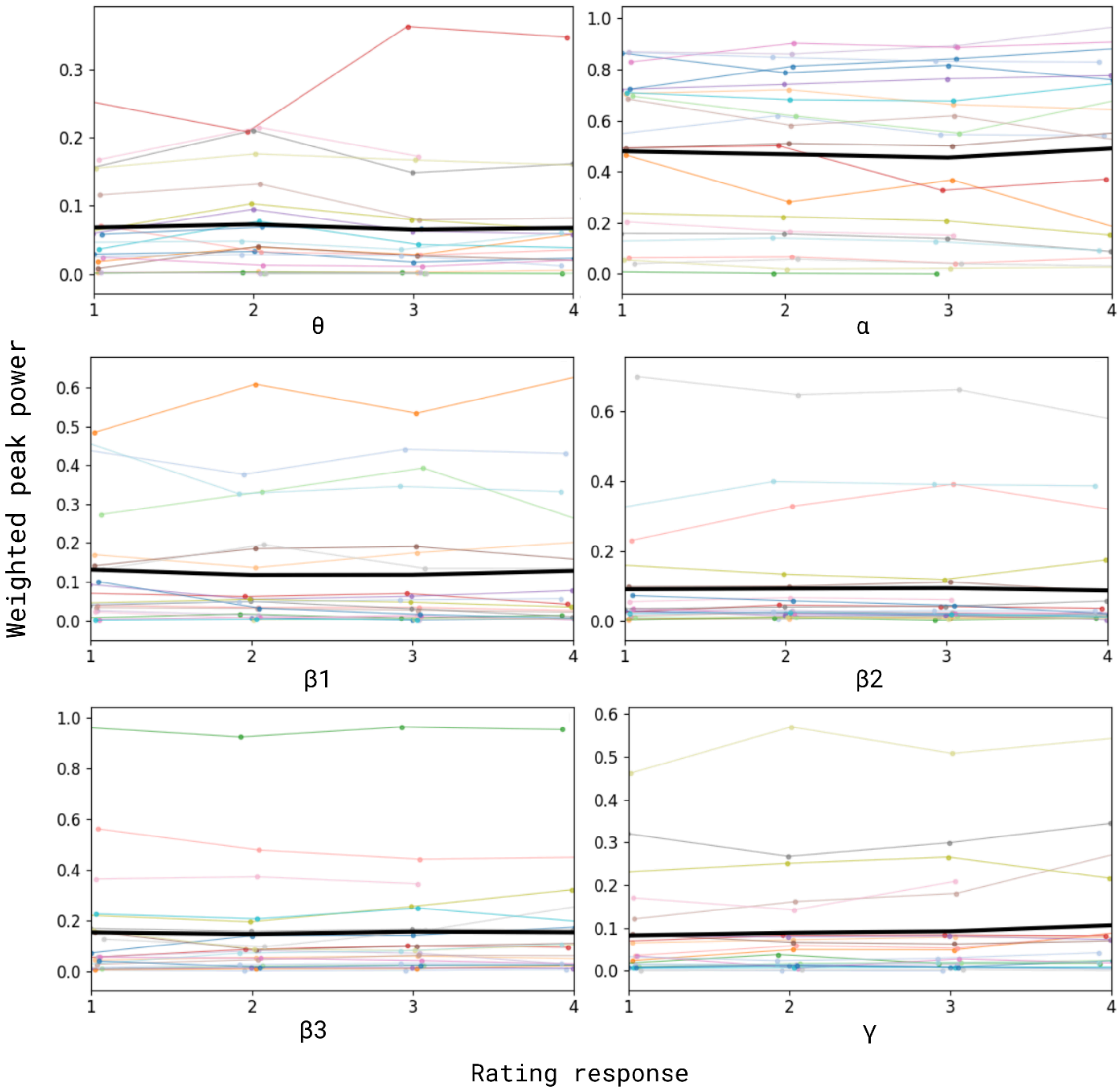


Figure 2A. Band-wise composition of periodic peak activity across ratings during T6. Spectral peaks were assigned to canonical bands (θ–γ) by centre frequency. Within each subject and rating, peak area (PW×BW) was summed per band and normalized by the subject's total PW×BW, yielding proportional (ROI-aggregated) band contributions. Thin lines show individuals; thick black lines indicate the group mean.

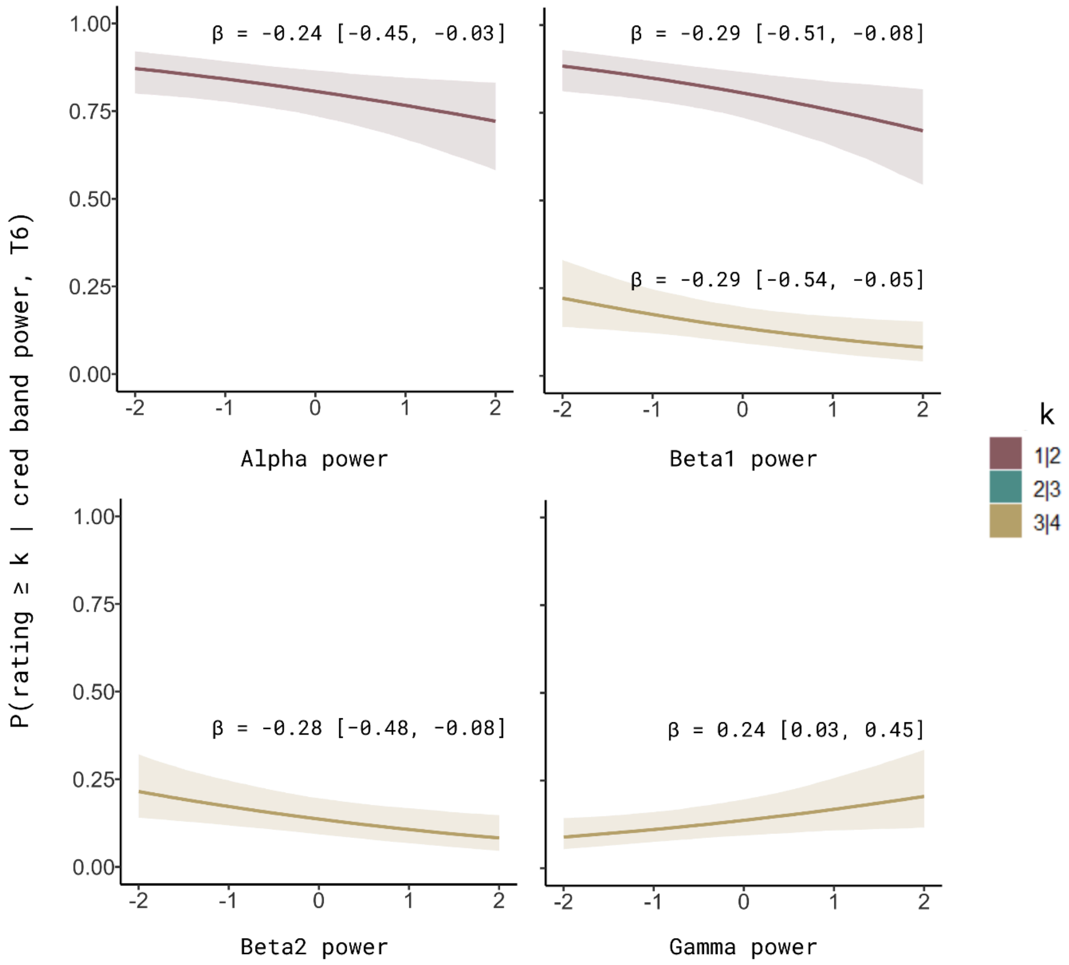

P(rating ≥ k | cred band power, T6)
β = -0.24 [-0.45, -0.03]
β = -0.29 [-0.51, -0.08]
β = -0.29 [-0.54, -0.05]
β = -0.28 [-0.48, -0.08]
β = 0.24 [0.03, 0.45]
1.00
0.75
0.50
0.25
0.00
-2
-1
0
1
2
Alpha power
Beta1 power
Beta2 power
Gamma power
k
1|2
2|3
3|4

Figure 3A. Population-level conditional effects of oscillatory band power during contemplation (T6). Posterior estimates show how z-scored power in canonical frequency bands (α, β1, β2, γ) modulates the probability of crossing selected cutpoints on the latent aesthetic response scale. Shaded areas indicate 95% credible intervals. Estimates are derived from a category-specific cumulative link mixed model (CSCLMM) fitted to ratings ($r_1$–$r_4$), including band-specific power predictors at T6 and random intercepts capturing inter-individual differences in baseline rating propensity.

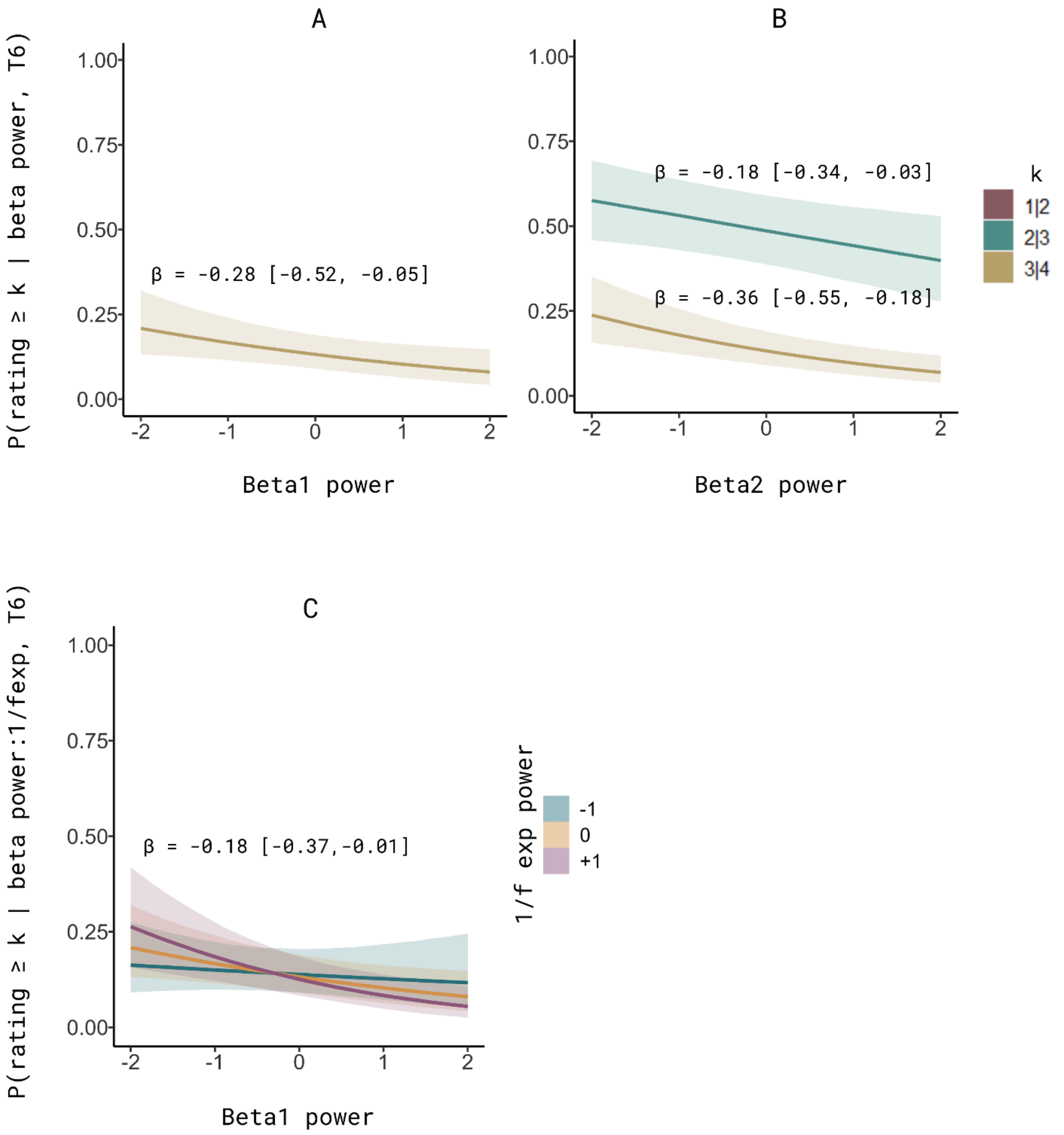


Figure 4A. Interactionist model for the contemplation window, T6 (model 9, without random effects). Graph A and B represent credible posterior estimates of β1 and β2 band power modulating the probability of crossing cutpoints selectively: β1 at 3|4 and β2 at 2|3 and 3|4. Graph C visualises the only credible interaction found. β1 and the aperiodic exponent interaction had a negative effect on the probability of crossing the highest cutpoint (3|4) on the latent scale. This interaction effect appears to be potentiated by the magnitude of the β1 power and the exponent; however, less aperiodic power seems to mitigate the negative effect. The shaded areas indicate the 95% credible intervals.

| ROI name | Electrodes | N channels |
|---|---|---|
| PF_left | FP1, AF7, AF3 | 3 |
| PF_mid | FPZ, AFZ | 2 |
| PF_right | FP2, AF8, AF4 | 3 |
| F_left | F7, F5, F3, F1 | 4 |
| F_mid | FZ | 1 |
| F_right | F8, F6, F4, F2 | 4 |
| FC_left | FC5, FC3, FC1 | 3 |
| FC_mid | FCZ | 1 |
| FC_right | FC6, FC4, FC2 | 3 |
| T_left | FT7, T7, TP7 | 3 |
| T_right | FT8, T8, TP8 | 3 |
| C_left | C5, C3, C1 | 3 |
| C_mid | CZ | 1 |
| C_right | C6, C4, C2 | 3 |
| CP_left | CP5, CP3, CP1 | 3 |
| CP_mid | CPZ | 1 |
| CP_right | CP6, CP4, CP2 | 3 |
| P_left | P7, P5, P3, P1 | 4 |
| P_mid | PZ | 1 |
| P_right | P8, P6, P4, P2 | 4 |
| PO_left | PO7, PO3 | 2 |

| | | |
|---|---|---|
| PO_mid | POZ | 1 |
| PO_right | PO8, PO4 | 2 |
| O_left | O1 | 1 |
| O_mid | OZ | 1 |
| O_right | O2 | 1 |

Table 1A. Region-of-interest (ROI) definitions based on electrodes of a 64-channel montage with the 10/20 coordinate system.

| Band | ICC power | CI95% | ICC center freq | CI95% | ICC bandwidth | CI95% |
|---|---|---|---|---|---|---|
| theta | 0.89 | [0.76, 0.95] | 0.84 | [0.65, 0.93] | 0.82 | [0.61, 0.92] |
| alpha | 0.88 | [0.74, 0.95] | 0.66 | [0.33, 0.84] | 0.80 | [0.57, 0.91] |
| beta1 | 0.86 | [0.7, 0.94] | 0.70 | [0.41, 0.87] | 0.88 | [0.73, 0.95] |
| beta2 | 0.83 | [0.64, 0.93] | 0.59 | [0.23, 0.81] | 0.78 | [0.54, 0.9] |
| beta3 | 0.95 | [0.89, 0.98] | 0.91 | [0.8, 0.96] | 0.89 | [0.75, 0.95] |
| gamma | 0.87 | [0.71, 0.94] | 0.97 | [0.92, 0.99] | 0.85 | [0.68, 0.94] |

Table 2A. Cross-window (T6/C3) reliability of periodic peak parameters across canonical frequency bands. Intraclass correlation coefficients (ICCs) were computed using subject as target and window (T6 vs. C3) as rater. CI95% = 95 % confidence interval.

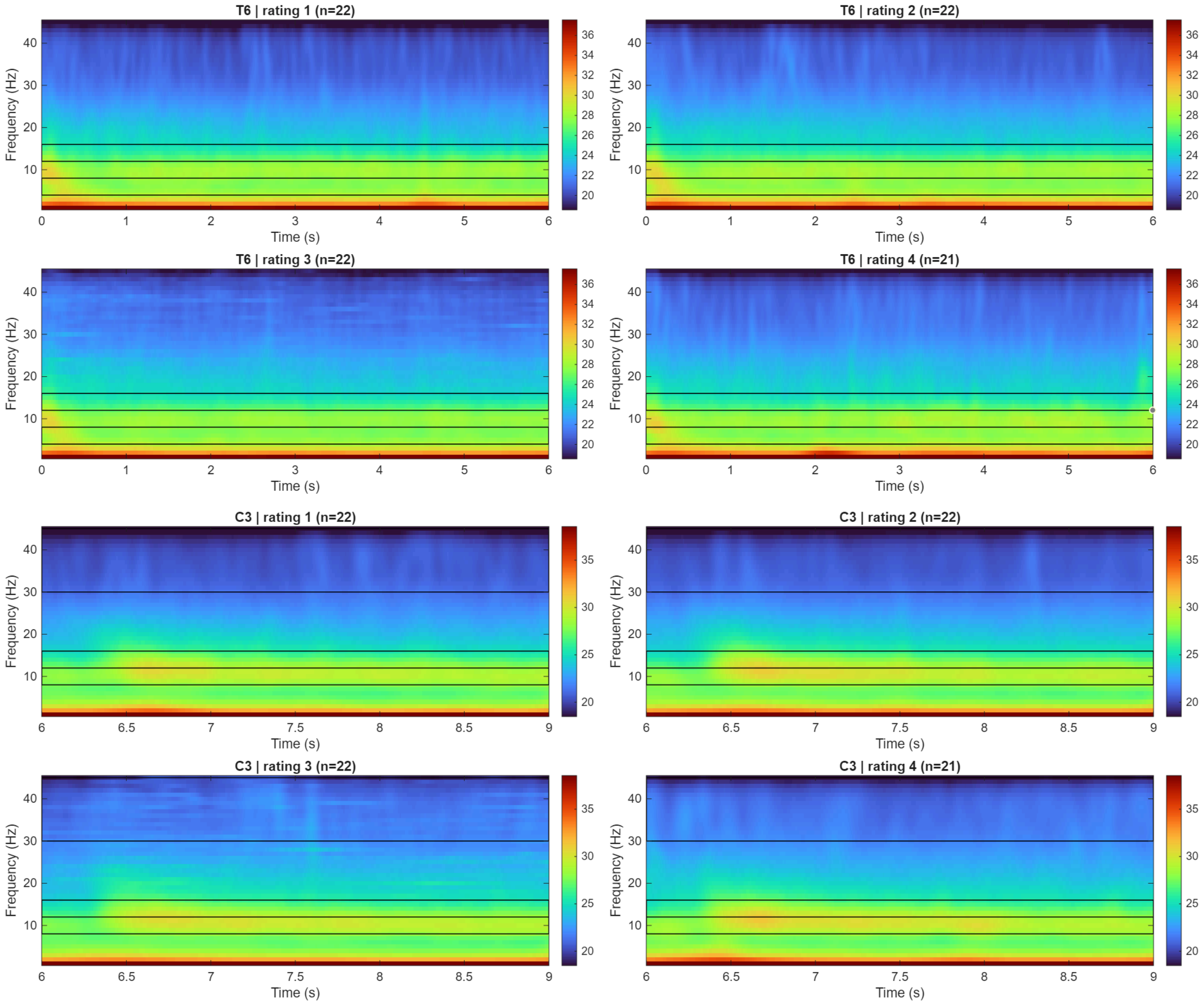


Figure 5A. Raw time–frequency representations for aesthetic ratings {1, 2, 3, 4} during T6 and C3 windows.

| Window | Contrast | Freq.band | Time bins | Sign | p value | Cluster mass | Channels |
|---|---|---|---|---|---|---|---|
| T6 | all vs. 4 | beta2 | 1000-1500 ms | pos | 0.004 | 21.2 | 8 |
| T6 | all vs. 4 | beta2 | 1500-2000 ms | pos | 0.005 | 23.2 | 8 |
| T6 | 1 vs. all | beta1 | 2000-2500 ms | pos | 0.012 | 18.5 | 7 |
| T6 | all vs. 4 | beta2 | 2000-2500 ms | pos | 0.013 | 19.3 | 8 |
| T6 | 1 vs. all | beta1 | 3000-3500 ms | pos | 0.015 | 18.6 | 7 |
| T6 | all vs. 4 | beta2 | 1000-1500 ms | pos | 0.016 | 12.8 | 5 |
| T6 | all vs. 4 | beta1 | 1000-1500 ms | pos | 0.018 | 21.2 | 9 |
| T6 | all vs. 4 | beta1 | 1000-1500 ms | pos | 0.04 | 10.2 | 4 |
| T6 | all vs. 4 | beta1 | 1500-2000 ms | pos | 0.044 | 9.3 | 4 |

Table 3A. Summary of significant clusters from the cluster-based permutation analysis. All significant clusters correspond to positive effects during the T6 window. Clusters are ordered by ascending p-value.

| ID model | Model class | Window/Data | R formula |
|---|---|---|---|
| 1 | Class 1/ Periodic | T6 | rating ~ cs(pw_delta) + cs(pw_theta) + cs(pw_alpha) + cs(pw_beta1) + cs(pw_beta2) + cs(pw_beta3) + cs(pw_gamma) + (1 \| subject) + (1 \| artworkn) |
| 2 | Class 1/ Periodic | C3 | rating ~ cs(pw_delta) + cs(pw_theta) + cs(pw_alpha) + cs(pw_beta1) + cs(pw_beta2) + cs(pw_beta3) + cs(pw_gamma) + (1 \| subject) + (1 \| artworkn) |
| 3 | Class 1/ Aperiodic | T6 | rating ~ cs(z_exponent) + cs(z_offset) + (1 \| subject) + (1 \| artworkn) |
| 4 | Class 1/ Aperiodic | C3 | rating ~ cs(z_exponent) + cs(z_offset) + (1 \| subject) + (1 \| artworkn) |
| 5 | Class 2/ Periodic + slopes | T6 | rating ~ cs(pw_alpha) + cs(pw_beta1) + cs(pw_beta2) + cs(pw_gamma) + (1 + pw_alpha + pw_beta1 + pw_beta2 + pw_gamma \|\| subject) + (1 \| artworkn) |
| 6 | Class 2/ Periodic + slopes | C3 | rating ~ cs(pw_alpha) + cs(pw_beta1) + cs(pw_beta2) + cs(pw_gamma) + (1 + pw_alpha + pw_beta1 + pw_beta2 + pw_gamma \|\| subject) + (1 \| artworkn) |
| 7 | Class 2/ Aperiodic exponent + slopes | T6 | rating ~ cs(z_exponent) + (1 + z_exponent \|\| subject) + (1 \| artworkn) |
| 8.a | Class 2/ Aperiodic + slopes | C3 | rating ~ cs(z_exponent) + cs(z_offset) + (1 + z_exponent \|\| subject) + (1 \| artworkn) |
| 8.b | Class 2/ Aperiodic exponent only + slopes | C3 | rating ~ cs(z_exponent) + (1 + z_exponent \|\| subject) + (1 \| artworkn) |

Table 4A. 12 CSCLMMs were fitted in complexity progression. The columns describe a number to identify each model, its class according to the definitions in the 'Method' section, the window of analysis and the R formulas.

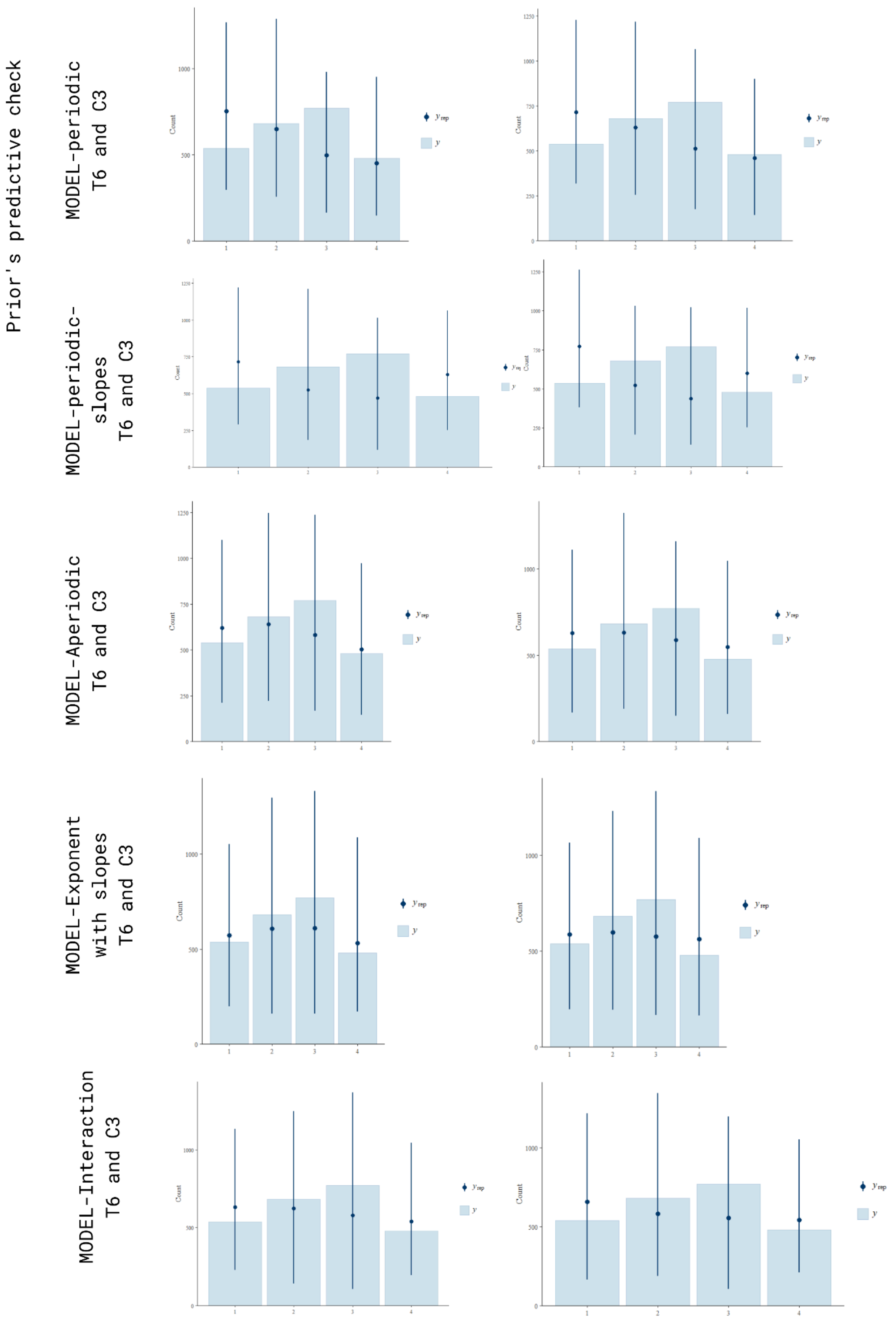


Figure 6A. Prior's predictive check for each model (rows), and windows (columns) T6 (left) and C3 (right). Simulated responses (dark blue dots with errors) based on the prior distributions compared to the observed response distribution (light bars). The X-axis shows ratings from 1 to 4, and the Y-axis shows the counts from 0 to 1000 or 1250 in steps of 250 or 500.

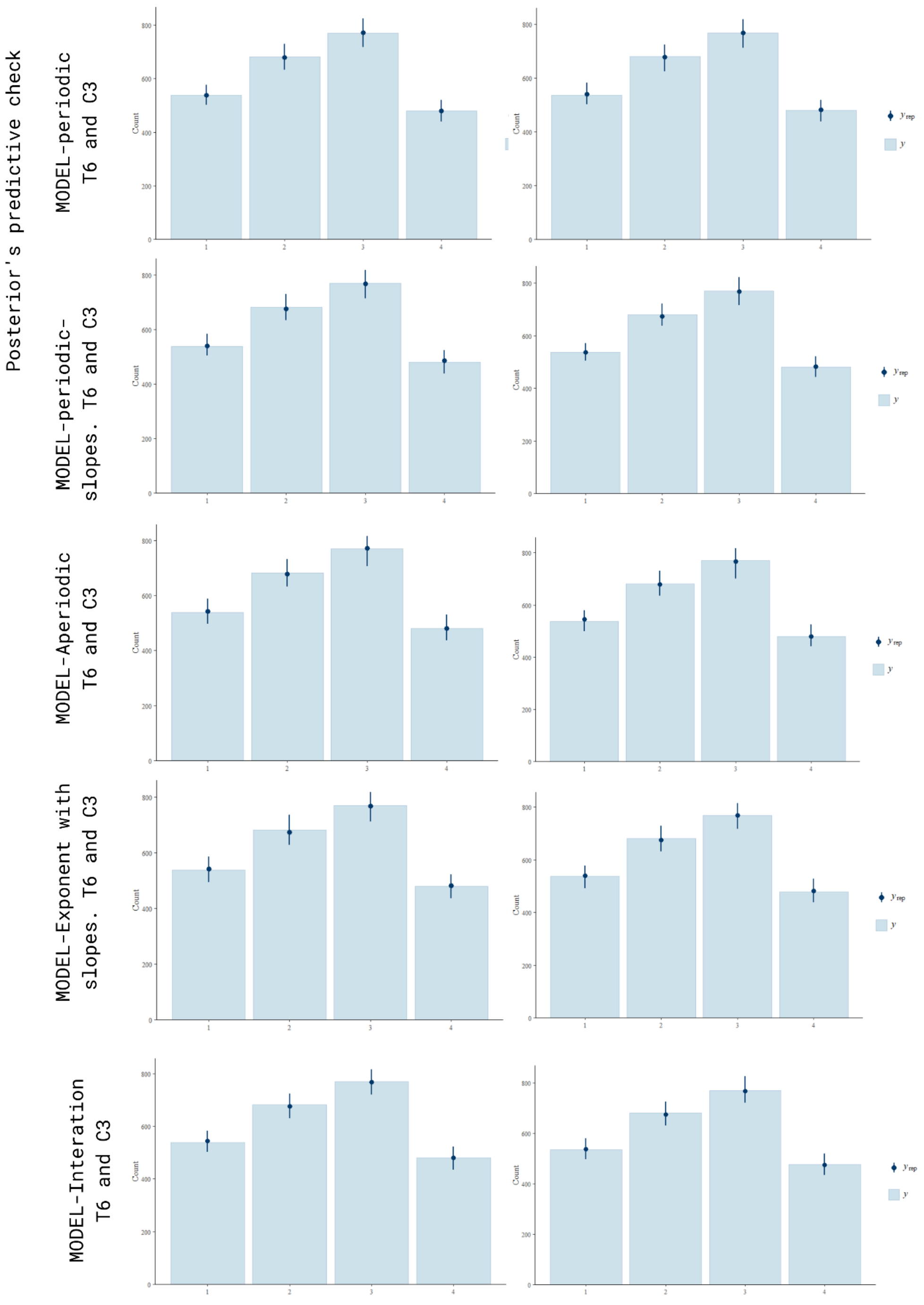


Figure 7A. Posterior's predictive check for each model (rows), and windows (columns) T6 (left) and C3 (right). Simulated responses (dark blue dots with errors) based on the posterior distributions compared to the observed response distribution (light bars). The X-axis shows ratings from 1 to 4, and the Y-axis shows counts from 0 to 800 in steps of 200.

| ID | Model | Win | N | Chains | Iter | Warmup | Draws | Divergence trans. | $\hat{R}$ max | ESS min (B/T) | Pareto k |
|---|---|---|---|---|---|---|---|---|---|---|---|
| 1 | Periodic, pop. | T6 | 2470 | 4 | 2000 | 1000 | 4000 | 15 (0.38%) | 1.01 | 814 / 1234 | < 0.7 |
| 2 | Periodic, pop. | C3 | 2465 | 4 | 2000 | 1000 | 4000 | 9 (0.23%) | 1.01 | 642 / 1212 | < 0.7 |
| 3 | Aperiodic, pop. | T6 | 2470 | 4 | 4000 | 2000 | 8000 | 0 | 1.00 | 1236 / 2272 | < 0.7 |
| 4 | Aperiodic, pop. | C3 | 2469 | 4 | 4000 | 2000 | 8000 | 0 | 1.00 | 1480 / 2773 | < 0.7 |
| 5 | Periodic + slopes | T6 | 2470 | 4 | 2000 | 1000 | 4000 | 0 | 1.01 | 720 / 510 | < 0.7 |
| 6 | Periodic + slopes | C3 | 2466 | 4 | 2000 | 1000 | 4000 | 0 | 1.01 | 779 / 1374 | < 0.7 |
| 7 | Aperiodic + slopes | T6 | 2470 | 4 | 4000 | 2000 | 8000 | 0 | 1.01 | 1195 / 1470 | < 0.7 |
| 8 | Aperiodic + Ap. Exp. slopes | C3 | 2469 | 4 | 4000 | 2000 | 8000 | 0 | 1.00 | 1964/ 3269 | < 0.7 |
| 8.b | Exponent only + Exp. slopes | C3 | 2469 | 4 | 4000 | 2000 | 8000 | 0 | 1.00 | 2388 / 3714 | < 0.7 |
| 9 | Interaction, pop. | T6 | 2470 | 4 | 4000 | 2000 | 8000 | 13 (0.16%) | 1.00 | 1339 / 2406 | < 0.7 |
| 10 | Interaction, pop. | C3 | 2465 | 4 | 4000 | 2000 | 8000 | 183 (2.3%) | 1.00 | 1892 / 3060 | < 0.7 |
| 11 | Interaction + slopes | T6 | 2470 | 4 | 4000 | 2000 | 8000 | 10 (0.13%) | 1.01 | 1615 / 1774 | < 0.7 |

Table 5A. Sampling specification and convergence diagnostics for all fitted models. Win. = time window; Iter = total iterations per chain; Draws = post-warmup draws (chains × [iter − warmup]); Divergent trans. = number of divergent transitions after warmup (percentage of total draws); $\hat{R}$ max = maximum R-hat across all parameters; ESS min (B/T) = minimum bulk / tail effective sample size; Pareto k = maximum Pareto-k diagnostic from LOO-CV. All models used the cmdstanr backend with overdispersed initial values and seed = 123.

§ Note on Table 5A and Figures 8A-12A:

Models 1, 2, 9, 10 and 11 produced divergent transitions. In all cases except for model 10, divergences were sparse, not concentrated along specific parameter pairs (no funnel geometry), and posterior summaries remained stable across refits with increased adapt_delta (≥ 0.999). Model 5 (T6) showed a minimum Tail ESS of 510 for sd(pw_gamma); all other parameters exceeded 700. Convergence and sampling diagnostics for models that produced divergent transitions are visualised in the subsequent Figures 8A-12A. Category-specific effects in cumulative models are labelled experimental in brms, which can increase geometric sampling difficulty.

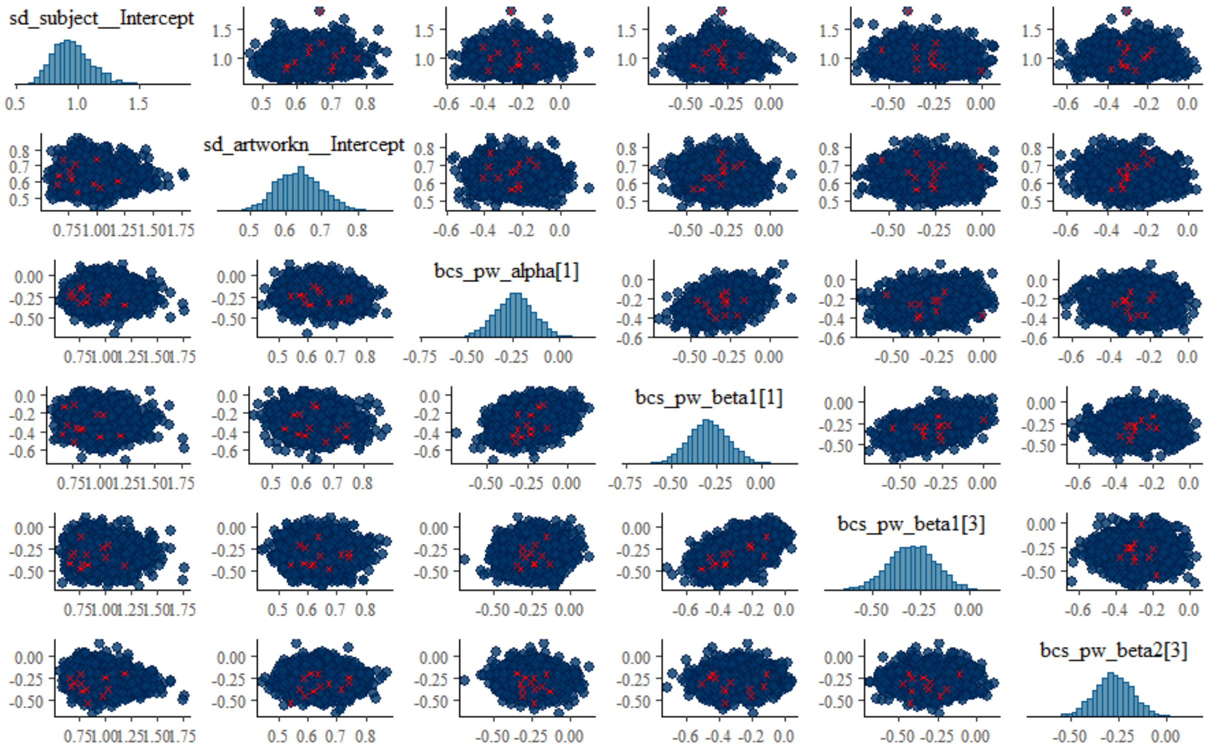


Figure 8A. Convergence and sampling diagnostics for Model 1, T6. Pairwise posterior draws for selected fixed effects and variance components are shown; divergent transitions are highlighted in red. The model produced 15 divergent transitions after warm-up (0.375% of post-warm-up draws, 15/4000).

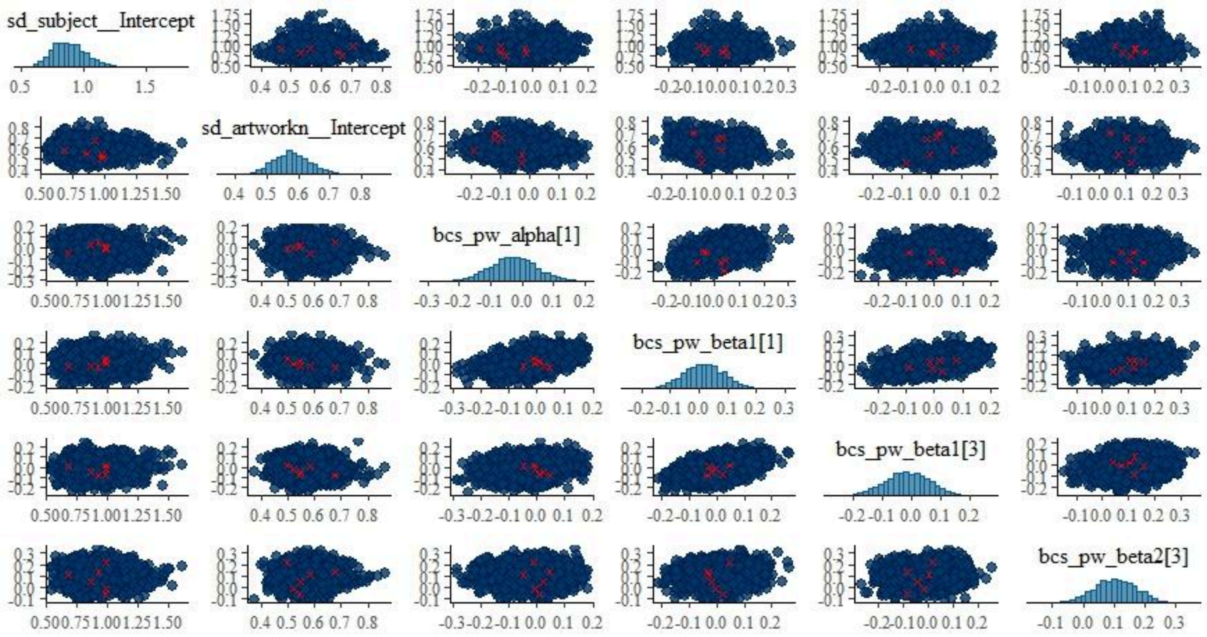


Figure 9A. Pairwise posterior distributions for key parameters of Model 2 (C3 window). Pairwise posterior draws for selected fixed effects and variance components are shown; divergent transitions are highlighted in red. The model produced 9 divergent transitions after warm-up (0.23% of post-warm-up draws).

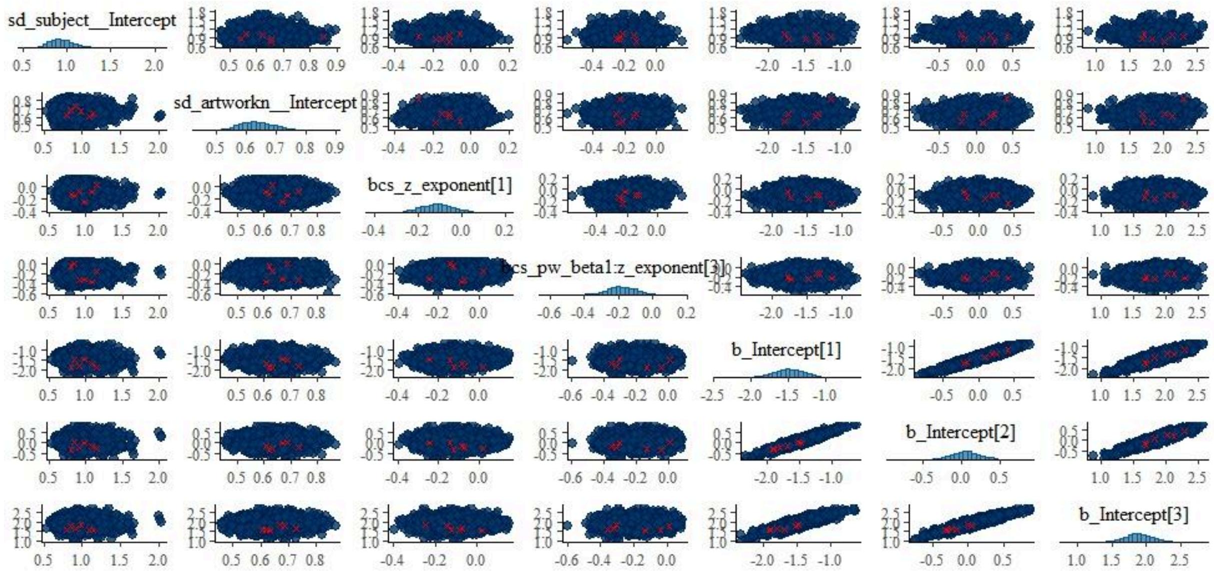


Figure 10A. Pairwise posterior distributions for key parameters of Model 9 (T6 window). Diagonal panels show marginal posterior densities, while off-diagonal panels show joint posterior samples. The model produced 13 divergent transitions after warm-up (0.16% of post-warm-up draws).

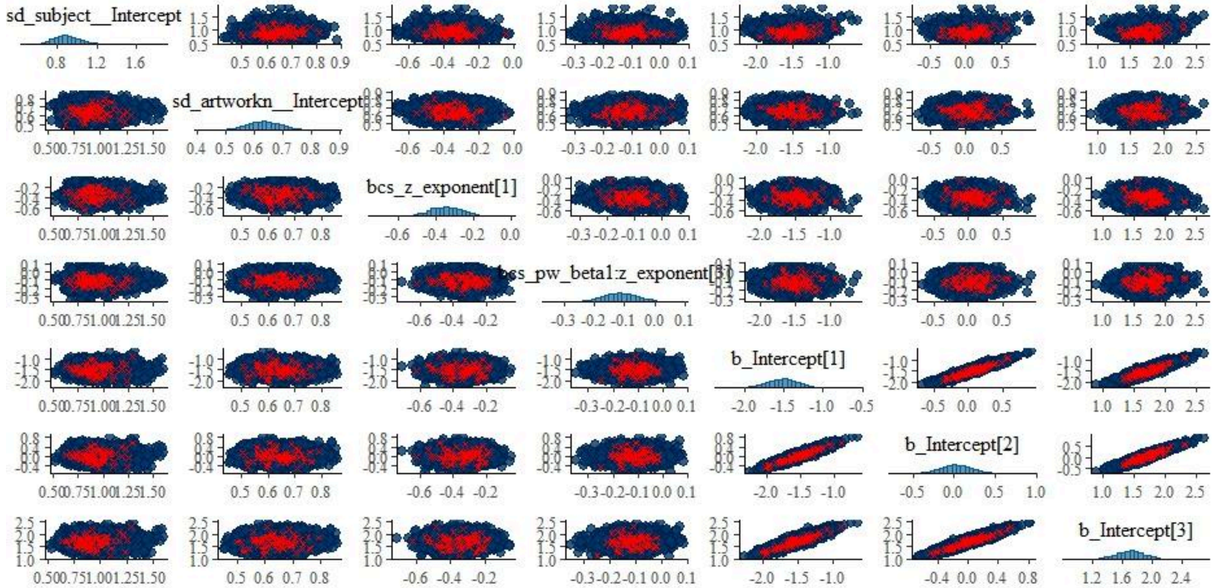


Figure 11A. Pairwise posterior distributions for key parameters of Model 10 (C3 window). The model produced 183 divergent transitions after warm-up (2.3% of post-warm-up draws).

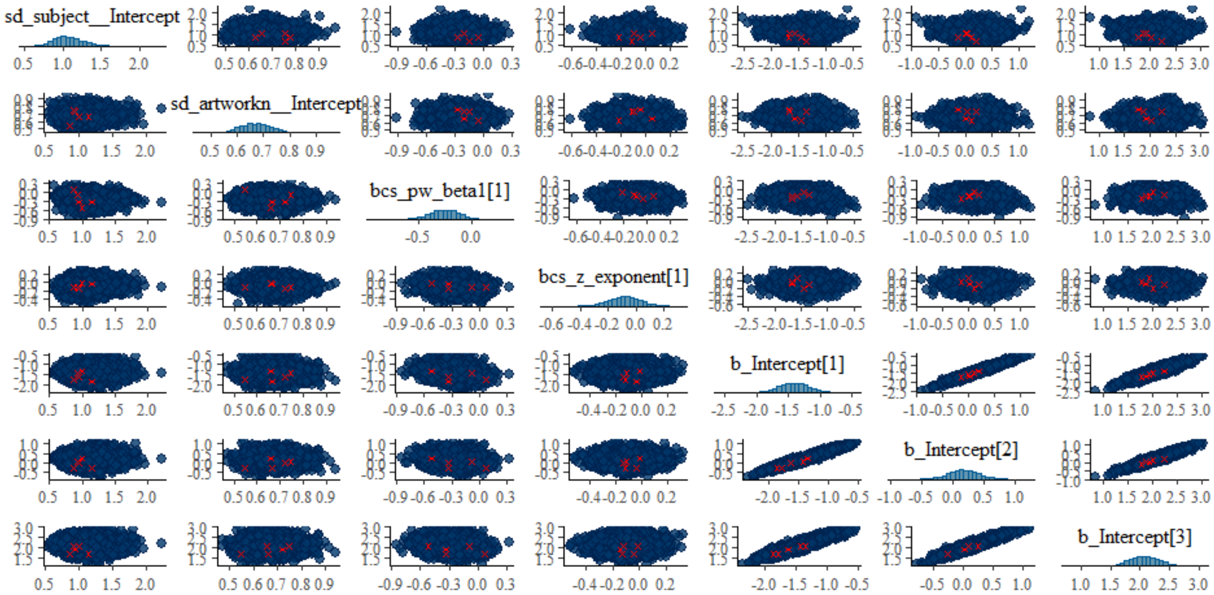


Figure 12A. Pairwise posterior distributions for key parameters of Model 11 (T6 window). The model produced 10 divergent transitions after warm-up (0.13% of post-warm-up draws).

MODEL 1. CSCLMM-periodic /T6

```
Family: cumulative
  Links: mu = logit; disc = identity
Formula: rating ~ cs(pw_delta) + cs(pw_theta) + cs(pw_alpha) + cs(pw_beta1) +
cs(pw_beta2) + cs(pw_beta3) + cs(pw_gamma) + (1 | subject) + (1 | artworkn)
   Data: per_t6 (Number of observations: 2470)
  Draws: 4 chains, each with iter = 2000; warmup = 1000; thin = 1;
         total post-warmup draws = 4000
```

```
Multilevel Hyperparameters:
~artworkn (Number of levels: 113)
              Estimate Est.Error l-95% CI u-95% CI Rhat Bulk_ESS Tail_ESS
sd(Intercept)     0.64      0.06     0.52     0.76 1.00     1841     2651

~subject (Number of levels: 22)
              Estimate Est.Error l-95% CI u-95% CI Rhat Bulk_ESS Tail_ESS
sd(Intercept)     0.96      0.16     0.69     1.32 1.00     1055     1568

Regression Coefficients:
             Estimate Est.Error l-95% CI u-95% CI Rhat Bulk_ESS Tail_ESS
Intercept[1]    -1.43      0.22    -1.87    -1.03 1.01      814     1493
Intercept[2]     0.00      0.21    -0.41     0.39 1.01      824     1473
Intercept[3]     1.84      0.22     1.41     2.28 1.01      891     1361
pw_delta[1]      0.07      0.07    -0.08     0.21 1.00     4247     3217
pw_delta[2]      0.07      0.06    -0.04     0.20 1.00     4157     3271
pw_delta[3]      0.00      0.07    -0.13     0.12 1.00     4303     3141
pw_theta[1]     -0.05      0.07    -0.18     0.09 1.00     3425     3213
pw_theta[2]     -0.03      0.06    -0.15     0.08 1.00     2811     2893
pw_theta[3]     -0.01      0.07    -0.15     0.13 1.00     3641     2867
pw_alpha[1]     -0.24      0.10    -0.44    -0.03 1.00     2487     3170
pw_alpha[2]     -0.12      0.10    -0.31     0.07 1.00     2333     2776
pw_alpha[3]     -0.14      0.11    -0.36     0.08 1.00     2623     2842
pw_beta1[1]     -0.29      0.11    -0.51    -0.07 1.00     2272     3272
pw_beta1[2]     -0.16      0.10    -0.36     0.04 1.00     1919     2954
pw_beta1[3]     -0.29      0.12    -0.53    -0.05 1.00     2170     2978
pw_beta2[1]     -0.09      0.09    -0.26     0.09 1.00     2514     3216
pw_beta2[2]     -0.13      0.08    -0.29     0.04 1.00     2362     3293
pw_beta2[3]     -0.28      0.10    -0.48    -0.08 1.00     2862     3619
pw_beta3[1]     -0.05      0.09    -0.23     0.14 1.00     2373     2959
pw_beta3[2]      0.10      0.09    -0.07     0.27 1.00     2341     2710
pw_beta3[3]      0.09      0.10    -0.11     0.29 1.00     2914     3114
pw_gamma[1]      0.06      0.10    -0.14     0.26 1.00     3016     3001
pw_gamma[2]      0.13      0.09    -0.04     0.31 1.00     3077     2758
pw_gamma[3]      0.24      0.11     0.02     0.45 1.00     3437     3147
Further Distributional Parameters:
```

```
     Estimate Est.Error l-95% CI u-95% CI Rhat Bulk_ESS Tail_ESS
disc     1.00      0.00     1.00     1.00   NA       NA       NA
Draws were sampled using sample(hmc). For each parameter, Bulk_ESS
and Tail_ESS are effective sample size measures, and Rhat is the potential
scale reduction factor on split chains (at convergence, Rhat = 1).
```

Table 6A. Output summary for: Model 1. CSCLMM-periodic /T6

MODEL 2. CSCLMM-periodic /C3

```
Family: cumulative
  Links: mu = logit; disc = identity
Formula: rating ~ cs(pw_delta) + cs(pw_theta) + cs(pw_alpha) + cs(pw_beta1) +
cs(pw_beta2) + cs(pw_beta3) + cs(pw_gamma) + (1 | subject) + (1 | artworkn)
   Data: per_c3 (Number of observations: 2466)
  Draws: 4 chains, each with iter = 2000; warmup = 1000; thin = 1;
         total post-warmup draws = 4000
```

```
Multilevel Hyperparameters:
~artworkn (Number of levels: 113)
              Estimate Est.Error l-95% CI u-95% CI Rhat Bulk_ESS Tail_ESS
sd(Intercept)     0.58      0.06     0.47     0.70 1.00     1355     2170
~subject (Number of levels: 22)
              Estimate Est.Error l-95% CI u-95% CI Rhat Bulk_ESS Tail_ESS
sd(Intercept)     0.90      0.15     0.65     1.23 1.00      826     1488
Regression Coefficients:
             Estimate Est.Error l-95% CI u-95% CI Rhat Bulk_ESS Tail_ESS
Intercept[1]    -1.54      0.21    -1.95    -1.13 1.01      672     1234
Intercept[2]     0.01      0.20    -0.40     0.41 1.01      642     1212
Intercept[3]     1.69      0.21     1.28     2.10 1.01      704     1240
pw_delta[1]      0.08      0.07    -0.06     0.22 1.00     3142     2888
pw_delta[2]     -0.01      0.06    -0.12     0.11 1.00     2366     2740
pw_delta[3]      0.03      0.06    -0.10     0.15 1.00     2698     2526
pw_theta[1]     -0.05      0.06    -0.16     0.07 1.00     2562     2730
pw_theta[2]      0.00      0.05    -0.10     0.11 1.00     2341     2853
pw_theta[3]      0.06      0.06    -0.06     0.19 1.00     2742     2895
pw_alpha[1]     -0.03      0.08    -0.18     0.12 1.00     1887     2894
pw_alpha[2]     -0.02      0.07    -0.16     0.11 1.00     1924     2511
pw_alpha[3]     -0.12      0.08    -0.28     0.04 1.00     2024     2811
pw_beta1[1]      0.02      0.07    -0.12     0.15 1.00     1716     2252
pw_beta1[2]      0.04      0.06    -0.09     0.17 1.00     1573     2203
pw_beta1[3]     -0.01      0.08    -0.16     0.13 1.00     1887     2511
pw_beta2[1]      0.10      0.07    -0.04     0.23 1.00     2509     2674
pw_beta2[2]      0.07      0.06    -0.05     0.19 1.00     1963     2497
pw_beta2[3]      0.10      0.07    -0.04     0.24 1.00     2209     2540
pw_beta3[1]     -0.12      0.07    -0.26     0.03 1.00     1823     2521
pw_beta3[2]     -0.04      0.07    -0.18     0.08 1.00     1703     2383
```

```
pw_beta3[3]     -0.11      0.08    -0.27     0.04 1.00     2026     2529
pw_gamma[1]      0.04      0.09    -0.12     0.22 1.00     2447     2760
pw_gamma[2]      0.08      0.07    -0.06     0.22 1.00     2156     2953
pw_gamma[3]      0.08      0.09    -0.08     0.25 1.00     2075     2402
```

Table 7A. Output summary for: Model 2. CSCLMM-periodic /C3

MODEL 3. CSCLMM-aperiodic/T6

```
Family: cumulative.
  Links: mu = logit; disc = identity
Formula: rating ~ cs(z_exponent) + cs(z_offset) + (1 | subject) + (1 | artworkn)
   Data: df_t6 (Number of observations: 2470)
  Draws: 4 chains, each with iter = 4000; warmup = 2000; thin = 1;
         total post-warmup draws = 8000
```

```
Multilevel Hyperparameters:
~artworkn (Number of levels: 113)
              Estimate Est.Error l-95% CI u-95% CI Rhat Bulk_ESS Tail_ESS
sd(Intercept)     0.63      0.06     0.52     0.76 1.00     3059     4731
~subject (Number of levels: 22)
              Estimate Est.Error l-95% CI u-95% CI Rhat Bulk_ESS Tail_ESS
sd(Intercept)     0.92      0.15     0.67     1.26 1.00     2239     3862
Regression Coefficients:
              Estimate Est.Error l-95% CI u-95% CI Rhat Bulk_ESS Tail_ESS
Intercept[1]     -1.55      0.21    -1.96    -1.14 1.00     1236     2272
Intercept[2]     -0.02      0.20    -0.42     0.39 1.00     1235     2394
Intercept[3]      1.71      0.21     1.31     2.12 1.00     1284     2640
z_exponent[1]    -0.24      0.15    -0.53     0.05 1.00     2754     4679
z_exponent[2]    -0.08      0.14    -0.35     0.19 1.00     2479     4184
z_exponent[3]    -0.03      0.15    -0.32     0.25 1.00     2810     4561
z_offset[1]       0.13      0.16    -0.18     0.44 1.00     2393     4085
z_offset[2]       0.07      0.15    -0.24     0.37 1.00     2247     3703
z_offset[3]      -0.03      0.16    -0.36     0.29 1.00     2512     4153
```

Table 8A. Output summary for: Model 3. CSCLMM-aperiodic/T6

MODEL 4. CSCLMM-aperiodic/C3

```
Family: cumulative
  Links: mu = logit; disc = identity
Formula: rating ~ cs(z_exponent) + cs(z_offset) + (1 | subject) + (1 | artworkn)
   Data: df_c3 (Number of observations: 2469)
  Draws: 4 chains, each with iter = 4000; warmup = 2000; thin = 1;
         total post-warmup draws = 8000
```

```
Multilevel Hyperparameters:
~artworkn (Number of levels: 113)
              Estimate Est.Error l-95% CI u-95% CI Rhat Bulk_ESS Tail_ESS
sd(Intercept)     0.63      0.06     0.52     0.76 1.00     3507     5557
~subject (Number of levels: 22)
              Estimate Est.Error l-95% CI u-95% CI Rhat Bulk_ESS Tail_ESS
sd(Intercept)     0.92      0.15     0.67     1.26 1.00     2095     3633
Regression Coefficients:
              Estimate Est.Error l-95% CI u-95% CI Rhat Bulk_ESS Tail_ESS
Intercept[1]     -1.55      0.20    -1.95    -1.16 1.00     1506     2856
Intercept[2]     -0.02      0.20    -0.41     0.38 1.00     1480     2958
Intercept[3]      1.73      0.20     1.32     2.12 1.00     1573     2773
z_exponent[1]    -0.43      0.14    -0.71    -0.15 1.00     3460     5277
z_exponent[2]    -0.29      0.13    -0.55    -0.05 1.00     2959     4648
z_exponent[3]    -0.25      0.14    -0.53     0.00 1.00     3406     5000
z_offset[1]       0.18      0.15    -0.12     0.48 1.00     3080     5201
z_offset[2]       0.16      0.14    -0.12     0.44 1.00     2775     4543
z_offset[3]       0.07      0.15    -0.24     0.37 1.00     3169     4771
```

Table 9A. Output summary for: Model 4. CSCLMM-aperiodic/C3

MODEL 5. CSCLMM-periodic-with slopes /T6

```
Family: cumulative
  Links: mu = logit; disc = identity
Formula: rating ~ cs(pw_alpha) + cs(pw_beta1) + cs(pw_beta2) + cs(pw_gamma) + (1 +
pw_alpha + pw_beta1 + pw_beta2 + pw_gamma || subject) + (1 | artworkn)
   Data: per_t6 (Number of observations: 2470)
  Draws: 4 chains, each with iter = 2000; warmup = 1000; thin = 1;
         total post-warmup draws = 4000
```

```
Multilevel Hyperparameters:
~artworkn (Number of levels: 113)
              Estimate Est.Error l-95% CI u-95% CI Rhat Bulk_ESS Tail_ESS
sd(Intercept)     0.66      0.06     0.54     0.79 1.00     1621     2281
~subject (Number of levels: 22)
              Estimate Est.Error l-95% CI u-95% CI Rhat Bulk_ESS Tail_ESS
sd(Intercept)     1.06      0.20     0.73     1.49 1.01     1190     2148
```

| | Estimate | Est.Error | l-95% CI | u-95% CI | Rhat | Bulk_ESS | Tail_ESS |
|---|---|---|---|---|---|---|---|
| sd(pw_alpha) | 0.61 | 0.15 | 0.35 | 0.94 | 1.00 | 1538 | 2069 |
| sd(pw_beta1) | 0.26 | 0.16 | 0.01 | 0.61 | 1.00 | 779 | 1408 |
| sd(pw_beta2) | 0.29 | 0.14 | 0.04 | 0.57 | 1.00 | 746 | 725 |
| sd(pw_gamma) | 0.29 | 0.14 | 0.03 | 0.58 | 1.01 | 720 | 510 |

Regression Coefficients:

| | Estimate | Est.Error | l-95% CI | u-95% CI | Rhat | Bulk_ESS | Tail_ESS |
|---|---|---|---|---|---|---|---|
| Intercept[1] | -1.37 | 0.25 | -1.84 | -0.89 | 1.00 | 1007 | 1392 |
| Intercept[2] | 0.18 | 0.24 | -0.28 | 0.65 | 1.00 | 1070 | 1809 |
| Intercept[3] | 2.05 | 0.25 | 1.56 | 2.54 | 1.00 | 1100 | 1561 |
| pw_alpha[1] | -0.29 | 0.17 | -0.62 | 0.04 | 1.00 | 1091 | 1873 |
| pw_alpha[2] | -0.26 | 0.17 | -0.59 | 0.07 | 1.00 | 1052 | 1932 |
| pw_alpha[3] | -0.26 | 0.17 | -0.61 | 0.07 | 1.00 | 1164 | 2020 |
| pw_beta1[1] | -0.31 | 0.13 | -0.58 | -0.05 | 1.00 | 1803 | 2030 |
| pw_beta1[2] | -0.20 | 0.13 | -0.46 | 0.04 | 1.00 | 1502 | 2178 |
| pw_beta1[3] | -0.32 | 0.14 | -0.60 | -0.05 | 1.00 | 1821 | 2267 |
| pw_beta2[1] | -0.07 | 0.12 | -0.30 | 0.18 | 1.00 | 1338 | 1838 |
| pw_beta2[2] | -0.13 | 0.12 | -0.35 | 0.11 | 1.01 | 1067 | 1639 |
| pw_beta2[3] | -0.28 | 0.13 | -0.53 | -0.01 | 1.00 | 1231 | 1622 |
| pw_gamma[1] | 0.14 | 0.12 | -0.09 | 0.37 | 1.00 | 1741 | 2239 |
| pw_gamma[2] | 0.11 | 0.11 | -0.10 | 0.33 | 1.00 | 1297 | 1955 |
| pw_gamma[3] | 0.20 | 0.12 | -0.04 | 0.43 | 1.00 | 1436 | 2061 |

Table 10A. Output summary for: Model 5. CSCLMM-periodic-with slopes /T6

MODEL 6. CSCLMM-periodic-with slopes /C3

```
Family: cumulative
  Links: mu = logit; disc = identity
Formula: rating ~ cs(pw_alpha) + cs(pw_beta1) + cs(pw_beta2) + cs(pw_gamma) + (1 +
pw_alpha + pw_beta1 + pw_beta2 + pw_gamma || subject) + (1 | artworkn)
   Data: per_c3 (Number of observations: 2466)
  Draws: 4 chains, each with iter = 2000; warmup = 1000; thin = 1;
         total post-warmup draws = 4000
```

Multilevel Hyperparameters:

~artworkn (Number of levels: 113)

| | Estimate | Est.Error | l-95% CI | u-95% CI | Rhat | Bulk_ESS | Tail_ESS |
|---|---|---|---|---|---|---|---|
| sd(Intercept) | 0.58 | 0.06 | 0.47 | 0.72 | 1.00 | 1820 | 2316 |

~subject (Number of levels: 22)

| | Estimate | Est.Error | l-95% CI | u-95% CI | Rhat | Bulk_ESS | Tail_ESS |
|---|---|---|---|---|---|---|---|
| sd(Intercept) | 0.94 | 0.16 | 0.67 | 1.31 | 1.00 | 1180 | 1608 |
| sd(pw_alpha) | 0.37 | 0.10 | 0.21 | 0.59 | 1.00 | 1714 | 2401 |
| sd(pw_beta1) | 0.11 | 0.07 | 0.01 | 0.27 | 1.00 | 1318 | 2202 |
| sd(pw_beta2) | 0.08 | 0.06 | 0.00 | 0.23 | 1.00 | 2057 | 1785 |
| sd(pw_gamma) | 0.11 | 0.08 | 0.00 | 0.30 | 1.00 | 1312 | 1513 |

Regression Coefficients:

```
             Estimate Est.Error l-95% CI u-95% CI Rhat Bulk_ESS Tail_ESS
Intercept[1]    -1.44      0.21    -1.87    -1.03 1.00      781     1581
Intercept[2]     0.08      0.21    -0.34     0.48 1.00      779     1374
Intercept[3]     1.80      0.21     1.38     2.22 1.00      813     1670
pw_alpha[1]      0.01      0.11    -0.20     0.23 1.01     1184     1575
pw_alpha[2]     -0.04      0.10    -0.24     0.17 1.01     1111     1638
pw_alpha[3]     -0.15      0.11    -0.38     0.06 1.01     1096     1722
pw_beta1[1]      0.03      0.08    -0.13     0.18 1.00     1656     2211
pw_beta1[2]      0.05      0.07    -0.10     0.19 1.00     1588     2342
pw_beta1[3]     -0.00      0.08    -0.16     0.15 1.00     1812     2291
pw_beta2[1]      0.08      0.07    -0.07     0.22 1.00     2261     2608
pw_beta2[2]      0.06      0.07    -0.07     0.19 1.00     2089     2673
pw_beta2[3]      0.09      0.07    -0.05     0.23 1.00     2646     2744
pw_gamma[1]      0.09      0.09    -0.07     0.26 1.00     2378     2680
pw_gamma[2]      0.06      0.08    -0.09     0.21 1.00     1885     2608
pw_gamma[3]      0.10      0.08    -0.06     0.27 1.00     2168     2793
```

Table 11A. Output summary for: Model 6. CSCLMM-periodic-with slopes /C3

MODEL 7. CSCLMM-exponent-with slopes/T6

```
Family: cumulative
  Links: mu = logit; disc = identity
Formula: rating ~ cs(z_exponent) + (1 + z_exponent || subject) + (1 | artworkn)
   Data: df_t6 (Number of observations: 2470)
  Draws: 4 chains, each with iter = 4000; warmup = 2000; thin = 1;
         total post-warmup draws = 8000
```

```
Multilevel Hyperparameters:
~artworkn (Number of levels: 113)
              Estimate Est.Error l-95% CI u-95% CI Rhat Bulk_ESS Tail_ESS
sd(Intercept)     0.63      0.06     0.52     0.76 1.00     2550     4553
~subject (Number of levels: 22)
               Estimate Est.Error l-95% CI u-95% CI Rhat Bulk_ESS Tail_ESS
sd(Intercept)      0.93      0.16     0.68     1.30 1.00     1766     3064
sd(z_exponent)     0.30      0.14     0.04     0.59 1.01     1195     1470
Regression Coefficients:
              Estimate Est.Error l-95% CI u-95% CI Rhat Bulk_ESS Tail_ESS
Intercept[1]     -1.54      0.20    -1.94    -1.15 1.00     1218     2302
Intercept[2]     -0.01      0.20    -0.40     0.38 1.00     1184     2173
Intercept[3]      1.73      0.20     1.33     2.13 1.00     1237     2279
z_exponent[1]    -0.17      0.11    -0.39     0.05 1.00     2115     2770
z_exponent[2]    -0.06      0.11    -0.28     0.15 1.00     2003     2448
z_exponent[3]    -0.09      0.11    -0.32     0.12 1.00     2194     2613
```

Table 12A. Output summary for: Model 7. CSCLMM-exponent-with slopes/T6

MODEL 8.a CSCLMM-APERIODIC-with slopes/C3

```
Family: cumulative
  Links: mu = logit; disc = identity
Formula: rating ~ cs(z_exponent) + cs(z_offset) + (1 + z_exponent || subject) + (1 |
artworkn)
   Data: df_c3 (Number of observations: 2469)
  Draws: 4 chains, each with iter = 4000; warmup = 2000; thin = 1;
         total post-warmup draws = 8000
```

```
Multilevel Hyperparameters:
~artworkn (Number of levels: 113)
              Estimate Est.Error l-95% CI u-95% CI Rhat Bulk_ESS Tail_ESS
sd(Intercept)     0.64      0.06     0.53     0.77 1.00     3306     5220

~subject (Number of levels: 22)
               Estimate Est.Error l-95% CI u-95% CI Rhat Bulk_ESS Tail_ESS
sd(Intercept)      0.92      0.16     0.66     1.27 1.00     2866     4445
sd(z_exponent)     0.48      0.14     0.25     0.78 1.00     3268     3824

Regression Coefficients:
              Estimate Est.Error l-95% CI u-95% CI Rhat Bulk_ESS Tail_ESS
Intercept[1]     -1.51      0.21    -1.91    -1.10 1.00     2000     3269
Intercept[2]      0.04      0.20    -0.35     0.44 1.00     1964     3401
Intercept[3]      1.79      0.21     1.39     2.20 1.00     2060     3442
z_exponent[1]    -0.42      0.18    -0.77    -0.08 1.00     2954     4791
z_exponent[2]    -0.28      0.17    -0.62     0.04 1.00     2758     3888
z_exponent[3]    -0.23      0.17    -0.58     0.12 1.00     2919     4880
z_offset[1]       0.05      0.16    -0.27     0.37 1.00     3574     5164
z_offset[2]       0.02      0.15    -0.27     0.32 1.00     3405     4991
z_offset[3]      -0.07      0.16    -0.39     0.25 1.00     3635     5480

Further Distributional Parameters:
     Estimate Est.Error l-95% CI u-95% CI Rhat Bulk_ESS Tail_ESS
disc     1.00      0.00     1.00     1.00   NA       NA       NA

Draws were sampled using sample(hmc). For each parameter, Bulk_ESS
and Tail_ESS are effective sample size measures, and Rhat is the potential
scale reduction factor on split chains (at convergence, Rhat = 1).
```

Table 13A. Output summary for: Model 8.a CSCLMM-APERIODIC-with slopes/C3

MODEL 8.b CSCLMM-APERIODIC exponent-with slopes/C3

```
Family: cumulative
  Links: mu = logit; disc = identity
Formula: rating ~ cs(z_exponent) + (1 + z_exponent || subject) + (1 | artworkn)
```

```
   Data: df_c3 (Number of observations: 2469)
  Draws: 4 chains, each with iter = 4000; warmup = 2000; thin = 1;
         total post-warmup draws = 8000
```

```
Multilevel Hyperparameters:
~artworkn (Number of levels: 113)
              Estimate Est.Error l-95% CI u-95% CI Rhat Bulk_ESS Tail_ESS
sd(Intercept)     0.64      0.06     0.52     0.77 1.00     3057     4332
~subject (Number of levels: 22)
               Estimate Est.Error l-95% CI u-95% CI Rhat Bulk_ESS Tail_ESS
sd(Intercept)      0.92      0.16     0.67     1.29 1.00     2388     4284
sd(z_exponent)     0.49      0.14     0.25     0.78 1.00     3354     3714
Regression Coefficients:
              Estimate Est.Error l-95% CI u-95% CI Rhat Bulk_ESS Tail_ESS
Intercept[1]     -1.52      0.20    -1.92    -1.11 1.00     1628     2987
Intercept[2]      0.03      0.20    -0.36     0.43 1.00     1591     2985
Intercept[3]      1.79      0.21     1.38     2.20 1.00     1610     3110
z_exponent[1]    -0.38      0.14    -0.66    -0.12 1.00     2792     3815
z_exponent[2]    -0.27      0.13    -0.53    -0.01 1.00     2666     3752
z_exponent[3]    -0.29      0.14    -0.57    -0.02 1.00     2697     3843
```

Table 14A. Output summary for: Model 8.b CSCLMM-APERIODIC exponent-with slopes/C3

MODEL 9. CSCLMM-interactions /T6

```
Family: cumulative
  Links: mu = logit; disc = identity
Formula: rating ~ cs(pw_beta1) + cs(pw_beta2) + cs(z_exponent) +
cs(pw_beta1:z_exponent) + cs(pw_beta2:z_exponent) + (1 | subject) + (1 | artworkn)
   Data: df_model (Number of observations: 2470)
  Draws: 4 chains, each with iter = 4000; warmup = 2000; thin = 1;
         total post-warmup draws = 8000
```

```
Multilevel Hyperparameters:
~artworkn (Number of levels: 113)
              Estimate Est.Error l-95% CI u-95% CI Rhat Bulk_ESS Tail_ESS
sd(Intercept)     0.64      0.06     0.52     0.77 1.00     2928     4636
~subject (Number of levels: 22)
              Estimate Est.Error l-95% CI u-95% CI Rhat Bulk_ESS Tail_ESS
sd(Intercept)     0.95      0.16     0.70     1.30 1.00     1827     3486
Regression Coefficients:
                       Estimate Est.Error l-95% CI u-95% CI Rhat Bulk_ESS Tail_ESS
Intercept[1]              -1.49      0.21    -1.92    -1.09 1.00     1367     2406
Intercept[2]               0.05      0.21    -0.37     0.46 1.00     1339     2592
Intercept[3]               1.88      0.22     1.46     2.31 1.00     1407     2786
pw_beta1[1]               -0.20      0.11    -0.41     0.01 1.00     5492     6027
pw_beta1[2]               -0.15      0.10    -0.34     0.03 1.00     4745     5357
```

```
pw_beta1[3]               -0.28      0.12    -0.52    -0.05 1.00     5540     5853
pw_beta2[1]               -0.13      0.09    -0.30     0.04 1.00     4989     5662
pw_beta2[2]               -0.18      0.08    -0.34    -0.03 1.00     4428     5628
pw_beta2[3]               -0.36      0.10    -0.55    -0.18 1.00     5548     6977
z_exponent[1]             -0.11      0.08    -0.27     0.05 1.00     3894     5300
z_exponent[2]             -0.03      0.08    -0.18     0.12 1.00     3590     4736
z_exponent[3]             -0.06      0.09    -0.24     0.12 1.00     4191     5165
pw_beta1:z_exponent[1]    -0.07      0.07    -0.21     0.06 1.00     5427     5878
pw_beta1:z_exponent[2]    -0.06      0.06    -0.18     0.06 1.00     5198     5910
pw_beta1:z_exponent[3]    -0.18      0.09    -0.37    -0.01 1.00     6491     6301
pw_beta2:z_exponent[1]    -0.04      0.07    -0.18     0.09 1.00     5752     5928
pw_beta2:z_exponent[2]     0.02      0.06    -0.10     0.15 1.00     5406     5934
pw_beta2:z_exponent[3]    -0.01      0.08    -0.17     0.15 1.00     6541     5735
```

Table 15A. Output summary for: Model 9. CSCLMM-interactions /T6

MODEL 10. CSCLMM-INTERACTIONS/C3

```
Family: cumulative
  Links: mu = logit; disc = identity
Formula: rating ~ cs(pw_beta1) + cs(pw_beta2) + cs(z_exponent) +
cs(pw_beta1:z_exponent) + cs(pw_beta2:z_exponent) + (1 | subject) + (1 | artworkn)
   Data: df_modelC3 (Number of observations: 2465)
  Draws: 4 chains, each with iter = 4000; warmup = 2000; thin = 1;
         total post-warmup draws = 8000
```

```
Multilevel Hyperparameters:
~artworkn (Number of levels: 113)
              Estimate Est.Error l-95% CI u-95% CI Rhat Bulk_ESS Tail_ESS
sd(Intercept)     0.64      0.06     0.52     0.77 1.00     2927     4459

~subject (Number of levels: 22)
              Estimate Est.Error l-95% CI u-95% CI Rhat Bulk_ESS Tail_ESS
sd(Intercept)     0.92      0.15     0.67     1.26 1.00     2221     3834

Regression Coefficients:
                       Estimate Est.Error l-95% CI u-95% CI Rhat Bulk_ESS Tail_ESS
Intercept[1]              -1.51      0.20    -1.91    -1.12 1.00     1933     3319
Intercept[2]               0.03      0.20    -0.37     0.42 1.00     1892     3060
Intercept[3]               1.71      0.20     1.32     2.10 1.00     1959     3560
pw_beta1[1]                0.04      0.07    -0.09     0.17 1.00     6034     5261
pw_beta1[2]                0.06      0.06    -0.06     0.17 1.00     5351     5405
pw_beta1[3]                0.01      0.07    -0.12     0.15 1.00     5960     5814
pw_beta2[1]                0.11      0.07    -0.02     0.25 1.00     7201     6002
pw_beta2[2]                0.06      0.06    -0.06     0.17 1.00     6143     6331
pw_beta2[3]                0.02      0.07    -0.11     0.15 1.00     6997     6390
z_exponent[1]             -0.35      0.09    -0.52    -0.18 1.00     5887     5416
```

```
z_exponent[2]              -0.28      0.08    -0.44    -0.12 1.00     5690     5503
z_exponent[3]              -0.16      0.09    -0.33     0.01 1.00     6211     5924
pw_beta1:z_exponent[1]      0.09      0.05    -0.01     0.19 1.00     7065     5611
pw_beta1:z_exponent[2]      0.03      0.05    -0.05     0.12 1.00     6703     5818
pw_beta1:z_exponent[3]     -0.11      0.06    -0.23     0.00 1.00     8748     6723
pw_beta2:z_exponent[1]     -0.06      0.06    -0.17     0.05 1.00     7823     6031
pw_beta2:z_exponent[2]      0.07      0.05    -0.03     0.16 1.00     7611     5836
pw_beta2:z_exponent[3]     -0.08      0.06    -0.19     0.04 1.00     9072     5807

Further Distributional Parameters:
     Estimate Est.Error l-95% CI u-95% CI Rhat Bulk_ESS Tail_ESS
disc     1.00      0.00     1.00     1.00   NA       NA       NA

Draws were sampled using sample(hmc). For each parameter, Bulk_ESS
and Tail_ESS are effective sample size measures, and Rhat is the potential
scale reduction factor on split chains (at convergence, Rhat = 1).
Warning messages:
1: Category specific effects for this family should be considered experimental and
may have convergence issues.
2: There were 183 divergent transitions after warmup. Increasing adapt_delta above
0.9999 may help. See
http://mc-stan.org/misc/warnings.html#divergent-transitions-after-warmup
3: Category specific effects for this family should be considered experimental and
may have convergence issues.
```

Table 16A. Output summary for: Model 10. CSCLMM-INTERACTIONS/C3

MODEL 11. CSCLMM-INTERACTIONS+SLOPES /T6

```
Family: cumulative
  Links: mu = logit; disc = identity
Formula: rating ~ cs(pw_beta1) + cs(pw_beta2) + cs(z_exponent) +
cs(pw_beta1:z_exponent) + cs(pw_beta2:z_exponent) + pw_alpha + pw_gamma + (1 |
subject) + (0 + pw_alpha | subject) + (0 + pw_beta1 | subject) + (0 + pw_beta2 |
subject) + (0 + pw_gamma | subject) + (0 + z_exponent | subject) + (1 | artworkn)
   Data: df_modelT6 (Number of observations: 2470)
  Draws: 4 chains, each with iter = 4000; warmup = 2000; thin = 1;
         total post-warmup draws = 8000
```

```
Multilevel Hyperparameters:
~artworkn (Number of levels: 113)
              Estimate Est.Error l-95% CI u-95% CI Rhat Bulk_ESS Tail_ESS
sd(Intercept)     0.67      0.06     0.56     0.80 1.00     2658     5042

~subject (Number of levels: 22)
               Estimate Est.Error l-95% CI u-95% CI Rhat Bulk_ESS Tail_ESS
sd(Intercept)      1.08      0.20     0.75     1.52 1.00     2547     3890
```

```
sd(pw_alpha)        0.61      0.16     0.34     0.96 1.00     2310     3975
sd(pw_beta1)        0.28      0.17     0.02     0.65 1.00     1639     2784
sd(pw_beta2)        0.33      0.14     0.07     0.64 1.00     1916     2077
sd(pw_gamma)        0.32      0.14     0.05     0.62 1.00     1615     1774
sd(z_exponent)      0.23      0.14     0.01     0.52 1.01     1623     2352

Regression Coefficients:
                       Estimate Est.Error l-95% CI u-95% CI Rhat Bulk_ESS Tail_ESS
Intercept[1]              -1.41      0.26    -1.91    -0.91 1.00     2076     3844
Intercept[2]               0.18      0.25    -0.31     0.68 1.00     2089     3684
Intercept[3]               2.07      0.26     1.57     2.59 1.00     2129     3630
pw_alpha                  -0.30      0.17    -0.64     0.01 1.00     3074     4640
pw_gamma                   0.16      0.11    -0.06     0.39 1.00     4673     4879
pw_beta1[1]               -0.25      0.14    -0.55     0.02 1.00     2871     3518
pw_beta1[2]               -0.21      0.13    -0.48     0.05 1.00     2642     3073
pw_beta1[3]               -0.33      0.15    -0.64    -0.05 1.00     2886     3700
pw_beta2[1]               -0.06      0.13    -0.30     0.21 1.00     2482     3189
pw_beta2[2]               -0.10      0.12    -0.33     0.15 1.00     2352     3087
pw_beta2[3]               -0.30      0.13    -0.54    -0.03 1.00     2609     2919
z_exponent[1]             -0.08      0.11    -0.31     0.14 1.00     3107     3526
z_exponent[2]              0.00      0.11    -0.22     0.21 1.00     2944     3500
z_exponent[3]             -0.02      0.12    -0.26     0.20 1.00     3088     4158
pw_beta1:z_exponent[1]    -0.12      0.09    -0.31     0.06 1.00     3342     4367
pw_beta1:z_exponent[2]    -0.10      0.09    -0.28     0.07 1.00     3222     4355
pw_beta1:z_exponent[3]    -0.22      0.11    -0.45    -0.01 1.00     3879     5077
pw_beta2:z_exponent[1]    -0.10      0.09    -0.29     0.08 1.00     3631     4940
pw_beta2:z_exponent[2]    -0.03      0.09    -0.20     0.14 1.00     3429     4253
pw_beta2:z_exponent[3]    -0.06      0.10    -0.26     0.14 1.00     4209     5311

Further Distributional Parameters:
     Estimate Est.Error l-95% CI u-95% CI Rhat Bulk_ESS Tail_ESS
disc     1.00      0.00     1.00     1.00   NA       NA       NA

Draws were sampled using sample(hmc). For each parameter, Bulk_ESS
and Tail_ESS are effective sample size measures, and Rhat is the potential
scale reduction factor on split chains (at convergence, Rhat = 1).
Warning messages:
1: Category specific effects for this family should be considered experimental and
may have convergence issues.
2: There were 10 divergent transitions after warmup. Increasing adapt_delta above
0.9999 may help. See
http://mc-stan.org/misc/warnings.html#divergent-transitions-after-warmup
3: Category specific effects for this family should be considered experimental and
may have convergence issues.
```

Table 17A. Output summary for: Model 11. CSCLMM-INTERACTIONS+SLOPES /T6

MODEL 11. CSCLMM-INTERACTIONS+SLOPES /T6

```
Family: cumulative
  Links: mu = logit; disc = identity
Formula: rating ~ cs(pw_beta1) + cs(pw_beta2) + cs(z_exponent) +
cs(pw_beta1:z_exponent) + cs(pw_beta2:z_exponent) + pw_alpha + pw_gamma + (1 |
subject) + (0 + pw_alpha | subject) + (0 + pw_beta1 | subject) + (0 + pw_beta2 |
subject) + (0 + pw_gamma | subject) + (0 + z_exponent | subject) + (1 | artworkn)
   Data: df_modelT6 (Number of observations: 2470)
  Draws: 4 chains, each with iter = 4000; warmup = 2000; thin = 1;
         total post-warmup draws = 8000
```

Multilevel Hyperparameters:

~artworkn (Number of levels: 113)

| | Estimate | Est.Error | l-95% CI | u-95% CI | Rhat | Bulk_ESS | Tail_ESS |
|---|---|---|---|---|---|---|---|
| sd(Intercept) | 0.67 | 0.06 | 0.56 | 0.80 | 1.00 | 2658 | 5042 |

~subject (Number of levels: 22)

| | Estimate | Est.Error | l-95% CI | u-95% CI | Rhat | Bulk_ESS | Tail_ESS |
|---|---|---|---|---|---|---|---|
| sd(Intercept) | 1.08 | 0.20 | 0.75 | 1.52 | 1.00 | 2547 | 3890 |
| sd(pw_alpha) | 0.61 | 0.16 | 0.34 | 0.96 | 1.00 | 2310 | 3975 |
| sd(pw_beta1) | 0.28 | 0.17 | 0.02 | 0.65 | 1.00 | 1639 | 2784 |
| sd(pw_beta2) | 0.33 | 0.14 | 0.07 | 0.64 | 1.00 | 1916 | 2077 |
| sd(pw_gamma) | 0.32 | 0.14 | 0.05 | 0.62 | 1.00 | 1615 | 1774 |
| sd(z_exponent) | 0.23 | 0.14 | 0.01 | 0.52 | 1.01 | 1623 | 2352 |

Regression Coefficients:

| | Estimate | Est.Error | l-95% CI | u-95% CI | Rhat | Bulk_ESS | Tail_ESS |
|---|---|---|---|---|---|---|---|
| Intercept[1] | -1.41 | 0.26 | -1.91 | -0.91 | 1.00 | 2076 | 3844 |
| Intercept[2] | 0.18 | 0.25 | -0.31 | 0.68 | 1.00 | 2089 | 3684 |
| Intercept[3] | 2.07 | 0.26 | 1.57 | 2.59 | 1.00 | 2129 | 3630 |
| pw_alpha | -0.30 | 0.17 | -0.64 | 0.01 | 1.00 | 3074 | 4640 |
| pw_gamma | 0.16 | 0.11 | -0.06 | 0.39 | 1.00 | 4673 | 4879 |
| pw_beta1[1] | -0.25 | 0.14 | -0.55 | 0.02 | 1.00 | 2871 | 3518 |
| pw_beta1[2] | -0.21 | 0.13 | -0.48 | 0.05 | 1.00 | 2642 | 3073 |
| pw_beta1[3] | -0.33 | 0.15 | -0.64 | -0.05 | 1.00 | 2886 | 3700 |
| pw_beta2[1] | -0.06 | 0.13 | -0.30 | 0.21 | 1.00 | 2482 | 3189 |
| pw_beta2[2] | -0.10 | 0.12 | -0.33 | 0.15 | 1.00 | 2352 | 3087 |
| pw_beta2[3] | -0.30 | 0.13 | -0.54 | -0.03 | 1.00 | 2609 | 2919 |
| z_exponent[1] | -0.08 | 0.11 | -0.31 | 0.14 | 1.00 | 3107 | 3526 |
| z_exponent[2] | 0.00 | 0.11 | -0.22 | 0.21 | 1.00 | 2944 | 3500 |
| z_exponent[3] | -0.02 | 0.12 | -0.26 | 0.20 | 1.00 | 3088 | 4158 |
| pw_beta1:z_exponent[1] | -0.12 | 0.09 | -0.31 | 0.06 | 1.00 | 3342 | 4367 |
| pw_beta1:z_exponent[2] | -0.10 | 0.09 | -0.28 | 0.07 | 1.00 | 3222 | 4355 |
| pw_beta1:z_exponent[3] | -0.22 | 0.11 | -0.45 | -0.01 | 1.00 | 3879 | 5077 |
| pw_beta2:z_exponent[1] | -0.10 | 0.09 | -0.29 | 0.08 | 1.00 | 3631 | 4940 |

```
pw_beta2:z_exponent[2]    -0.03      0.09    -0.20     0.14 1.00     3429     4253
pw_beta2:z_exponent[3]    -0.06      0.10    -0.26     0.14 1.00     4209     5311

Further Distributional Parameters:
     Estimate Est.Error l-95% CI u-95% CI Rhat Bulk_ESS Tail_ESS
disc     1.00      0.00     1.00     1.00   NA       NA       NA

Draws were sampled using sample(hmc). For each parameter, Bulk_ESS
and Tail_ESS are effective sample size measures, and Rhat is the potential
scale reduction factor on split chains (at convergence, Rhat = 1).
Warning messages:
1: Category specific effects for this family should be considered experimental and
may have convergence issues.
2: There were 10 divergent transitions after warmup. Increasing adapt_delta above
0.9999 may help. See
http://mc-stan.org/misc/warnings.html#divergent-transitions-after-warmup
3: Category specific effects for this family should be considered experimental and
may have convergence issues.
```

Table 18A. Output summary for sensitivity to trial position T6 model.
